# Sexually divergent development of depression-related brain networks during healthy human adolescence

**DOI:** 10.1101/2020.07.06.184473

**Authors:** L. Dorfschmidt, R.A.I. Bethlehem, J. Seidlitz, F. Váša, S.R. White, R. Romero-García, M.G. Kitzbichler, A. Aruldass, S.E. Morgan, I.M. Goodyer, P. Fonagy, P.B. Jones, R.J. Dolan, the NSPN consortium, N.A. Harrison, P.E. Vértes, E.T. Bullmore

## Abstract

We hypothesized that there are sexual differences in human brain network development underlying the female > male divergence in adolescent depression. We tested for sex differences in parameters of brain network development (accelerated longitudinal fMRI, N=298 healthy adolescents, each scanned 1 to 3 times). Sexually divergent development of functional connectivity was located in default mode network (DMN), limbic cortex, and subcortical nuclei. Females had a more “disruptive” pattern of development, where weak functional connectivity at age 14 became stronger during adolescence. This fMRI-derived map of divergent adolescent development was co-located with (i) a map of functional dysconnectivity associated with adult major depressive disorder (MDD); and (ii) an adult brain gene expression pattern enriched for genes on the X chromosome, neurodevelopmental genes, and risk genes for MDD. Sexual divergence in disruptive development of DMN, limbic and subcortical functional networks is potentially relevant to the increased risk of depression in adolescent females.

## Introduction

Adolescence is a period of critical development of the brain, characterized by changes in both structure [1, 2, 3, 4] and function [5, 6], that coincide with changes in cognition and behaviour. It is also a time of increasing incidence of many psychiatric disorders, including depression, which occurs more frequently in females than males [7, 8]. Small sex differences in mood have been reported from the age of 11, and by the age of 15 females are about twice as likely to be depressed as males [7, 8, 9]. Recent work has supported the idea that sexually divergent risk for mood disorders could be related to sex-differences in adolescent brain network development [10].

Functional brain networks derived from resting state functional magnetic resonance imaging (rs-fMRI) can be used to study complex network organization in the brain. Each node of these networks is an anatomical region and each edge weight is an estimator of association, so-called functional connectivity, typically the correlation or coherence between the two fMRI signals simultaneously measured for each possible pair of nodes in the network [11, 12].

The brain is plastic and undergoes maturational changes throughout life. Primary sensory and motor areas mature most rapidly during childhood, while association areas undergo their most profound changes during late adolescence [6, 13, 14]. Previous resting-state fMRI studies have reported a shift from local to distributed networks [15], and an increase in the strength of long-range connections [16, 17], in the course of adolescence.

However, it has since been noted that in-scanner head motion may have confounded many of the effects previously attributed to age, particularly in younger participants [18, 19, 20]. Developmental imaging studies have therefore employed different strategies to address these concerns, e.g., by restricting analysis to motion-uncontaminated sub-samples of acquired data with no detectable head motion [6], or by regressing each nodal fMRI signal on the global average fMRI signal, a.k.a. global signal regression (GSR) [21]. Issues concerning optimal head motion correction for pre-processing fMRI data remain controversial [22, 23, 24].

It is not yet clear how functional connectivity differs between males and females, either during adolescence or adulthood. One widely reported sex difference is increased functional connectivity of the default mode network (DMN) in females [25, 26, 27, 28, 29]. Female-increased (or female > male) connectivity has also been reported in the subcortex and limbic areas (cingulate gyrus, amygdala, hippocampus) [30]; whereas male > female connectivity has been reported for sensori-motor areas [30, 25, 28]. However, these effects are not consistently found across studies [27, 26, 31]. Importantly, most research on sex differences has focused on pre-selected regions, often including the amygdala [32, 33], with few studies having investigated sex differences comprehensively over all brain regions [25, 28, 34, 35, 36]. Most prior fMRI studies of brain development have focused on estimating “average” effects of age across both sexes, e.g. by including sex as a covariate in the statistical model for estimation of developmental parameters (Supplementary Table 4) and few prior studies have reported age-by-sex interactions or the conditioning of developmental parameters by sex [35, 30].

We start from the hypothesis that the sexually divergent risk trajectory for depression, with higher depressive symptom scores for adolescent females than males [7, 8], could be the psychological or clinical representation of underlying sex differences in adolescent brain network development [34, 27, 28]. To investigate this overarching hypothesis experimentally, we first identified sexually divergent systems of healthy adolescent brain development, and then tested the anatomical co-location of sexually divergent fMRI systems with prior maps of human brain gene expression and depression-related fMRI phenotypes.

Using fMRI data from a previously published [6] accelerated longitudinal study (N=298; age range 14-26 years; 51% female; Table 1), stratified by age and balanced for sex per age stratum [37], we estimated the effects of sex on three parameters of adolescent development of resting-state functional connectivity: (1) baseline connectivity at age 14 *FC*_14_ and (2) the adolescent rate of change *FC*_14-26_ estimated at the node and edgewise-level; and the maturational index for each node (3), that is the signed correlation coefficient between *FC*_14_ and *FC*_14-26_ across all edges connecting a given node to the rest of the network. The sign of the maturational index (positive or negative) has been reported to reflect two distinct modes of adolescent development of brain functional connectivity [6]: so-called, conservative and disruptive modes. A conservative node is defined by a positive MI -indicating that it is highly connected or “hub-like” at baseline (14 y) and becomes even more strongly connected over the course of adolescence (14-26 y). Theoretically, conservative nodes could also be weakly connected at baseline and become even more weakly connected during adolescence; however, empirically, we found that this is not the case (SI Fig. S14). A disruptive node is defined by a negative MI -indicating either that it is weakly connected at age 14 but becomes more strongly connected or hub-like during adolescence; or that is a strongly connected node at 14 y but becomes more weakly connected or less hub-like during adolescence. The disruptive developmental profile of weak-getting-stronger during adolescence hypothetically represents a “re-wiring in the functional connectome, which could be relevant to the acquisition of social cognitive and other skills [6]. It has also been argued that brain networks that are most developmentally active during adolescence are most likely to contribute to the coincidentally increased risk of mental health symptoms, i.e., “moving parts get broken” [10]. For these reasons, our analysis focused particularly on sexual differences in weak-getting-stronger disruption in cortico-subcortical networks; results for strong-getting-stronger conservative development are summarised in Supplementary Fig. 16.

**Table 1:**
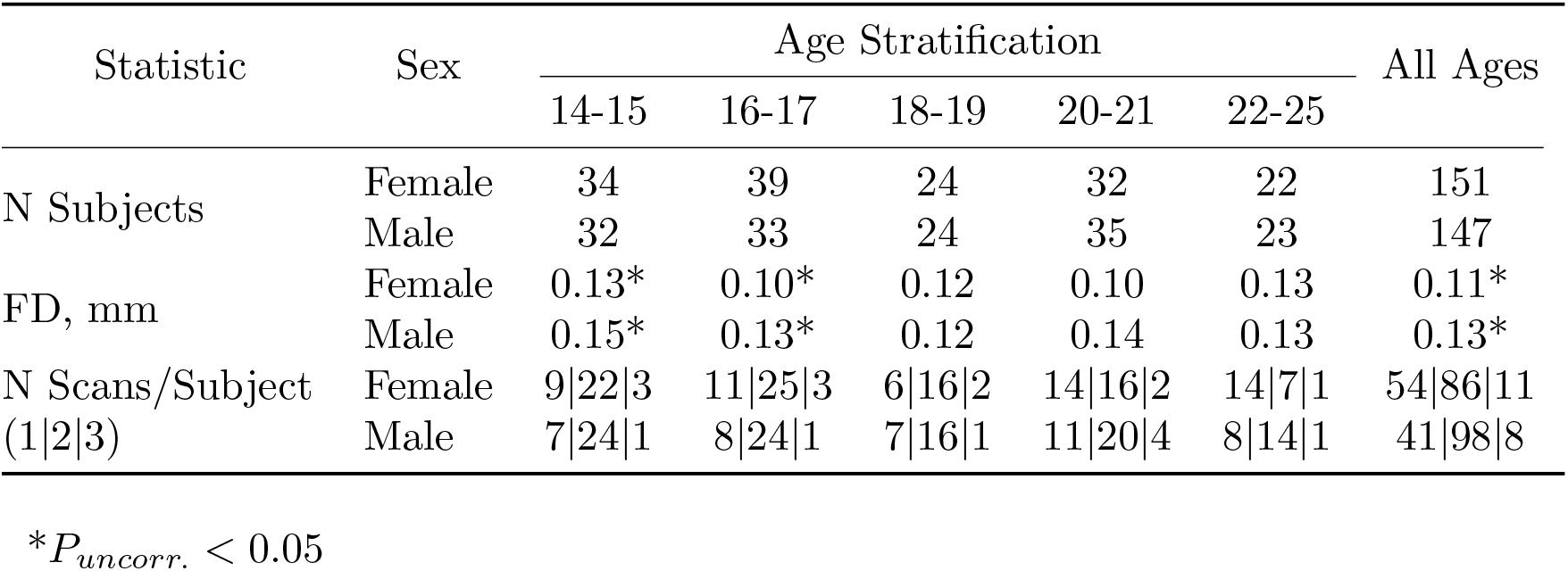
Adolescent developmental MRI sample. Total N = 298 healthy young people participated in an accelerated longitudinal MRI study, with recruitment balanced for sex in each of five age-defined strata, and each subject scanned between 1 and 3 times (with follow up scans taking place approximately 6 and 18 months after baseline). FD = framewise displacement, a measure of head movement in mm, was significantly greater in males compared to females on average over all ages, and in the youngest two age strata specifically *(P* < 0.05, uncorrected; Supplementary Fig. 5).

We found that females had predominantly more disruptive development of functional connectivity in a default mode cortical, limbic and subcortical network. We then tested the hypotheses that this developmentally divergent brain system was co-located with expression of a weighted function of the whole genome enriched for X chromosome genes, genes related to perinatal and post-natal brain development, and genes related to major depressive disorder (MDD). We also tested the hypothesis that the sexually divergent system was co-located with an anatomical map of depression-related differences in functional connectivity from an independent casecontrol fMRI study of MDD [38].

There was a significant sex difference in head motion (framewise displacement, FD) during fMRI scanning, so we pre-processed the data to mitigate the potentially confounding effects of head motion and subsequently demonstrated the robustness of the key results to alternative fMRI pre-processing strategies (Supplementary Fig. 2-3; Supplementary Text *“Sensitivity Analysis”*).

## Results

### Analysable sample and head motion

A total of 36 scans were excluded by quality control criteria including high in-scanner motion (mean FD>0.3 mm or maximum FD>1.3 mm), co-registration errors, or extensive dropout. The analyzable sample thus consisted of 520 scans from 298 participants (151 females; Table 1). Males had significantly more head movement than females in the youngest two age strata (P<0.05, uncorrected) and on average over all ages (Table 1, Supplementary Fig. 2-3).

After pre-processing for within-subject correction of head motion effects on individual fMRI time series, functional connectivity was positively correlated with individual differences in mean FD, and this effect scaled with distance between the nodes (Supplementary Fig. 3A). We therefore also corrected for between-subject differences in head motion by regressing each inter-regional correlation on mean FD across all participants. This removed the relationship between connectivity and FD, as well as the distance-dependence in this relationship [18, 39] (Supplementary Fig. 3D). To assess the robustness of key results to this two-step process for head motion correction, we conducted three sensitivity analyses (Supplementary Text *“Sensitivity Analysis”*): (i) Sex-specific motion correction -FC matrices were regressed on FD separately for males and females (Supplementary Fig. 28-31); (ii) GSR correction -the fMRI time series at each node were regressed on the global fMRI signal per participant (Supplementary Fig. 25-27); (iii) Motion-matched sub-sample analysis -we used a subset of data (N=314), comprising equal numbers of males and females, for which there was no statistical difference in FD (Supplementary Fig. 21-24). Given the male > female sex difference in intra-cranial volume, related to the sex difference in body size, we ran a fourth sensitivity analysis: (iv) Intra-cranial volume correction -we regressed all fMRI metrics on intra-cranial volume (ICV) estimated from structural MRI data on the same sample (Supplementary Fig. 34-35). There was a significant correlation between the developmental parameters (*FC*_14_, *FC*_14-26_, and *MI*) estimated by each of these alternative analyses, and the same parameters estimated by our principal method (Supplementary Fig. 30-34). The key finding of divergent adolescent development of functional connectivity between DMN, limbic and subcortical regions, as reported below, was conserved in all cases.

### Age and sex effects on functional connectivity

We modeled age and sex effects on global functional connectivity of each participant, estimated as mean weighted degree, using linear mixed effects models (LMEs). Functional connectivity increased with age (*t*(219)=2.3, *P*<0.05) and males had higher global mean weighted degree than females (*t*(296)=5.5, *P*<0.0001) (Fig. 1A; Supplementary Table 2).

**Figure 1:**
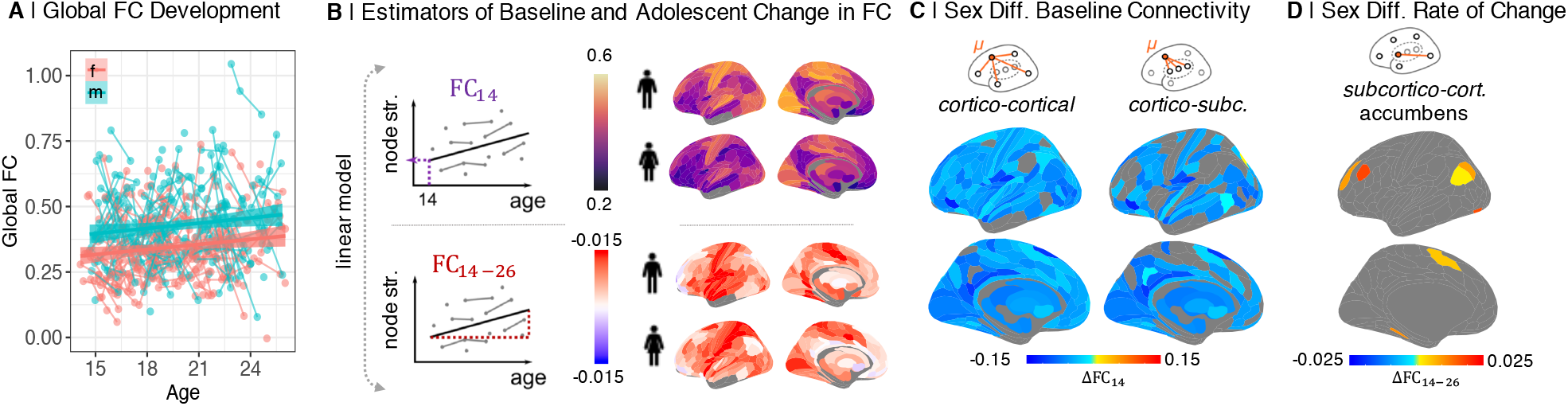
Sex differences in functional connectivity at age 14 *(FC*_14_) and adolescent rate of change of connectivity *(FC*_14-26_) per year: (A) Global functional connectivity (FC) strength increased with age (*t*(219)=2.3, *P*<0.05) and was higher in males (*t*(296)=5.5, *P*<0.0001). (B) To estimate two parameters of development at each regional node, we fit a linear model to the relationship between age and weighted degree (nodal strength of connectivity to the rest of the network) for males and females separately. The two model parameters are the intercept, or “baseline” connectivity at age 14 *(FC* _14_), and the linear rate of change in connectivity during adolescence *(FC*_14-26_). (C) We found 321/330 regions had significantly increased cortico-cortical connectivity, and 230/330 regions had increased cortico-subcortical connectivity *(P*_*FDR*_<0.05), at baseline, *FC*_14_, in males. (D) *FC*_14-26_ was only significantly different between sexes, decreased in females, in 27/330 subcortico-cortical connections of the nucleus accumbens.

### Sex differences in parameters of adolescent development

Regional functional connectivity was estimated between and within cortical and subcortical subsets of nodes by averaging the relevant parts of the connectivity matrix (Supplementary Fig. 7). To model development of functional connectivity during adolescence, we focused on three parameters: regional baseline connectivity at age 14, *FC*_14_; regional linear change in connectivity between 14-26 years, *FC*_*14*-*26*_ (Fig. 1B); and the signed Spearman correlation of these two parameters, termed maturational index (— 1 < *MI* < +1) (Fig. 2A; [6]). Previous work on this sample has reported developmental change (controlling for sex) in terms of these parameters estimated at each regional node of a whole brain fMRI network [6]. Here we estimated each of these parameters for males and females separately, and the between-sex difference for each parameter, e.g., Δ*MI = MI*_*female*_*-MI*_*male*_. We tested the significance of the between-sex difference in each parameter at each regional node using parametric tests (Supplementary Text *“Analysis of Effects on Parameters of Adolescent Brain Development”*).

**Figure 2:**
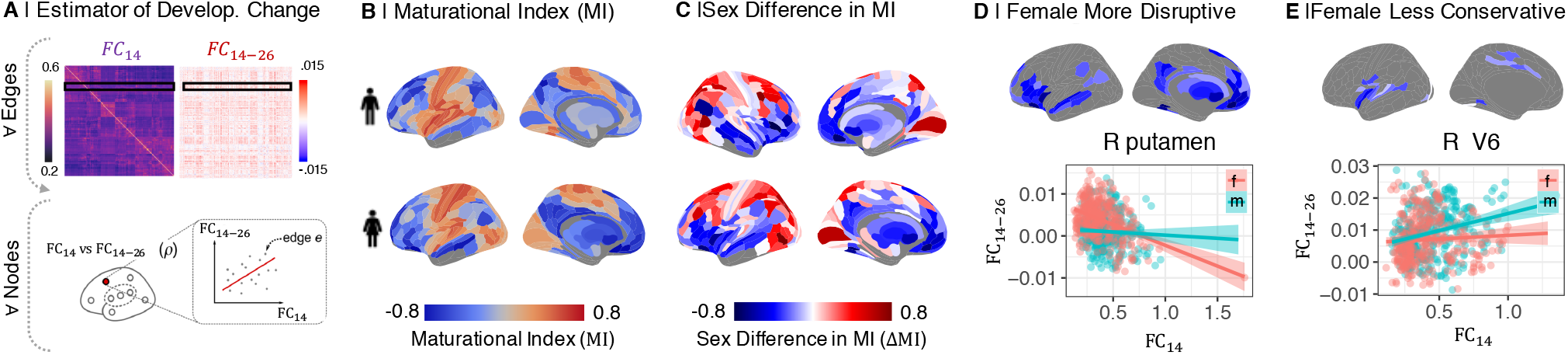
Sex differences in maturational index (MI): (A) The maturational index (MI) was estimated as the correlation between edgewise baseline connectivity at age 14 *(FC*_14_), and the adolescent rate of change in connectivity per year *(FC*_14-26_), at each regional node. (B) MI maps for males and females separately. MI was generally negative (blue) in frontal and association cortical areas, and positive (orange) in primary motor and sensory cortices. (C) The sex difference in MI, Δ*MI = MI*_*female*_ *— MI*_*male*_, was significant in 230/346 regional nodes *(P*_*FDR*_ = 0.05). Δ*MI* was significantly negative in the ventral and medial prefrontal gyrus, ventrolateral prefrontal cortex, anterior and posterior cingulate gyrus, medial temporal gyrus, and subcortical nuclei (Supplementary Table 3), indicating sex differences in adolescent development of connectivity of these regions. More specifically, negative Δ*MI* defined a set of brain regions where adolescent development was either more disruptive (weak connections at 14 years became stronger during adolescence, or strong connections became weaker) or less conservative (strong connections at 14 years became stronger or weak connections became weaker during adolescence) in females compared to males. (D) Map of brain regions where development was more disruptive in females. As exemplified by right putamen, functional connections of disruptively developing nodes that were strong at 14 years (high *FC*_14_, x-axis) became weaker over the period 14-26 years *(FC*_14-26_ < 0, y-axis), and edges that were weakly connected at 14 y became stronger over the course of adolescence, especially in females. (E) Map of brain regions where development was less conservative in females. As exemplified by right visual area V6, connections that were strong at baseline become stronger over the period 14-26 years, especially in males.

Qualitatively, *baseline connectivity at age 14* was apparently greater in primary sensorimotor cortex than in association cortex for both sexes (Fig. 1B, Supplementary Fig. 8-9). As predicted by the sex difference in global functional connectivity at all ages (Fig. 1A), males had significantly stronger baseline connectivity than females at 14 years, i.e., Δ*FC*_14_ = *FC*_14,*female*_ — *FC*_14,*male*_ > 0, in cortico-cortical and cortico-subcortical connections (Fig. 1C).

The pattern of *adolescent rate of change in connectivity* was strongly positive in sensorimotor cortex, and appeared less positive or slightly negative in association cortical and limbic areas, for both sexes. There were was no significant sex difference, i.e., Δ*FC*_*14*-*26*_ = 0, for cortico-cortical or cortico-subcortical connectivity; but a subset of 27 subcortico-cortical connections, involving the nucleus accumbens, had significantly more negative rates of change in females compared to males (Fig. 1D, Supplementary Fig. 10-11; *P*_*FDR*_ < 0.05).

*Maturational index* was positive in sensorimotor cortex, and negative in association cortex and subcortical areas, in both sexes separately (Fig. 2B; Supplementary Fig. 12), as previously reported for both sexes on average [6]. However, there were many areas of significant sex difference in MI *(P*_*FDR*_< 0.05; Supplementary Fig. 13). Females had more negative MI than males in 107 regions (Fig. 2B; for a full list see Supplementary Table 3; effect sizes are shown in Supplementary Fig. 13). In 84 of these regions, exemplified by the right putamen (Fig. 2D), there was more disruptive development in females, i.e., weak connections at 14 years became stronger during adolescence, or strong connections became weaker, in females compared to males. In 23 regions, exemplified by right visual area V6 (Fig. 2D), there was less conservative development in females, i.e., strong connections at 14 years became stronger during adolescence in males compared to females (Supplementary Fig. 15). Thus the brain system defined by regions with a negative Δ*MI* is predominantly characterized by a weak-getting-stronger profile of developmental change in functional connectivity that was greater in females than males.

The unthresholded map of Δ*MI* was co-registered with a prior map of cortical cytoarchitectonic classes (Fig. 3B) and a prior map of resting state networks [40] from an independent component analysis of adult fMRI data (Fig. 3C). Regions of negative Δ*MI* were concentrated in secondary sensory, limbic, and insular classes of cortex, and in subcortical structures, defined anatomically (Fig. 3B); and in default mode, limbic, ventral attentional and subcortical systems defined functionally (Fig. 3C).

**Figure 3:**
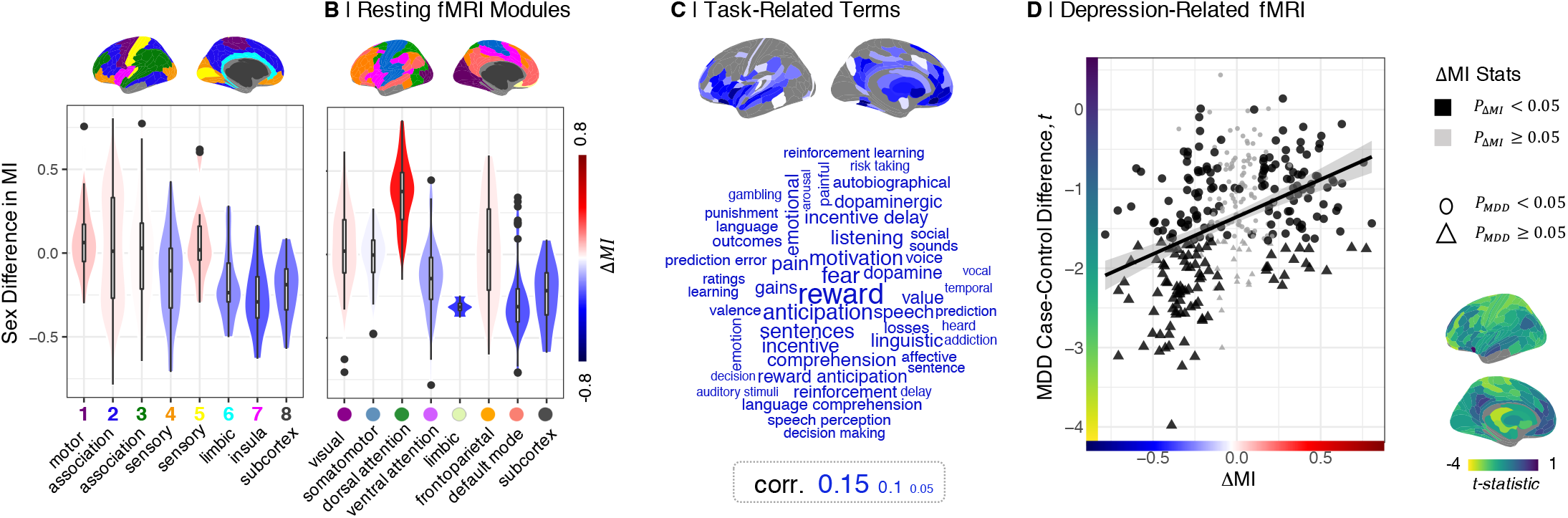
Sex difference in maturational index in psychological and psychiatric context: (A) Δ*MI* was most negative in secondary sensory, limbic, and insula cortex, and subcortical structures defined by cell type, (B) as well as functionally defined default mode network (DMN), ventral attention network, limbic, and subcortical structures. (C) Wordcloud of Neurosynth meta-analytical cognitive terms scaled according to their strength of association with the disruptively developing brain regions (cortical map of Δ*MI* < 0). (D) Top, identically parcellated brain map of MDD case-control differences in weighted degree, *t* statistics. Bottom, Δ*MI* (x-axis) was positively correlated with case-control *t*-statistics (y-axis; *ρ* = 0.4, *P* < 0.001). Regions with sexually divergent disruptive development in adolescence (negative Δ*MI*) had reduced degree of connectivity in adult MDD cases.

Automated meta-analytic referencing of the unthresholded map of negative Δ*MI* was conducted using the Neurosynth database of task-related fMRI activation coordinates [41]. This indicated that regions with more disruptive (or less conservative) development in females were typically activated by tasks related to reward processing, emotion, motivation, incentive delay, and dopamine (Fig. 3C).

### Gene transcriptional enrichment of sexually divergent developing brain systems

To investigate the relationships between gene transcriptional profiles and sexually divergent adolescent brain development, we used partial least squares (PLS) regression to find the weighted gene expression pattern that was most closely co-located with the Δ*MI* map (Fig. 4A; [5, 13, 42]). Whole genome transcripts were estimated for the average of each of 180 bilaterally homologous cortical regions using adult post-mortem data (N=6) provided by the Allen Human Brain Atlas [43].

**Figure 4:**
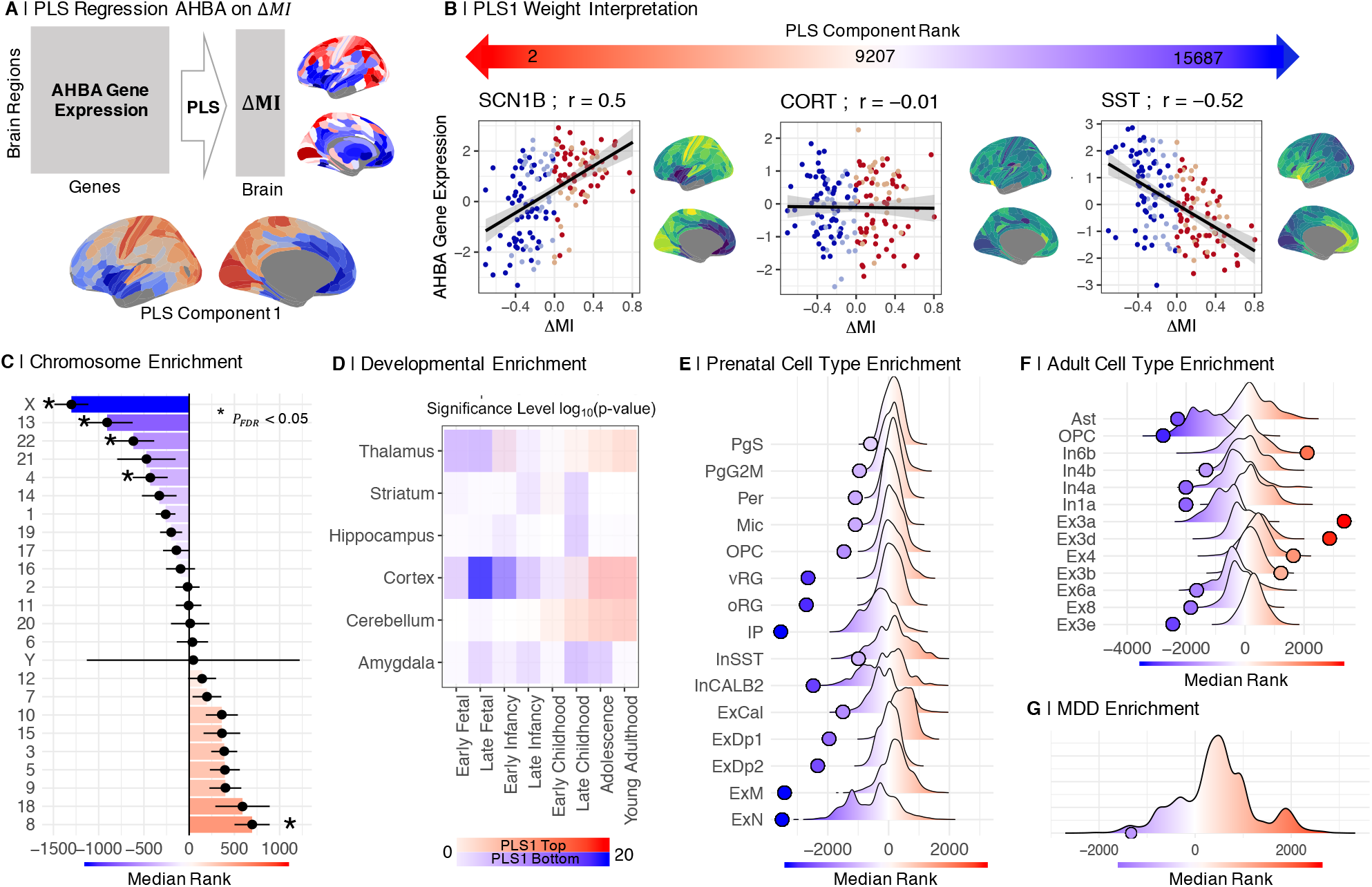
Sexually divergent disruptive brain systems are co-located with brain tissue transcripts enriched for X chromosome, neurodevelopmental, and MDD risk genes: (A) We used partial least squares (PLS) regression to map the Allen Human Brain Atlas (AHBA) gene expression data [43] onto the Δ*MI* map. (B) Relationship of Δ*MI* to expression of exemplary single genes: Sodium Voltage-Gated Channel Beta Subunit 1 *(SCN1B*), a positively weighted gene close to the top of the ranked list of PLS1 weights; cortistatin *(CORT*), a near-zero weighted gene in the middle of the list; and somatostatin *(SST*), a negatively weighted gene close to the bottom. Negatively weighted genes were more strongly expressed in regions of negative Δ*MI*, that is predominantly female > male disruptive regions ; whereas positively weighted genes were more strongly expressed in regions with female > male conservative development indicated by positive Δ*MI*. (C) Enrichment analysis for chromosomal genes. Plot of the median rank of genes from each chromosome on PLS1, with standard deviations. (D) Enrichment analysis for neurodevelopmental genes. Negatively weighted genes (blue) were enriched for genes typically expressed in cortex during late fetal and early post-natal development and for genes expressed in the amygdala, hippocampus and striatum during late childhood and adolescence. Positively weighted genes (red) were enriched for genes typically expressed in cortex and cerebellum during adolescence and early adult life. (E) Enrichment analysis for prenatal cell type-specific genes. Negatively weighted genes (blue) were significantly enriched for genes expressed by prenatal radial glia (vRG, oRG), microglia (Mic), oligodendrocyte progenitor cells (OPG), and excitatory neurons. (F) Enrichment analysis for adult cell typespecific genes. Negatively weighted genes were significantly enriched for genes expressed by adult astrocytes, OPC, and excitatory neurons. (G) Enrichment analysis for MDD-related genes. Negatively weighted genes were significantly enriched for genes associated with major depressive disorder by an independent genome wide association study [49].

The first PLS component (PLS1; Fig. 4A) explained 34.6% of the variance in Δ*MI*, significantly more than expected by chance (*P*_*perm*_ <0.05). The PLS1 gene expression weights were positively correlated with Δ*MI*, thus negatively weighted genes were over-expressed in regions with negative Δ*MI*, and positively weighted genes were under-expressed in regions of negative Δ*MI* (Fig. 4B).

To test the hypothesis that sex chromosomal gene expression was related to the sexual differences in adolescent brain development, we assessed chromosomal enrichment of the genes on PLS1. In particular, we hypothesised that gene expression patterns related to sex differences in adolescent brain development might be enriched for X chromosomal genes because the X chromosome is diploid in females (XX) and haploid in males (XY) and previous studies have shown effects of sex chromosome aneuploidies associated with neurodevelopmental disorders [44]. We found that the most negatively weighted genes, which were highly expressed in disruptively developing regions, were most strongly enriched for X chromosome genes (*P*<0.001; Fig. 4C, Supplementary Table 5; Supplementary Fig. 17).

Regional differences in cortical gene expression have been attributed to different proportions of functionally specialised neuronal, glial and other cell types in different cortical areas [45]. We therefore used the Cell Specific Enrichment Analysis tool (CSEA) tool [46] to assess cell type enrichment of the most positively and negatively weighted genes on PLS1. We found that negatively weighted genes (Z<-2.58) were enriched for genes with cortical expression in late fetal and early postnatal life, and for genes with amygdala, hippocampal and striatal expression in late childhood and adolescence (Fig. 4D). In contrast, positively weighted genes (Z>2.58) were enriched for genes with cortical, cerebellar and thalamic expression during adolescence and young adulthood (Fig. 4D). This result indicates that the negatively weighted genes, most strongly co-expressed with the disruptively developing cortico-subcortical system, were specialised for perinatal and adolescent phases of cortical and subcortical development, respectively.

We further explored developmental aspects of the sexually divergent system by testing for enrichment by genes specific to pre-natal and post-natal cell types [47, 48]. We found that genes which were over-expressed in disruptively developing brain regions were enriched for pre-natal cell types [48], including oligodendroglial precursor cells (OPC), microglia, astrocyte progenitor radial cells, inhibitory and excitatory cortical neurons (Fig. 4E), as well as for multiple adult glial and neuronal cell classes (Fig. 4F, Supplementary Fig. 18).

### Sexually divergent brain development and depression

Extending the enrichment analysis to consider depression-related genes, we found that the list of genes strongly co-expressed with sexually divergent disruptive brain systems was significantly enriched for risk genes for MDD from an epigenetically-informed, large prior GWAS study [49]. Over 80% of risk variants identified by genome wide association studies (GWAS) are found in the non-coding genome, which makes the interpretation of underlying biological mechanisms challenging. Non-coding single nucleotide polymorphisms (SNPs) can regulate distal genes via long-range regulatory interactions, since the 3D structure of the genome allows for distal enhancers to be brought into contact with promoters far downstream. Therefore, we used a gene list which mapped SNP hits from one of the largest available MDD GWAS studies [50] to functionally relevant genes using epigenetic (Hi-C) data, to guide the interpretation of hits in non-coding loci [51, 52]. Our enrichment results showed that the MDD risk genes were negatively weighted and ranked towards the bottom of the PLS1 list, indicating that they were more highly-expressed in brain regions with disruptive development, indexed by negative Δ*MI*.

To assess the anatomical correspondence between the sexually divergent disruptive brain system, and mood disorder-related changes in fMRI connectivity, we used resting state fMRI data from a prior case-control study of adult MDD cases (N=50) and healthy controls (N=46); see (Supplementary Table 4). The parcellated, unthresholded map of MDD case-control differences in weighted degree (comprising 346 regional *t* statistics), was significantly co-located with the identically parcellated, unthresholded map of Δ*MI (ρ* = 0.4, *P*<0.001; Fig. 3D). Brain regions with sexually divergent development in adolescence (negative Δ*MI*) had reduced degree of functional connectivity in MDD cases compared to controls.

## Discussion

In this accelerated longitudinal fMRI study of healthy young people, we have identified human brain systems that demonstrated a significantly different pattern of adolescent development in females compared to males. We found sex differences in several aspects of functional connectivity (FC): females had lower global mean FC across all ages, and reduced nodal strength of connectivity in most regional nodes at 14 years *FC*_14_. However, there were more anatomically specific sex differences in two developmentally sensitive parameters: the rate of change in FC during adolescence, *FC*_14-*26*_, was significantly reduced in females for connections between one cortical nucleus (nucleus accumbens) and 27 cortical structures; and the maturational index, *MI*, a coefficient of the linear relationship between edgewise *FC*_14_ and *FC*_*14*-*26*_ at each node, was significantly more negative in females for 107 cortical areas concentrated in the default mode network (DMN), ventral attentional and limbic networks, as well as subcortical nuclei.

The first explanatory hypothesis we considered was that the sex differences in developmental fMRI parameters were driven by sex differences in head motion during scanning. Males, especially younger males, had more head movement than females. We addressed this potential confound by a two-stage pre-processing pipeline which statistically corrected each participant’s functional connectome for between-subject differences in head motion, indexed by framewise displacement (FD). These pre-processed data passed standard quality control criteria for movement-related effects on functional connectivity. Moreover, we repeated the entire analysis for male and female data separately, for a “motion-matched” subset of the data in which there was no significant sex difference in FD, and for all data after global signal regression. In all four sensitivity analyses, the key results of our principal analysis were qualitatively and quantitatively conserved. We therefore consider that head motion can be discounted as a sufficient explanation for sex differences in these developmental parameters.

An alternative explanation is that sex differences in *FC*_*14*-*26*_ and *MI* reflect divergent development of specific cortico-subcortical circuits. In particular, females had a predominately more disruptive pattern of adolescent development, indexed by negative Δ*MI*, whereby functional connections that were weak at 14 years became stronger, and connections that were strong became weaker, over the course of adolescence. We hypothesized this as relating to sex differences in an underlying process of reconfiguration or remodelling of cortico-subcortical connectivity at a synaptic or neuronal scale.

To assess the plausibility of this biological interpretation, we used pre-existing data on human brain gene expression, and the dimension-reducing multivariate method of partial least squares (PLS), to identify the set of genes that were most over-or under-expressed in brain regions corresponding to the divergent system defined by developmental fMRI. Enrichment analysis demonstrated that the genes that were most strongly expressed in brain regions with more disruptive (or less conservative) development in females included significantly more X chromosome genes than expected by chance. The same set of genes was also significantly enriched for genes that are known *a priori* to be expressed in cortical areas during early (perinatal) development and in subcortical structures, such as amygdala, during adolescent development.

Sexual differentiation of the brain has been proposed to occur in two stages: an initial “organizational” stage before and immediately after birth, and a later “activational” stage during adolescence [53]. It has long been argued that these events are driven by gonadal hormones. However, more recent work suggests a complex interplay of sex chromosomes and their downstream products leading to sexual differentiation of brain cells [54, 55, 56]. The results of our enrichment analysis suggest a co-location of the sexually divergent fMRI derived map of female more disruptive brain development during adolescence and early life gene differentiation. In short, sexual differences in adolescent development of cortico-subcortical connectivity could be driven by developmentally phased programs of gene expression in the brain.

Finally, we assessed the pathogenic relevance of this sexually divergent developing brain system to the greater risk of depression in adolescent females. Macroscopically, the DMN and subcortical structures that had predominantly more disruptive development in females, e.g., ventral medial prefrontal cortex, medial temporal gyrus, anterior and posterior cingulate cortex, have previously been implicated as substrates of depressive disorder [57, 58]. This anatomical convergence was quantified by the significant spatial correlation between the whole brain map of sex differences in MI and an independent map of MDD case-control differences in nodal degree of functional connectivity. At a microscopic scale, the (PLS1) list of genes transcriptionally co-located with the divergentally developing fMRI system was enriched for risk genes from prior genome-wide association studies of MDD, as well as genes specifically expressed by adult neuronal and glial cells that have been linked to neuroimaging phenotypes of depression ([59]; Supplementary Fig. 19-20). Psychologically, cognitive functions that typically activated within the divergent system, based on meta-analysis of a large prior database of task-related fMRI studies, included reward-related processes that are fundamental to the core depressive symptom of anhedonia.

### Methodological issues

This analysis has focused on group differences between males and females. Previous work has found a substantial amount of distributional overlap between the sexes in the distributions of multiple or all brain measures [60, 61]. The metrics analysed here (*FC*_14_, *FC*_*14*-*26*_ and Δ*MI*) are group level parameters, thus all reported sex differences are reflective of a group mean difference, estimated from functional connectivity distributions that substantially overlap between the sexes. On this basis, we are not arguing that female and male brains are distinctly dimorphic [62].

It is a strength of the study that our analysis of a sexually divergent developing brain system is based on a large longitudinal fMRI dataset with approximately equal numbers of males and females in each stratum of the adolescent age range. Head motion is known to be a potentially problematic confound in developmental fMRI [18, 19, 20]. We have mitigated its impact on these data by a two-step pre-processing pipeline that satisfied prior quality control criteria; and we have demonstrated the robustness of our key results to three alternative motion correction strategies, including GSR [23] (Supplementary Fig. 21-33). Limitations of the study include our reliance on gene expression maps from post mortem examination of six adult, mostly male, brains. Biological validation of this divergent fMRI system would be more directly informed by sex-specific human brain maps of whole genome transcription in adolescence; but to the best of our knowledge such data are not currently available. It will also be important in future to test the hypothesis that an anatomically homologous cortico-subcortical system has divergent adolescent development in animal models that allow more precise but invasive analysis of the cellular and molecular substrates of fMRI phenotypes than is possible in humans. We have shown that this normative developmental dimorphism is anatomically, genetically and psychologically relevant to depression. However, further studies will be needed to test the hypothesis that the emergence of depressive symptoms in adolescent females is directly related to the sexually divergent disruptive development of DMN, limbic and subcortical connectivity that we have identified in this cohort of healthy participants.

### Conclusion

We conclude that there is sexual divergence in adolescent development of functional connectivity between nodes of a human cortico-subcortical system that is likely anatomically, genetically and psychologically relevant to depression.

## Methods

### Study sample

A total of 520 analysable fMRI scans were available for N=298 healthy participants, aged 14 to 26 years, each scanned one to three times as part of an accelerated longitudinal study of adolescent brain development (Neuroscience in Psychiatry Network, NSPN; [37, 13, 6]; Supplementary Text *“Data”*). Participants self-identified their sex as either male or female. There were approximately equal numbers of males and females in each of five age-defined strata at baseline (Table 1). All participants aged 16 years or older gave informed consent; participants younger than 16 gave informed assent and consent was provided by their parent or guardian. The study was ethically approved by the National Research Ethics Service and conducted in accordance with UK National Health Service research governance standards.

### fMRI data acquisition

Functional MRI data were acquired at three sites, on three identical 3T Siemens MRI scanners (Magnetom TIM Trio, VB17 software version), with a standard 32-channel radio-frequency (RF) receive head coil and RF body coil for transmission using a multi-echo echo-planar (ME-EPI) imaging sequence [63] with the following scanning parameters: repetition time (TR): 2.42s; GRAPPA with acceleration factor 2; Flip Angle: 90; matrix size: 64×64×34; FOV: 240×240mm; in-plane resolution: 3.75×3.75 mm; slice thickness: 3.75 mm with 10% gap, sequential slice acquisition, 34 oblique slices; bandwidth 2368 Hz/pixel; echo times (TE) 13, 30.55 and 48.1 ms.

### fMRI data pre-processing

Functional MRI data were preprocessed using multi-echo independent component analysis (ME-ICA;[64, 65]) which identifies and removes sources of variance in the times series that do not scale linearly with TE and are therefore not representative of the BOLD signal. Ventricular time series, representing variance in cerebrospinal fluid (CSF), were regressed from parenchymal time series using AFNI [66]. Scans were parcellated into 360 bilateral cortical regions using the Human Connectome Project (HCP; [67]) template and 16 bilateral subcortical regions (amygdala, caudate, diencephalon, hippocampus, nucleus accumbens, pallidum, putamen, and thalamus) provided by Freesurfer’s ‘aseg’ parcellation template [68, 69]. Regional time series were averaged over all voxels within each parcel and bandpass filtered by the discrete wavelet transform, corresponding to frequency range 0.025-0.111 Hz [70].

After pre-processing and quality control of each individual scan, we retained regional time series for 330 cortical and 16 subcortical nodes. 30 cortical regions were excluded due to low regional mean signal, defined by a low Z-score of mean signal intensity in at least one participant (Z<-1.96; Supplementary Fig. 1 for details on retained regions). Individual functional connectivity matrices {346 × 346} were estimated by Pearson’s correlation for each possible of pair of nodes. Finally, we regressed each pairwise correlation or edge on the time-averaged head motion of each participant (mean FD). The residuals of this regression were the estimates of functional connectivity used for further analysis (Supplementary Fig. 3).

### Estimating parameters of adolescent development and testing sex differences

Previous work on this dataset did not find evidence for non-linear trajectories of development of functional connectivity between the majority of all possible pairs of regional nodes ([6]; Supplementary Text *“Non-Linear Effects of Age”*). Therefore we used a linear function to model the fixed effect of age on regional and edge-wise metrics of cortico-cortico, subcortico-cortical and cortico-subcortical functional connectivity (Supplementary Fig. 7), also including the fixed effect of site and a subject-specific intercept as a random effect, in linear mixed effects models fit separately for males and females (Supplementary Text *“Sex Stratified Analysis of Developmental Parameters”)*.

Baseline connectivity (*FC*_14_) was estimated as the predicted FC at age 14, the adolescent rate of change (*FC*_14-26_) as the slope of the model (Fig. 1). We calculated the maturational index (MI), as the Spearman correlation of edge-wise *FC*_14_ and *FC*_44-26_ (Fig. 2).

We parametrically tested for the significance of the sex difference in all parameters using a *Z*-test [71] (Supplementary Text *“Significance Testing of Sex Effects on Developmental Parameter”*).

### Enrichment analysis

We extracted the first component (PLS1) of a partial least squares regression of Δ*MI* on post mortem gene expression data from the Allen Human Brain Atlas collected from 6 donor brains (5 males) [43] (Fig. 4). We then used a median rank-based approach to assess the enrichment of PLS1 on several published gene lists [44]. This approach estimates the degree to which the spatial expression of PLS1 genes is related to a given gene set by comparing the observed (PLS1) median gene rank to a null distribution of median ranks from genes that were randomly sampled from the PLS component and matched by gene length (see Supplementary Text *“Enrichment Analysis”* for details). Thus if a gene set’s real median rank is significantly lower than expected by chance the gene set is associated with the bottom of PLS1 and if it is higher, its genes are enriched towards the top end of the component.

### FMRI connectivity in major depressive disorder

We constructed a MDD case-control map by conducting multiple *t*-tests for the difference in nodal weighted degree of functional connectivity between two groups of resting state fMRI data from an independent sample [38] of 46 healthy controls and 50 MDD cases (Supplementary Text *Co-location with Depression*, Supplementary Table 4). We then correlated the Δ*MI* map with the MDD case-control *t* map (Fig. 3).

## Data and code availability

Data used in this analysis will be made available upon publication. The analysis code can be found at: https://github.com/LenaDorfschmidt/sex_differences_adolescence.git.

## Acknowledgements

We appreciate Dr Varun Warrier’s advice on statistical methods for gene set enrichment analyses. We thank Dr Julia Spindel for sharing her expertise on epigenetic data. This study was funded by a collaborative award from the Wellcome Trust for the Neuroscience in Psychiatry Network at University College London and the University of Cambridge. Additional funding was provided by the National Institute of Health Research (NIHR) Cambridge Biomedical Research Centre. ETB is an NIHR Senior Investigator. L.D. was supported by the Gates Cambridge Trust. P.E.V. is a Fellow of MQ: Transforming Mental Health (MQF17_24) and of the Alan Turing Institute funded by EPSRC grant EP/N510129/1. R.A.I.B. was supported by a British Academy Post-Doctoral fellowship and the Autism Research Trust. S.E.M. was funded by a Fellowship from The Alan Turing Institute, London and a Henslow Fellowship at Lucy Cavendish College, University of Cambridge, funded by the Cambridge Philosophical Society. F.V. was supported by the Data to Early Diagnosis and Precision Medicine Industrial Strategy Challenge Fund, UK Research and Innovation (UKRI). Data were curated and analysed using a computational facility funded by an MRC research infra-structure award (MR/M009041/1) and supported by the NIHR Cambridge Biomedical Research Centre. The views expressed are those of the authors and not necessarily those of the NHS, the NIHR or the Department of Health and Social Care. Some data in this study was funded by an award from the Wellcome Trust (grant number: 104025/Z/14/Z) for the Neuroimmunology of Mood Disorders and Alzheimer’s Disease (NIMA) consortium, which was also funded by Janssen, GlaxoSmithKline, Lundbeck and Pfizer.

## Author Contributions

L.D., P.E.V., and E.T.B. designed research. L.D., R.A.I.B., and J.S. analyzed data. L.D., R.R.-G, F.V., M.G.K., P.E.V. performed research. J.S., R.A.I.B, S.E.M., M.G.B., and A.A. contributed new reagents<analytical tools. S.R.W. advised on statistical methods. E.T.B., P.J., R.D., I.G., and P.F. designed the NSPN study. N.A.H. designed the Biodep study. E.T.B. and L.D. wrote the paper.

## References

[1] Sowell, E. R., Thompson, P. M. & Toga, A. W. Mapping changes in the human cortex throughout the span of life. The Neuroscientist 10, 372–392 (2004).

[2] Giedd, J. N. Structural magnetic resonance imaging of the adolescent brain. Annals of the New York Academy of Sciences 1021, 77–85 (2004).

[3] Raznahan, A. et al. Patterns of coordinated anatomical change in human cortical development: a longitudinal neuroimaging study of maturational coupling. Neuron 72, 873–884 (2011).

[4] Vàša, F. et al. Adolescent Tuning of Association Cortex in Human Structural Brain Networks. Cerebral Cortex 28, 281–294 (2018).

[5] Vértes, P. E. et al. Gene transcription profiles associated with inter-modular hubs and connection distance in human functional magnetic resonance imaging networks. Philosophical Transactions of the Royal Society B: Biological Sciences 371, 20150362 (2016).

[6] Vàša, F. et al. Conservative and disruptive modes of adolescent change in human brain functional connectivity. Proceedings of the National Academy of Sciences 117, 3248–3253 (2020).

[7] Faravelli, C., Alessandra Scarpato, M., Castellini, G. & Lo Sauro, C. Gender differences in depression and anxiety: The role of age. Psychiatry Research 210, 1301–1303 (2013).

[8] Cyranowski, J. M., Frank, E., Young, E. & Shear, K. M. Adolescent Onset of the Gender Difference in Lifetime Rates of Major Depression. Archives of General Psychiatry 57, 21 (2000).

[9] Hankin, B. L. et al. Development of depression from preadolescence to young adulthood: Emerging gender differences in a 10-year longitudinal study. Journal of Abnormal Psychology 107, 128–140 (1998).

[10] Paus, T., Keshavan, M. & Giedd, J. N. Why do many psychiatric disorders emerge during adolescence? Nature Reviews Neuroscience 9, 947–957 (2008).

[11] Biswal, B., FZerrin Yetkin, Haughton V. M. & Hyde, J. S. Functional connectivity in the motor cortex of resting human brain using echo-planar mri. Magnetic Resonance in Medicine 34, 537–541 (1995).

[12] Fornito, A., Zalesky, A. & Bullmore, E. T. Fundamentals of brain network analysis / Alex Fornito, Andrew Zalesky, Edward T. Bullmore (Elsevier/Academic Press Amsterdam ; Boston, 2016).

[13] Whitaker, K. J. et al. Adolescence is associated with genomically patterned consolidation of the hubs of the human brain connectome. Proceedings of the National Academy of Sciences of the United States of America 113, 9105–10 (2016).

[14] Mills, K. L., Goddings, A.-L., Clasen, L. S., Giedd, J. N. & Blakemore, S.-J. The Developmental Mismatch in Structural Brain Maturation during Adolescence. Developmental Neuroscience 36, 147–160 (2014).

[15] Fair, D. A. et al. Functional Brain Networks Develop from a “Local to Distributed” Organization. PLoS Computational Biology 5, e1000381 (2009).

[16] Fair, D. A. et al. Development of distinct control networks through segregation and integration. Proceedings of the National Academy of Sciences 104, 13507–13512 (2007).

[17] Dosenbach, N. U. F. et al. Prediction of Individual Brain Maturity Using fMRI. Science 329, 1358–1361 (2010).

[18] Power, J. D., Barnes, K. A., Snyder, A. Z., Schlaggar, B. L. & Petersen, S. E. Spurious but systematic correlations in functional connectivity MRI networks arise from subject motion. NeuroImage 59, 2142–54 (2012).

[19] Satterthwaite, T. D. et al. Impact of in-scanner head motion on multiple measures of functional connectivity: Relevance for studies of neurodevelopment in youth. NeuroImage 60, 623 – 632 (2012).

[20] Satterthwaite, T. et al. Heterogeneous impact of motion on fundamental patterns of developmental changes in functional connectivity during youth. NeuroImage 83, 45 – 57 (2013).

[21] Power, J. D., Plitt, M., Kundu, S. J. G. P., Voon, V. & Martin, P. A. B. A. Ridding fmri data of motion-related influences: Removal of signals with distinct spatial and physical bases in multiecho data. Proceedings of the National Academy of Sciences 115, E2105–E2114 (2018).

[22] Murphy, K., Birn, R. M., Handwerker, D. A., Jones, T. B. & Bandettini, P. A. The impact of global signal regression on resting state correlations: are anti-correlated networks introduced? NeuroImage 44, 893–905 (2009).

[23] Schölvinck, M. L., Maier, A., Ye, F. Q., Duyn, J. H. & Leopold, D. A. Neural basis of global resting-state fmri activity. Proceedings of the National Academy of Sciences 107, 10238–10243 (2010).

[24] Maknojia, S., Churchill, N. W., Schweizer, T. A. & Graham, S. J. Resting state fmri: Going through the motions. Front Neurosci 13 (2019).

[25] Filippi, M. et al. The organization of intrinsic brain activity differs between genders: A resting-state fMRI study in a large cohort of young healthy subjects. Human Brain Mapping 34, 1330–1343 (2013).

[26] Tomasi, D. & Volkow, N. D. Aging and functional brain networks. Molecular Psychiatry 17, 549–558 (2012).

[27] Allen, E. A. et al. A baseline for the multivariate comparison of resting-state networks. Frontiers in systems neuroscience 5, 2 (2011).

[28] Biswal, B. B. et al. Toward discovery science of human brain function. Proceedings of the National Academy of Sciences of the United States of America 107, 4734–9 (2010).

[29] Bluhm, R. L. et al. Default mode network connectivity: effects of age, sex, and analytic approach. NeuroReport 19, 887–891 (2008).

[30] Scheinost, D. et al. Sex differences in normal age trajectories of functional brain networks. Human Brain Mapping 36, 1524–1535 (2015).

[31] Weissman-Fogel, I., Moayedi, M., Taylor, K. S., Pope, G. & Davis, K. D. Cognitive and default-mode resting state networks: Do male and female brains “rest” differently? Human Brain Mapping 31, /a-n/a (2010).

[32] Alarcón, G., Cservenka, A., Rudolph, M. D., Fair, D. A. & Nagel, B. J. Developmental sex differences in resting state functional connectivity of amygdala sub-regions. NeuroImage 115, 235–244 (2015).

[33] Kilpatrick, L. A., Zald, D. H., Pardo, J. V. & Cahill, L. F. Sex-related differences in amygdala functional connectivity during resting conditions. NeuroImage 30, 452–461 (2006).

[34] Zhang, C. et al. Sex and Age Effects of Functional Connectivity in Early Adulthood. Brain Connectivity 6, 700–713 (2016).

[35] Zhang, C., Dougherty, C. C., Baum, S. A., White, T. & Michael, A. M. Functional connectivity predicts gender: Evidence for gender differences in resting brain connectivity. Human Brain Mapping 39, 1765–1776 (2018).

[36] Casanova, R., Whitlow, C. T., Wagner, B., Espeland, M. A. & Maldjian, J. A. Combining graph and machine learning methods to analyze differences in functional connectivity across sex. The open neuroimaging journal 6, 1–9 (2012).

[37] Kiddle, B. et al. Cohort Profile: The NSPN 2400 Cohort: a developmental sample supporting the Wellcome Trust NeuroScience in Psychiatry Network. International Journal of Epidemiology 47, 18–19g (2017).

[38] Kitzbichler, M. G. et al. Peripheral inflammation is associated with micro-structural and functional connectivity changes in depression-related brain networks. medRxivs (2020).

[39] Patel, A. X. et al. A wavelet method for modeling and despiking motion artifacts from resting-state fMRI time series. Neuroimage 95, 287–304 (2014).

[40] Yeo, B. T. et al. The organization of the human cerebral cortex estimated by intrinsic functional connectivity. Journal of neurophysiology 106, 1125–1165 (2011).

[41] Yarkoni, T., Poldrack, R. A., Nichols, T. E., Essen, D. C. V. & Wager, T. D. Large-scale automated synthesis of human functional neuroimaging data. Nature methods 8, 665–670 (2011).

[42] Morgan, S. E. et al. Cortical patterning of abnormal morphometric similarity in psychosis is associated with brain expression of schizophrenia-related genes. Proceedings of the National Academy of Sciences of the United States of America 116, 9604–9609 (2019).

[43] Hawrylycz, M., Lein, E., Guillozet-Bongaarts, A. & et al. An anatomically comprehensive atlas of the adult human brain transcriptome. Nature 391–399 (2012).

[44] Seidlitz, J. et al. Transcriptomic and cellular decoding of regional brain vulnerability to neurogenetic disorders. Nature Communications 11 (2020).

[45] Zhu, Y. et al. Spatiotemporal transcriptomic divergence across human and macaque brain development. Science 362 (2018).

[46] Xu, X., Wells, A. B., O’Brien, D. R., Nehorai, A. & Dougherty, J. D. Cell type-specific expression analysis to identify putative cellular mechanisms for neurogenetic disorders. Journal of Neuroscience 34, 1420–1431 (2014).

[47] Lake, N. et al. Integrative single-cell analysis of transcriptional and epigenetic states in the human adult brain. Nature Biotechnology 36, 70–80 (2018).

[48] Polioudakis, D. et al. A single-cell transcriptomic atlas of human neocortical development during midgestation. Neuron 103, 785 - 801.e8 (2019).

[49] Li, M. et al. Integrative functional genomic analysis of human brain development and neuropsychiatric risks. Science 362 (2018).

[50] Wray, N. R., Ripke, S., Mattheisen, M. & et al. Genome-wide association analyses identify 44 risk variants and refine the genetic architecture of major depression. Nature Genetics 668–681 (2018).

[51] Sey, N. Y. A. et al. A computational tool (h-magma) for improved prediction of brain-disorder risk genes by incorporating brain chromatin interaction profiles. Nature Neuroscience 583–593 (2020).

[52] Edwards, S. L., Beesley, J., French, J. D. & Dunning, A. M. Beyond gwass: Illuminating the dark road from association to function. The American Journal of Human Genetics 93, 779–797 (2013).

[53] Phoenix, C. H., Goy, R. W., Gerall, A. A. & Young, W. C. Organizing action of prenatally administered testosterone propionate on the tissue mediating mating behavior in the female guinea pig. Endocrinology 65, 369–382 (1959).

[54] Carruth, L., Reisert, I. & Arnold, A. Sex chromosome genes directly affect brain sexual differentiation. Nature neuroscience 5, 933–4 (2002).

[55] Arnold, A. P. et al. Minireview: Sex Chromosomes and Brain Sexual Differentiation. Endocrinology 145, 1057–1062 (2004).

[56] McCarthy, M. M. & Arnold, A. P. Reframing sexual differentiation of the brain. Nature neuroscience 14, 677–83 (2011).

[57] Cullen, K. R. et al. A preliminary study of functional connectivity in comorbid adolescent depression. Neuroscience Letters 460, 227–231 (2009).

[58] Connolly, C. G. et al. Resting-State Functional Connectivity of Subgenual Anterior Cingulate Cortex in Depressed Adolescents. Biological Psychiatry 74, 898–907 (2013).

[59] Anderson, K. M. et al. Convergent molecular, cellular, and neural signatures of major depressive disorder. bioRxiv (2020).

[60] Kaczkurkin, A. N., Raznahan, A. & Satterhwaite, T. D. Sex differences in the developing brain: insights from multimodal neuroimaging. Neuropsychopharmacology 71–85 (2019).

[61] Cahill, L. Why sex matters for neuroscience. Nat Rev Neurosci 477–484 (2006).

[62] Eliot, L., Ahmed, A., Khan, H. & Patel, J. Dump the “dimorphism”: Comprehensive synthesis of human brain studies reveals few male-female differences beyond size. Neuroscience & Biobehavioral Reviews (2021).

[63] Barth, M., Reichenbach, J. R., Venkatesan, R. & Haacke, E. M. M. E. High-resolution, multiple gradient-echo functional MRI at 1.5 T. Magnetic Resonance Imaging 17, 321–329 (1999).

[64] Kundu, P., Inati, S. J., Evans, J. W., Luh, W.-M. & Bandettini, P. A. Differentiating BOLD and non-BOLD signals in fMRI time series using multi-echo EPI. Neuroimage 60, 1759–1770 (2012).

[65] Kundu, P. et al. Integrated strategy for improving functional connectivity mapping using multiecho fMRI. Proceedings of the National Academy of Sciences of the United States of America 110, 16187–92 (2013).

[66] Cox, R. W. AFNI: Software for Analysis and Visualization of Functional Magnetic Resonance Neuroimages. Computers and Biomedical Research 29, 162–173 (1996).

[67] Glasser, M. F. et al. A multi-modal parcellation of human cerebral cortex. Nature 536, 171–178 (2016).

[68] Filipek, P. A., Richelme, C., Kennedy, D. N. & Caviness, V. S. The Young Adult Human Brain: An MRI-based Morphometric Analysis. Cerebral Cortex 4, 344–360 (1994).

[69] Fischl, B. et al. Automated labeling of neuroanatomical structures in the human brain. Cell Press 33, 341–355 (2002).

[70] Bullmore, E. T. et al. Wavelets and functional magnetic resonance imaging of the human brain. NeuroImage 23, S234–S249 (2004).

[71] Paternoster, R., Brame, R., Mazerolle, P. & Piquero, A. Using the Correct Statistical Test for the Equality of Regression Coefficients. Criminology 36, 859–866 (1998).

[72] Vàša, F. et al. Data for “Conservative and disruptive modes of adolescent change in human brain functional connectivity”. figshare (2020). URL https://figshare.com/articles/dataset/Data_for_Conservative_and_disruptive_modes_of_adolescent_change_in_human_brain_functional_connectivity_/11551602.

