## Supplementary for "Sexually divergent development of depression-related brain networks during healthy human adolescence"

** *A complete list of the NSPN Consortium can be found in the SI Appendix.*

*20/4/2021*

**NSPN Consortium Author List** **51**

**NIMA Consortium Author List** **52**

**References** **55**

#### Data

*Data Collection*

The data was collected as part of the Neuroscience in Psychiatry Network (NSPN), a joint initiative by the University of Cambridge and the University College London, with the aim of collecting a general population sample in an accelerated longitudinal design to measure developmental change in the population of Greater London and Cambridgeshire, broadly representative of the populations of England and Wales. The final sample consists of 2402 subjects, aged 14 to 26, that underwent a self-assessment of well-being and demographics, with a subset undergoing functional and structural imaging procedures as detailed below.
All participants aged 16 and older provided informed written consent for each aspect of the study, and parental consent was obtained for those aged 14–15 years. The study was ethically approved by the National Research Ethics Service and was conducted in accordance with NHS research governance standards.

*MRI Sample*

A subsample of 306 adolescents was invited to undergo functional and structural neuroimaging assessments. The exclusion criteria for this sample included: a current or past history of neurological disorder or learning disability, and current treatment for psychiatric disorder or drug or alcohol dependence. Each participant in the scanning sample was invited to provide MRI data on at least two occasions; at baseline and at follow-up 12-18 months later, with 29 participants additionally invited to attend follow-up scanning six months after baseline.. The fMRI scan was the first in a series of scans collected in each scanning session which also included structural MRI using the multi-parameter mapping (MPM) sequence (1), and diffusion weighted imaging. A total of 556 functional scans were available after repeated scanning and quality control of this cohort.

| Sex | # Scans | # Scanned | | | At Baseline | | | | # Subj./Agebin | | | | |
|---|---|---|---|---|---|---|---|---|---|---|---|---|---|
| | | 1 | 2 | 3 | $\mu$ Age | $\sigma$ Age | $\mu$ FD | $\sigma$ FD | 1 | 2 | 3 | 4 | 5 |
| female | 259 | 54 | 86 | 11 | 19.8 | 2.9 | 0.11 | 0.05 | 34 | 39 | 24 | 32 | 22 |
| male | 261 | 41 | 98 | 8 | 19.2 | 3.8 | 0.13 | 0.05 | 32 | 33 | 24 | 35 | 23 |

**Table S1 NSPN sample overview:** A total of N = 298 healthy young people participated in an accelerated longitudinal MRI study. The recruitment was balanced for sex in each of five age-defined strata. Subjects were scanned between 1 and 3 times with scans taking place at baseline, 6 and 18 months later. FD = framewise displacement, a measure of head movement in mm, was significantly greater in males compared to females on average over all ages, and in the youngest two age strata specifically ($P < 0.05, uncorrected$).

*MRI Data Acquisition*

Functional MRI data were acquired at three sites, on three identical 3T Siemens MRI scanners (Magnetom TIM Trio, VB17 software version), with a standard 32-channel radio-frequency (RF) receive head coil and RF body coil for transmission using a multi-echo echo-planar (EPI) imaging sequence (2) with the following scanning parameters: repetition time (TR): 2.42s; GRAPPA with acceleration factor 2; flip angle: 90; matrix size: 64x64x34; FOV: 240x240mm; in-plane resolution: 3.75x3.75 mm; slice thickness: 3.75 mm with 10% gap, sequential slice acquisition, 34 oblique slices; bandwidth 2368 Hz/pixel; echo times (TE) 13, 30.55 and 48.1 ms; total scan time 11 minutes.

*MRI Data Preprocessing*

Freesurfer v5.3.0 was used to process individual structural scans with a pipeline compromising skull-stripping, segmentation of cortical grey and white matter and reconstruction of the cortical surface and grey-white matter boundary (3). Subsequently, all scans were stringently quality controlled by re-running the reconstruction algorithm after the addition of control points and white matter edits as previously described (4; 5).

AFNI was used for basic preprocessing of functional MRI scans as follows: All volumes acquired during steady-state equilibration (15 s) were discarded. Motion correction parameters and parameters for anatomical-functional coregistration were calculated from the images acquired with TE = 30.55 ms. The first volume after equilibration was used as the base EPI image. Matrices for de-obliquing and six-parameter rigid body motion correction were computed. Then, 12-parameter affine anatomical-functional coregistration was computed using the LPC cost functional (6), with the EPI base image as the LPC weight mask. Matrices for de-obliquing, motion correction, and anatomical-functional coregistration were combined into a single alignment matrix using the concatenation approach from the AFNI tool $align_{e}pi_{a}nat.py$. The images for each TE were then slice-time corrected and spatially aligned through application of the alignment matrix. Coregistration of structural and functional scans was visually assessed.

Functional MRI data were preprocessed using multi-echo independent component analysis (ME-ICA;(7; 8)) which identifies and removes sources of variance in the times series that do not scale linearly with TE and are therefore not representative of the BOLD signal. The retained independent components, representing BOLD contrast, were optimally recomposed to generate a broadband denoised fMRI time series at each voxel. Regional time series were averaged over all voxels within each parcel and bandpass filtered by the discrete wavelet transform, corresponding to a frequency range of 0.025-0.111 Hz (9).

An overall estimate of head motion by each participant, mean framewise displacement (FD), was calculated from the six motion parameter time series (three rotation and three translation parameters) estimated during scan re-alignment. More specifically, framewise displacement was calculated as:

$$FD_t = \sum_d |d_{t-1} * d_t| + 50 * \frac{\pi}{180} * \sum_d |r_{t-1} * r_t| \tag{1}$$

Scans were parcellated into 360 bilateral cortical regions using the Human Connectome Project (HCP; (10)) template and 16 bilateral sub-cortical regions (amygdala, caudate, diencephalon, hippocampus, nucleus accumbens, pallidum, putamen, and thalamus) defined by Freesurfer's *aseg* parcellation (11). After within-subject preprocessing and quality control, we retained regional time series for 330 cortical and 16 subcortical nodes. 30 cortical regions were excluded due to low regional mean signal (Z <-1.96); see SI Fig. S1 for a map of retained regions. Individual functional connectivity matrices $\{346 \times 346\}$ were estimated by Pearson's correlation for each possible of pair of nodes. Finally, we regressed each pairwise correlation or edge on the time-averaged head motion of each participant (mean FD). The residuals of this regression were the estimates of functional connectivity used for further analysis.

*Exclusion Criteria*

A total of 36 scans were excluded.17 scans were excluded due to high in-scanner motion (defined as mean FD $>$ 0.3 mm or maximum FD $>$ 1.3 mm), 9 due to coregistration errors, 7 due to a lack of convergence of the ME-ICA algorithm, 2 due to parcellation errors, and 1 due to extensive signal dropout (as defined above).

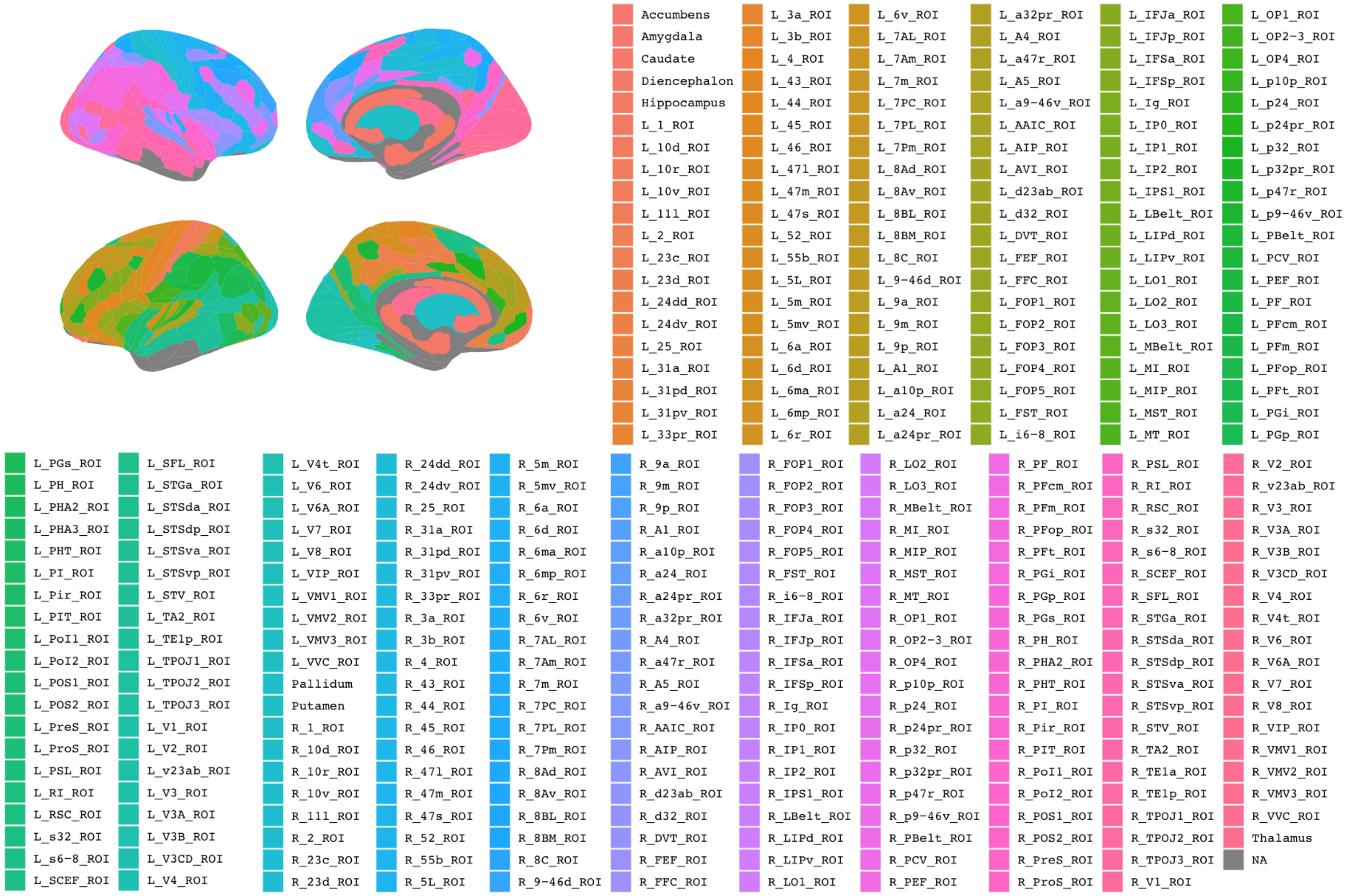

**Fig. S1: ROIs included after preprocessing and quality control:** We parcellated the scans into 376 ROIs (360 cortical regions in the Human Connectome Atlas, and 16 subcortical regions from the Freesurfer Software). 30 cortical regions were excluded due to low regional mean signal (Z<-1.96) in at least one scan. These regions were: *L 10pp, L 13l, L OFC, L EC, L H, L PeEc, L PHA1, L TGd, L TE1a, L TE2a, L TF, L TE2p, L pOFC, L TGv, L TE1m, R 10pp, R 13l, R OFC, R EC, R H, R PeEc, R PHA1, R PHA3, R TGd, R TE2a, R TF, R TE2p, R pOFC, R TGv, R TE1m.*

#### Head Motion

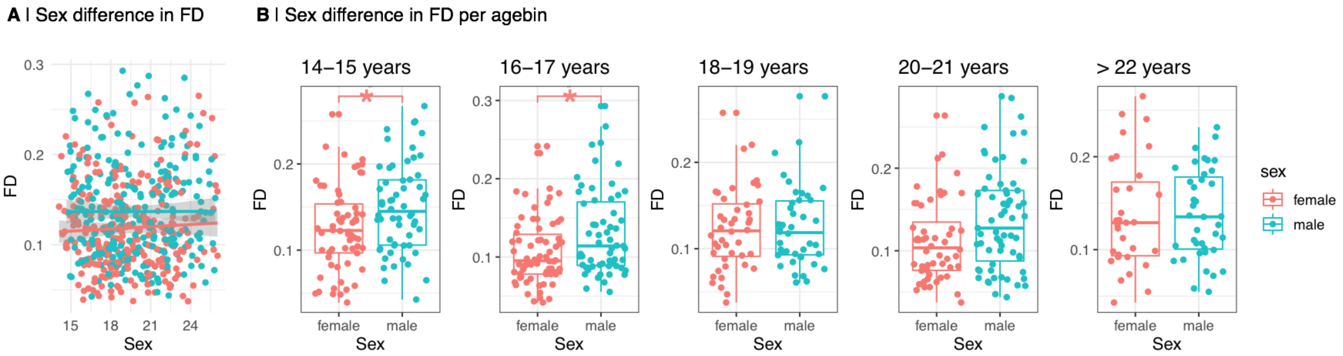

**Fig. S2: Sex differences in framewise displacement:** (A) Across all agebins, males showed higher framewise displacement (FD) than females ($P_{Sex} < 0.05, t(296) = 3.25$). (B) Males showed increased FD in the first two agebins ($P_{Sex} < 0.05$), however the effect did not survive correction for multiple comparisons.

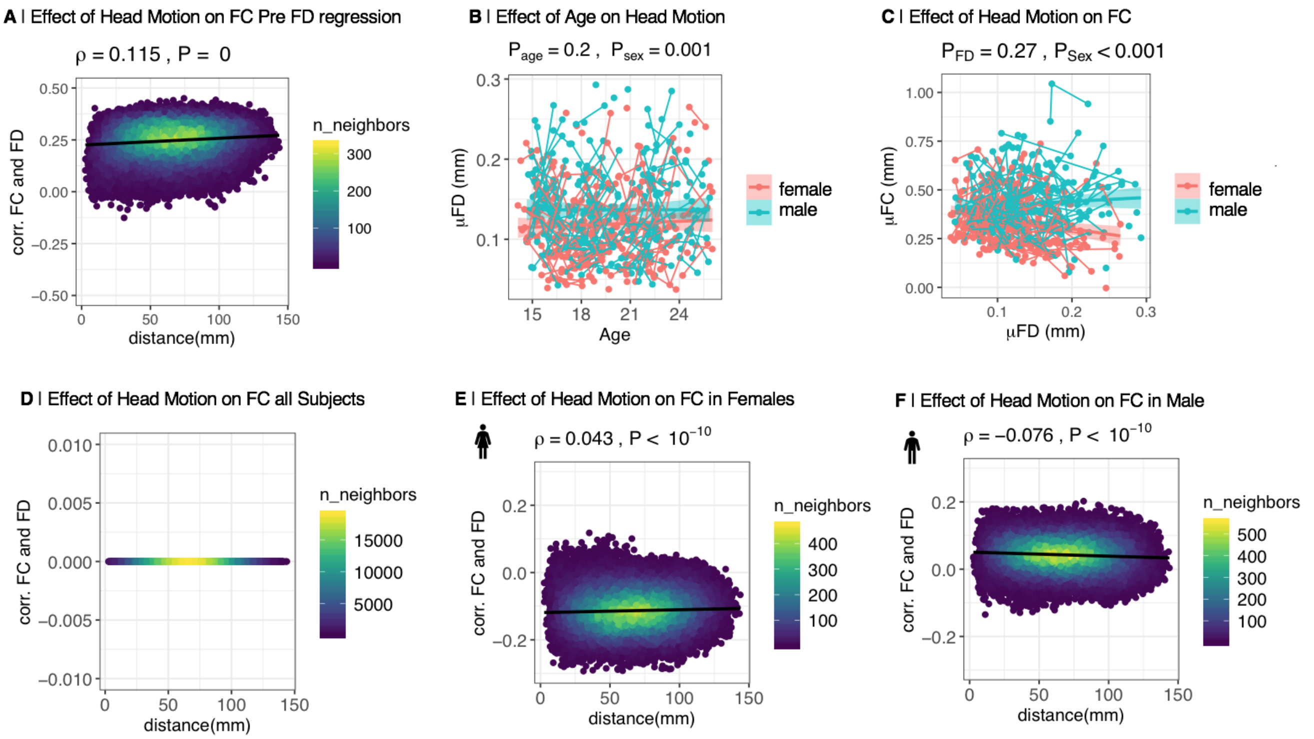

**Fig. S3: Effect of head motion (FD) on functional connectivity (FC) in the original sample:** (A) After ME-ICA preprocessing, we found a weak relationship between the correlation of FC and head motion (across participants) and the Euclidean distance spanned by edges. Further, the average edge-wise correlation between FC and motion is non-zero. (B) To remove the dependence of FC on motion in our sample, mean FD was regressed from each edge; the residuals constitute participant-specific FD-corrected FC, with intercepts retained to maintain the relative importance of edges across the group as well as the interpretability of FC values. Thus in the full sample, average head motion, quantified as mean frame-wise displacement (FD), did not change with age ($P_{age} = 0.2, t(220) = 125$). However, there was a weak, but significant effect of sex on FD ($\beta_{sex} = 0.02, P_{sex} < 0.01, t(296) = 3.3$). (C) The effect of participants' motion (across participants) on global FC was not significant ($P_{FD} = 0.27$). (D) By definition, there was no effect of distance on the correlation between FC and motion, and the average edge-wise correlation between FC and motion was non-zero (as evidenced by a non-zero intercept of the linear regression on the y-axis). (E) However, since our motion correction was performed across all subjects, we still observed weak, but significant effects of distance on the correlation of FC and FD for females ($\rho = 0.04, P < 10^{-10}$) (F) and males ($\rho = -0.08, P < 10^{-10}$) (G) separately.

#### Analysis of Effects on Parameters of Adolescent Brain Development

*Non-Linear Effects of Age*

It is conceivable that there could be non-linear effects of age on functional connectivity. However, in previous work on the same data set (12), we investigated potential non-linear effects of age and found no substantial evidence for non-linearity in these data. Specifically, Váša et al. (2020), fitted smoothing splines (generalized additive mixed models) to edge-wise trajectories of functional connectivity development, using the "gamm" function in R, with the effect of age modelled as the weighted sum of 10 cubic b-splines with knots placed at quantiles of the data and smoothing optimized using restricted maximum likelihood (13). This modeling strategy is adaptive to the non-linearity of age effects, such that non-linear trajectories will be best fitted by spline functions with degrees of freedom greater than 2, whereas linear trajectories will be best fitted by spline functions with 2 df, analogous to the intercept and gradient parameters of a simple linear model. We found that approximately 70% of all edges had linear trajectories that were best fit by spline functions with 2 df (Supplementary Figure S8, (12)). Moreover, there was no evidence from this analysis that the minority of edges with non-linear trajectories were concentrated on anatomically specific brain regions or systems. For these reasons, we adopted a linear function for age-related effects on functional connectivity in our modeling of these data (Equation 2). However, to mitigate any residual concerns that non-linearity of age-related changes in functional connectivity might confound sex differences in MI, we examined the relationship between $\Delta MI$ and the nodal mean degrees of freedom for spline functions of age. As shown in Figure S4 there was no significant correlation ($\rho = 0.006, P = 0.9$). This indicates that there is no evidence for a relationship between non-linear trajectories of functional connectivity development and sex differences in maturational index.

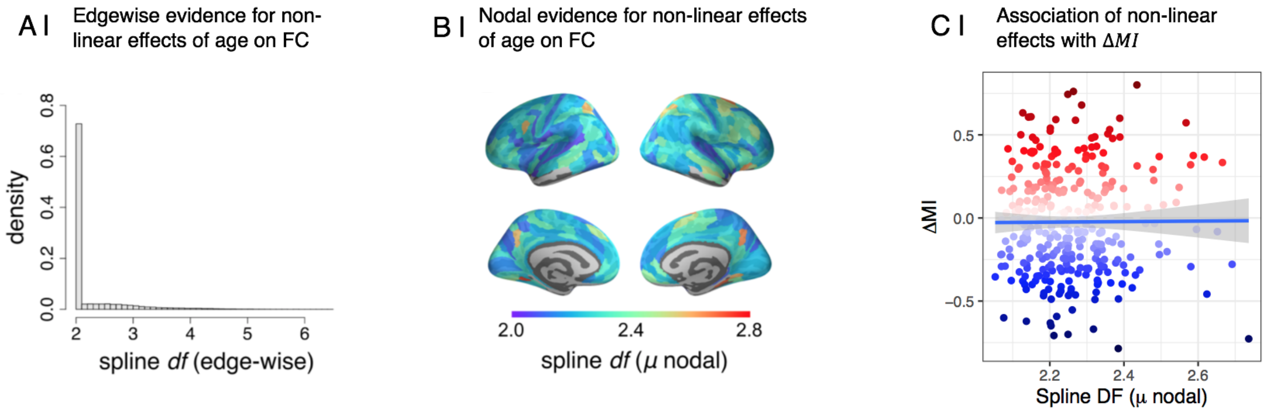

**Fig. S4: Non-linear Effects of Age on FC:** In previous work on this sample, Váša et al., fitted locally adaptive mixed effect smoothing splines to edgewise trajectories of functional connectivity development to inspect potential non-linear effects of age on FC. In these models, non-linear trajectories will be best fitted by spline functions with degrees of freedom greater than 2, whereas linear trajectories will be best fitted by spline functions with 2 df, analogous to the intercept and gradient parameters of a simple linear model. (A) Distribution of effective df of smoothing splines across edges: 71.7% trajectories had df $< 2.1$, suggesting that most trajectories are linear. (B) Cortical distribution of average nodal df (averaged across all of a node's edges). (C) Lastly, we correlated the cortical map of sex differences in maturational index, $\Delta MI$, with the cortical map of average nodal df. We show that there is no association between evidence for non-linearities in functional connectivity development with age, as indicated by an effective df $> 2$ ($\rho = 0.006, p = 0.9$). Panels A and B are reproduced, with permission, from the Supplementary Information for (12).

*Sex Stratified Analysis of Developmental Parameters*

For the principal analyses reported in this paper, we used a sex-stratified approach to analysis. This means we fit a linear mixed effects (LME) model to estimate $FC_{14}$ and $FC_{14-26}$ separately for each sex, as detailed below.
For each sex, we predicted functional connectivity at edge level using linear mixed effects models. These models included age as the main fixed effect of interest and scanner site as a fixed effect covariate, as well as a subject-specific intercept as a random effect, as follows:

$$FC_{edge} \sim 1 + \beta_{age} * age + \beta_{site} * site + \gamma_{subject} * (1|subject) + \epsilon \quad (2)$$

where FC refers to the functional connectivity at edge level, $\beta$ refers to coefficients for the fixed effects, $\gamma_s ubject$ refers to the coefficients for random effects, and $\epsilon$ represents the residual error.

We then derived baseline connectivity at age 14 from 2 as:

$$FC = \beta_{age} * 14 + \beta_{site_2} * (1/3) + \beta_{site_3} * (1/3) \quad (3)$$

Whereas the adolescent rate of change is simply the $\beta$ coefficient of age from 2 as:

$$FC_{14-26} = \beta_{age} \quad (4)$$

We then combined the sex specific estimates of these two developmental parameters to estimate a sex-specific estimate of MI. Thus in each sex we evaluated at each node the linear relationship (using Spearman's $\rho$) between the ranked edge-wise parameters $FC_{14}$ and $FC_{14-26}$ for each sex, e.g.:

$$MI_{node} = \frac{cov(FC_{14}, FC_{14-26})}{\sigma_{FC_{14}}, \sigma_{FC_{14-26}}} \quad (5)$$

Where $\sigma_{FC_{14}}$ and $\sigma_{FC_{14-26}}$ are the standard deviations of the ranked variables.

Finally, between-sex differences were estimated by subtracting each male-specific parameter from the corresponding female-specific parameter:

$$\Delta FC_{14} = FC_{14_{female}} - FC_{14_{male}} \quad (6)$$

$$\Delta FC_{14-26} = FC_{14-26_{female}} - FC_{14-26_{male}} \quad (7)$$

$$\Delta MI = MI_{female} - MI_{male} \quad (8)$$

Our principal reason for choosing a sex stratified approach was that it allows the variance of the random effects estimated in Equation 2, Var($\gamma_{subject}$), to differ between sexes. As shown in Fig. S5, the distributions of random effects were indeed not identical in males and females – males were more variable. Higher variance of random effects was negatively correlated with lower residual variance (denoted $\epsilon$ in 2) in both sexes; but the strength of correlation was greater in females than males, as shown in Fig S5B. Thus, although the between-sex difference in random effects variance was not statistically significant by a permutation test (Fig S5C) there was a degree of difference which would influence the residual variance and therefore the significance of the standardised developmental parameters. It is for this reason that we principally used the sex-stratified approach to linear mixed effects modeling of these longitudinal data.

*Age x Sex Effects on Maturational Index*

To demonstrate robustness of our results to an alternative modeling strategy, we also analysed all the data (male and female combined) using a linear mixed effects model to estimate the main effects of age and sex, and the age-by-sex interaction effect, on functional connectivity (FC) at each edge:

$$FC_{edge}\ 1 + \beta_{age} * age + \beta_{sex} * sex + \beta_{age*sex} * age * sex + \beta_{site} * site + \gamma_{subject} * (1|subject) + \epsilon \quad (9)$$

where FC refers to the functional connectivity at edge level, $\beta$ refers to coefficients for the fixed effects, $\gamma$ refers to coefficients for random effects and $\epsilon$ represents the residual error.

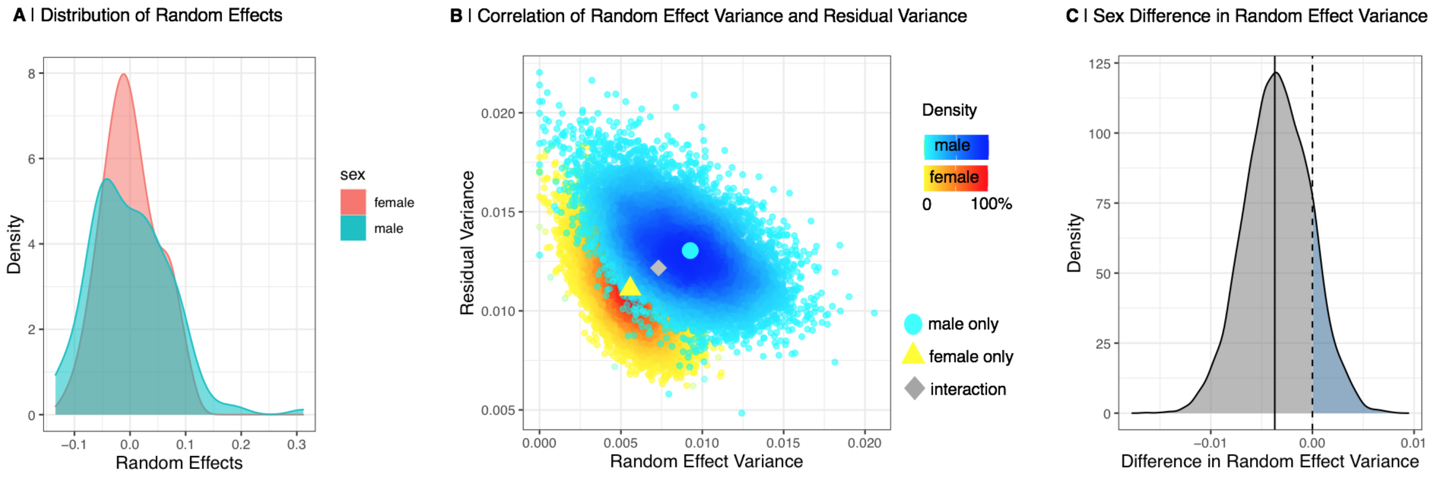

**Fig. S5: Random effect variance:** We estimated the random effects of sex on global functional connectivity in a sex stratified approach. we employed a bootstrap approach, resampling subjects with replacement 10,000 times within each age bin, thus creating a distribution of random effect and residual variance. (A) Distribution of random effects estimated in a sex stratified approach. (B) Correlation of random effect variance and residual variance for males and females estimated in a bootstrap approach. The real male and female, as well as the random effect variance of an interaction model are marked. This plot shows a clear separation of male and female random effect variance. (C) We estimated whether the random effect variance of males and females was equal. In a permutation approach, we estimated whether the difference in random effect variance between males and females significantly differed from 0. We find a p-value of $p = 0.14$.

On this basis, we can then estimate FC14 for males and females as follows:

$$FC14_{female} = \beta_{age} * 14 + \beta_{sex} * 0 + \beta_{age*sex} * 0 * 14 + \beta_{site_2} * (1/3) + \beta_{site_3} * (1/3) \tag{10}$$

$$FC14_{male} = \beta_{age} * 14 + \beta_{sex} * 1 + \beta_{age*sex} * 1 * 14 + \beta_{site_2} * (1/3) + \beta_{site_3} * (1/3) \tag{11}$$

And likewise we can estimate $FC_{14-26}$ for males and females:

$$FC_{14-26_female} = \beta_{age} + \beta_{age*sex} * 0 \tag{12}$$

$$FC_{14-26_male} = \beta_{age} + \beta_{age*sex} * 1 \tag{13}$$

Finally these sex-specific estimates of $FC_{14}$ and $FC_{14-26}$ can be combined to estimate sex-specific estimates of MI and the between-sex difference in MI, $\Delta MI$, as:

$$\Delta MI = MI_{female} - MI_{male} \tag{14}$$

As in our principal analysis, we tested the significance of the sex difference in MI by a parametric approach, comparing the slopes of the regression of $FC_{14}$ and $FC_{14-26}$ (14).
We found that the correlation between our principal sex—stratified results and the results of this alternative analysis, based on a fixed term for the sex-by-age interaction were nearly identical, with parameters estimated by sex-stratified and sex-by-age interaction models demonstrating a high degree of correlation, ($\rho > 0.9$; SI Fig. S6). Thus we conclude that our principal results from sex-stratified modelling are robust to an alternative modelling strategy that explicitly includes a fixed term for sex-by-age interaction. However, we continue to prefer the sex-stratified approach because of its greater adaptivity to the evident between-sex differences in random effects variance (SI Fig. S5).

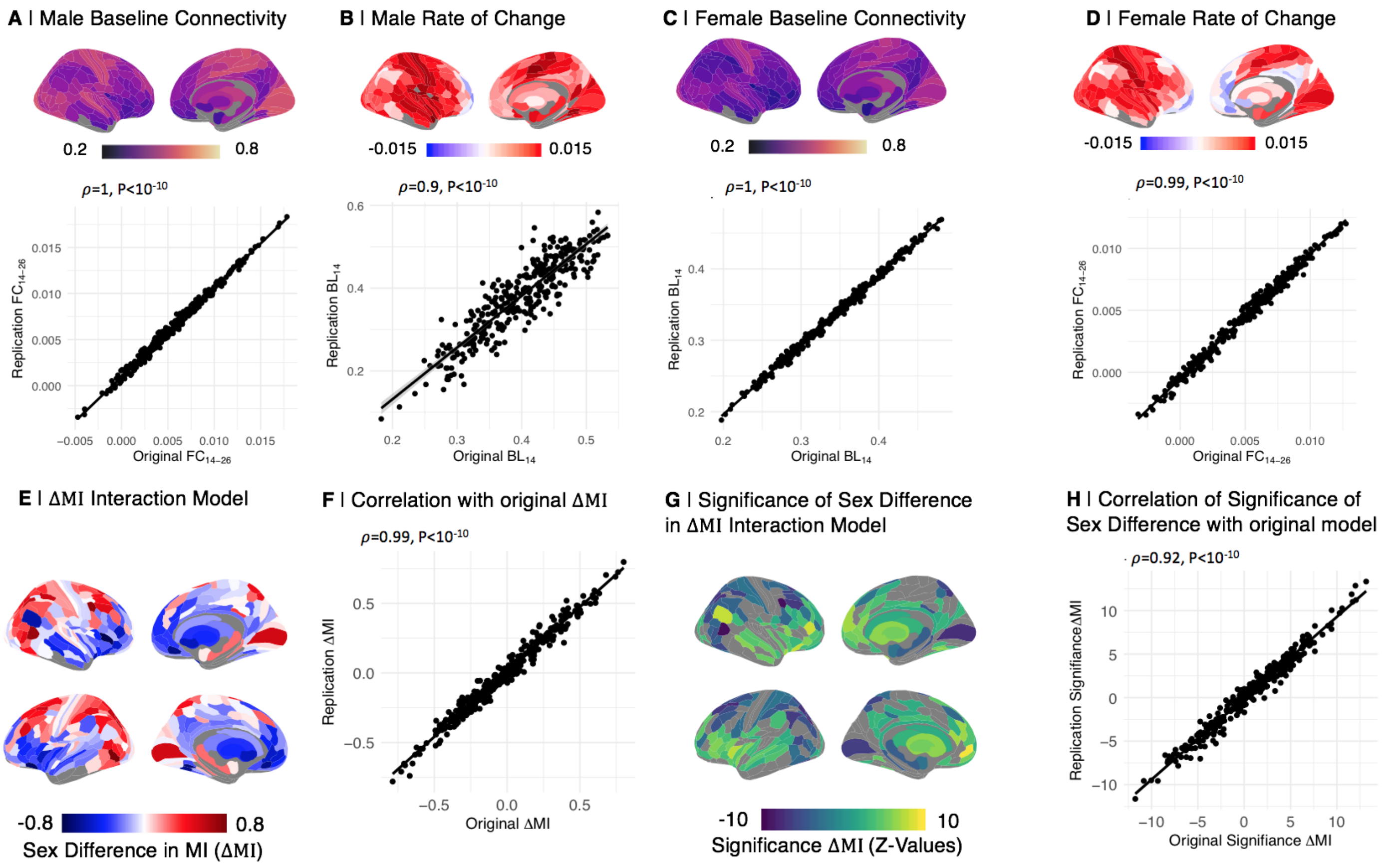

**Fig. S6: Interaction Model:** We modeled functional connectivity maturational in a joint model for males and females and included an interaction term for the interaction of age and sex. (A) (B)-(E) From this model, we derived updated $FC_{14}$ and $FC_{14-26}$ for males and females. We find that those updated measures are highly correlated with our original measures from the sex stratified approach. (F)-(G) We further find that the sex difference in maturational index, $\Delta MI$, as well as the map of the effect size of this sex differnece (z-values), are highly correlated with our main analysis.

*Significance Testing of Sex Effects on Developmental Parameter*

We parametrically tested for sex difference in baseline connectivity ($FC_{14}$) and the adolescent rate of change per year ($FC_{14-26}$) respectively, as:

$$z = \frac{p_{female} - p_{male}}{SE_{p_{female} - p_{male}}} \tag{15}$$

where $p_{female}$ and $p_{male}$ are the respective parameters of developmental change ($FC_{14}$,$FC_{14-26}$,$MI$) and $SE_{p_{female}-p_{male}}$ is the standard error of the difference in developmental parameters, calculated from the respective models.

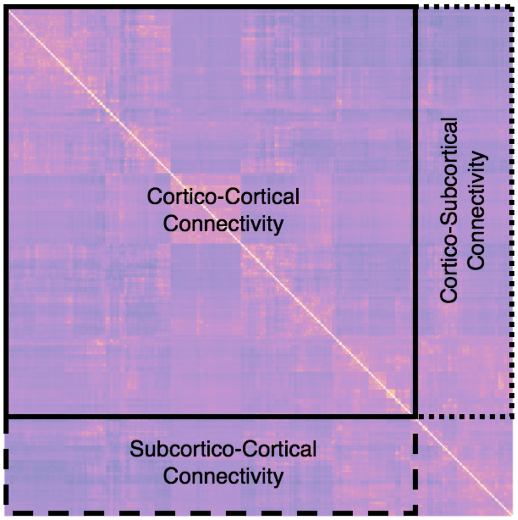

**Fig. S7: Functional connectivity estimation:** We estimated functional connectivity by averaging the strength of each node's edges across different parts of the matrix: (1) cortico-cortical connectivity was estimated as the mean over a cortical node's connections with all 330 other cortical nodes; (2) cortico-subcortical connectivity was estimated as the mean over a cortical node's connections with all 16 subcortical nodes; (3) subcortico-cortical connectivity was estimated as the mean over a subcortical node's connections with all 330 cortical nodes.

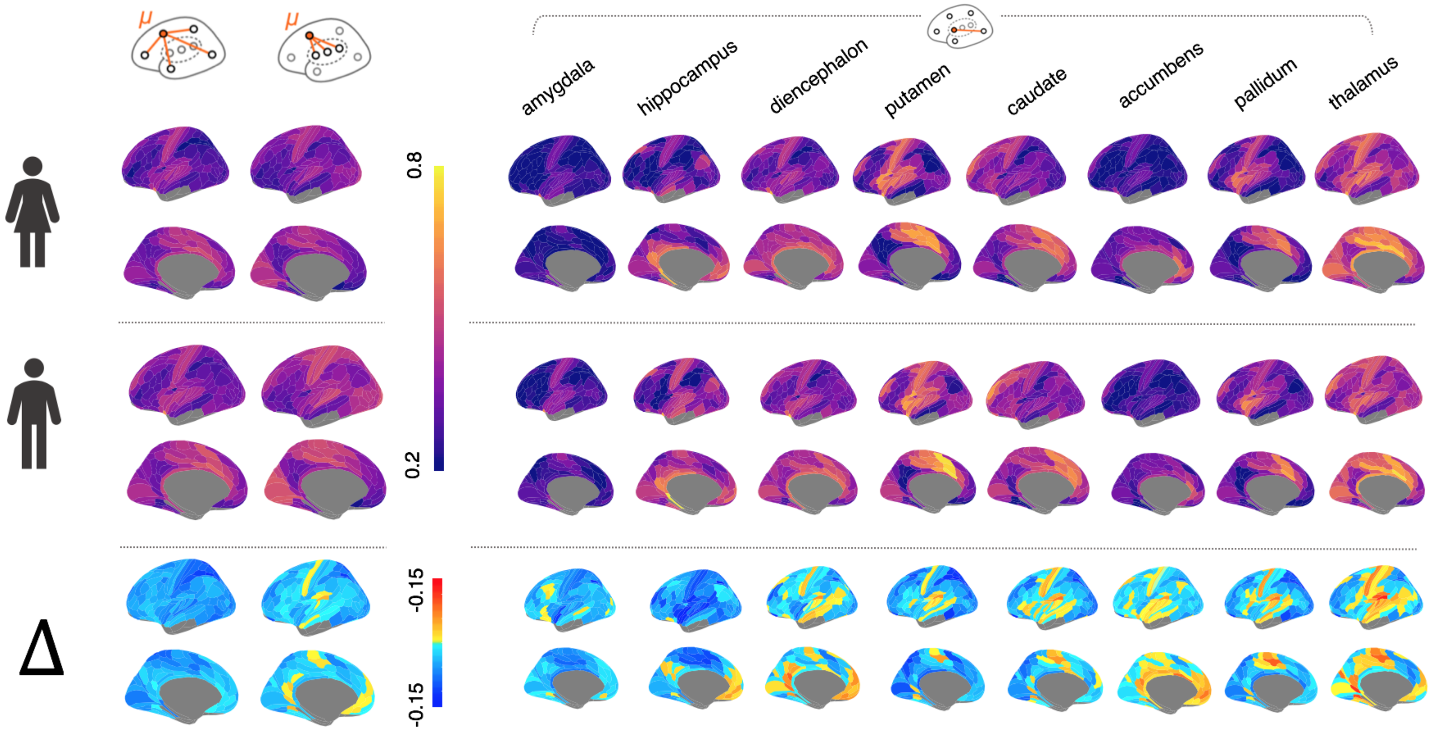

**Fig. S8: All baseline connectivity $FC_{14}$ plots:** Separate linear mixed effects models were fit for both sexes to model regional functional connectivity development as predicted by age and site for cortical-cortical, cortical-subcortical and subcortical-cortical connections. Predicted adolescent baseline connectivity at baseline, $FC_{14}$, was estimated using these models for males and females and a difference map was constructed.

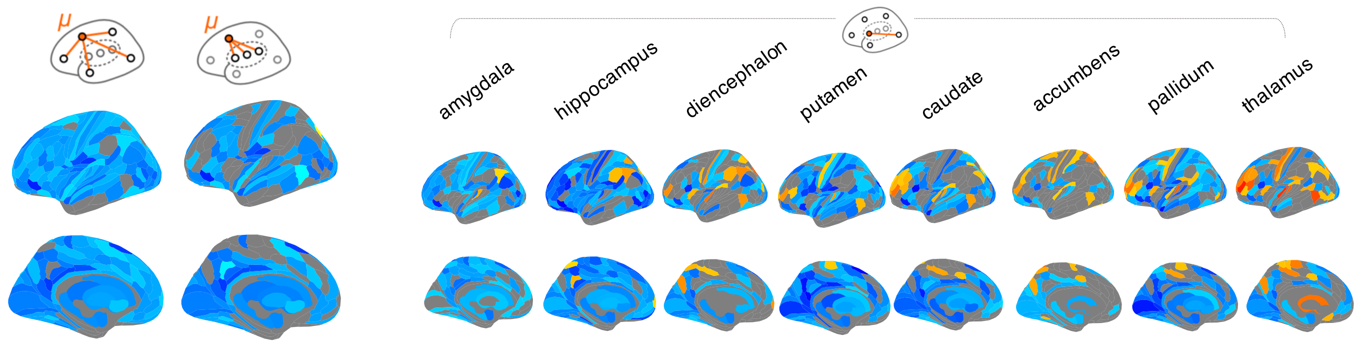

**Fig. S9: Significance of sex difference in baseline connectivity $FC_{14}$:** Representing all plots with more than two significant ROIs.

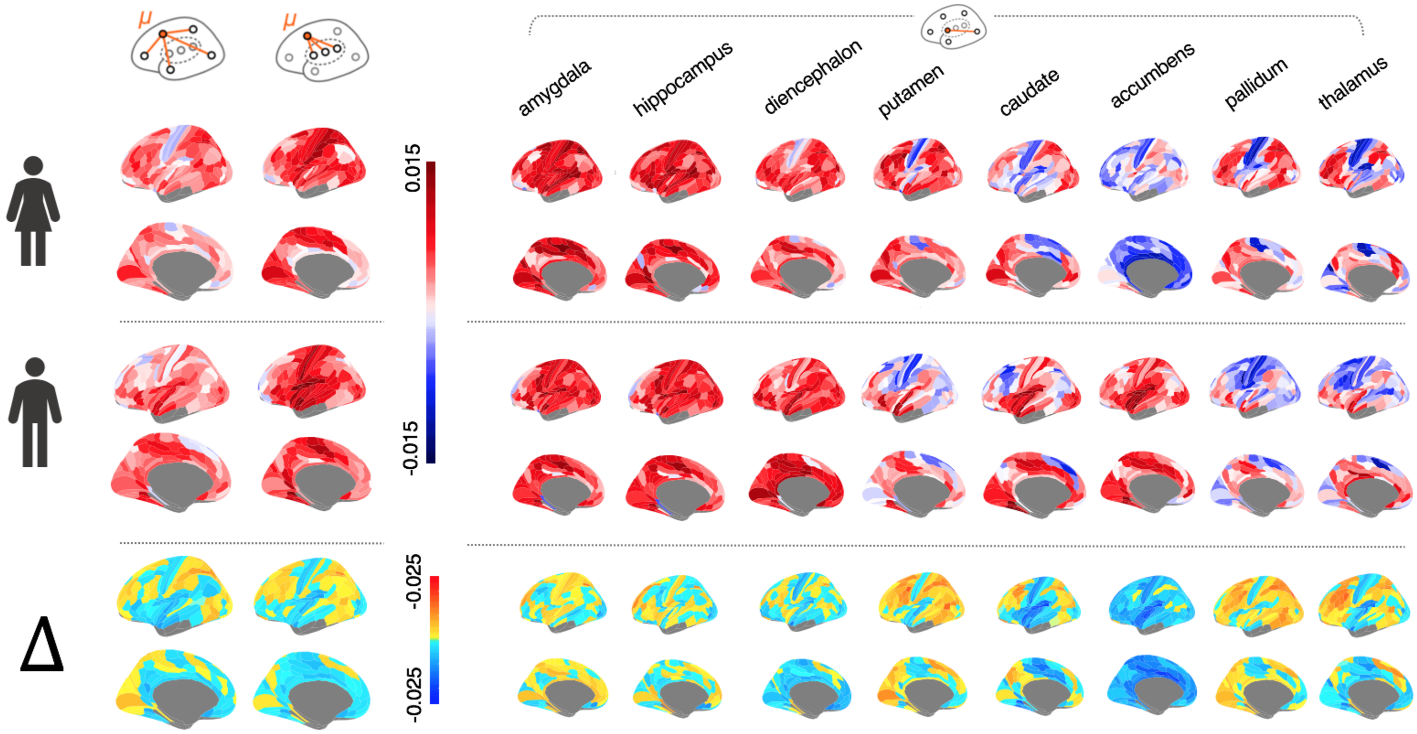

**Fig. S10: All adolescent rate of change $FC_{14-26}$ plots:** The adolescent rate of change, $FC_{14-26}$, was derived as the $\beta_{age}$ coefficient from the linear mixed effects model.

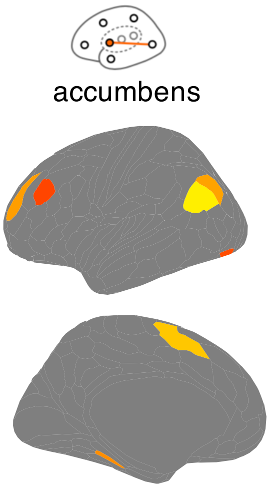

**Fig. S11: Significance of sex difference in adolescent rate of change $\Delta FC_{14-26}$:** Representing all plots with more than two significant ROIs.

| | female | male |
|---|---|---|
| uncorrected all | 93 | 83 |
| fdr-corrected all | 0 | 12 |
| uncorrected cortico-cortical | 101 | 87 |
| fdr-corrected cortico-cortical | 0 | 12 |
| uncorrected cortico-subcortical | 10 | 13 |
| fdr-corrected cortico-subccortical | 0 | 0 |
| uncorrected thalamus | 14 | 4 |
| fdr-corrected thalamus | 5 | 26 |
| uncorrected caudate | 26 | 11 |
| fdr-corrected caudate | 10 | 1 |
| uncorrected putamen | 27 | 30 |
| fdr-corrected putamen | 61 | 71 |
| uncorrected pallidum | 20 | 61 |
| fdr-corrected pallidum | 15 | 35 |
| uncorrected hippocampus | 0 | 0 |
| fdr-corrected hippocampus | 0 | 0 |
| uncorrected amygdala | 0 | 0 |
| fdr-corrected amygdala | 0 | 0 |
| uncorrected accumbens | 0 | 0 |
| fdr-corrected accumbens | 0 | 0 |
| uncorrected diencephalon | 0 | 0 |
| fdr-corrected diencephalon | 0 | 0 |

**Table S2 Age effects on FC per sex:** For each sex separately, we modelled the linear effect of age on functional connectivity for all, cortico-cortical, cortico-subcortical and subcortico-cortical connectivity. We extracted the significance value of the coefficient of age from the model. Here, we calculated the sum of regions that showed a significant effect of age ($p_{\beta_{age}} < 0.05$), uncorrected and FDR-corrected respectively.

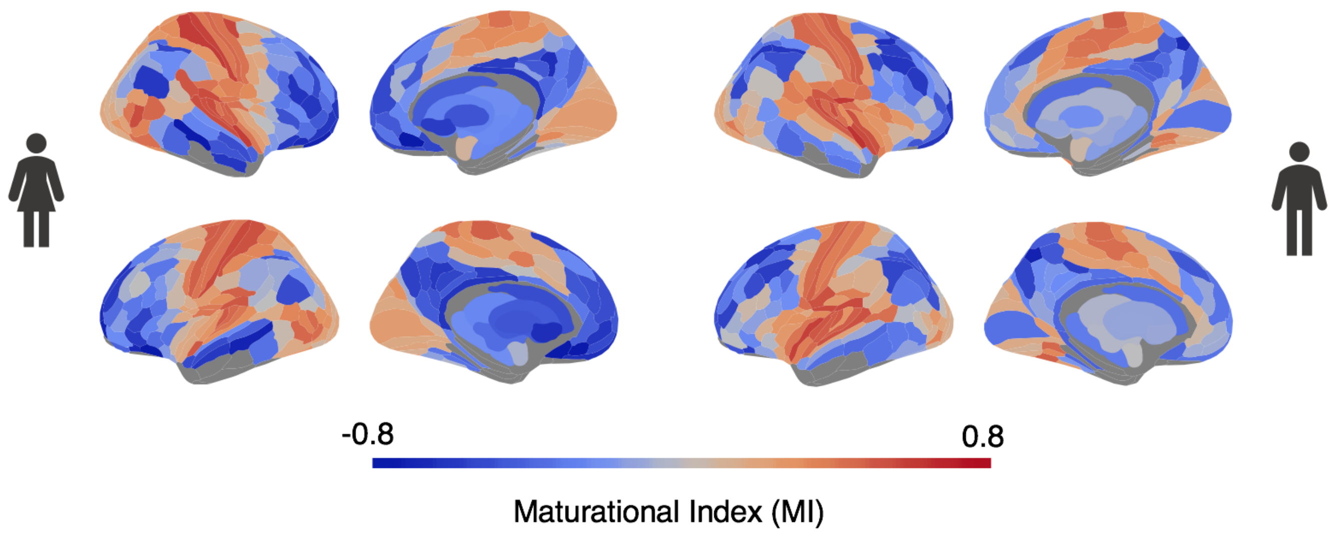

**Fig. S12: All maturational index plots:** Maturational index for males and females separately, all sides of the brain displayed.

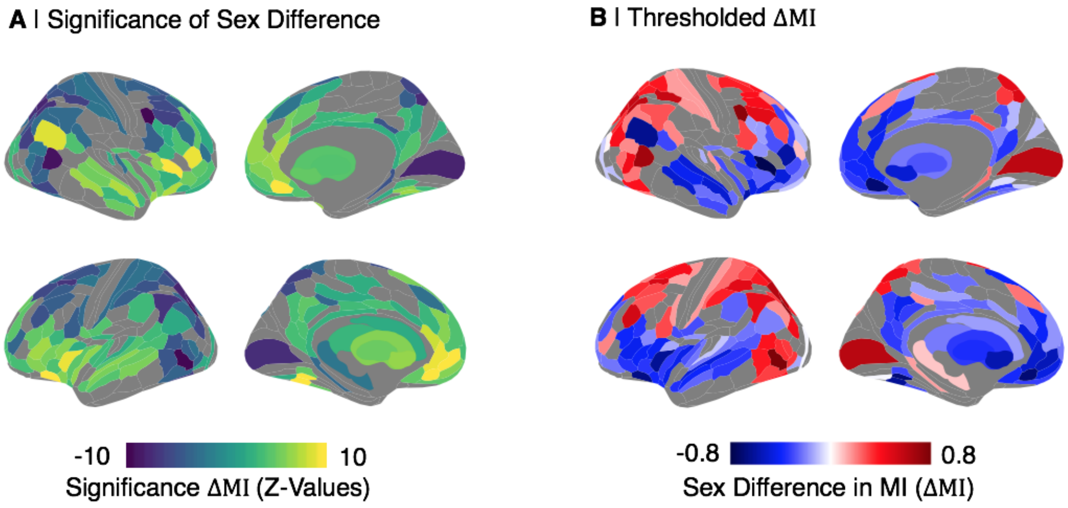

**Fig. S13: Significance of Sex Difference in MI:** We then estimated the significance of this sex difference in a parametric approach (14), by recalculating $MI$ for each sex as a linear regression of edge-wise $FC_{14}$ on $FC_{14-26}$ and testing for the equivalence of the slopes, using their standard errors (SE). (A) 230 ROIs displayed significantly sex divergent behaviour ($P(\Delta MI = 0) < 0.05$). (B) $\Delta MI$ thresholded by significance ($P(\Delta MI = 0) < 0.05$).

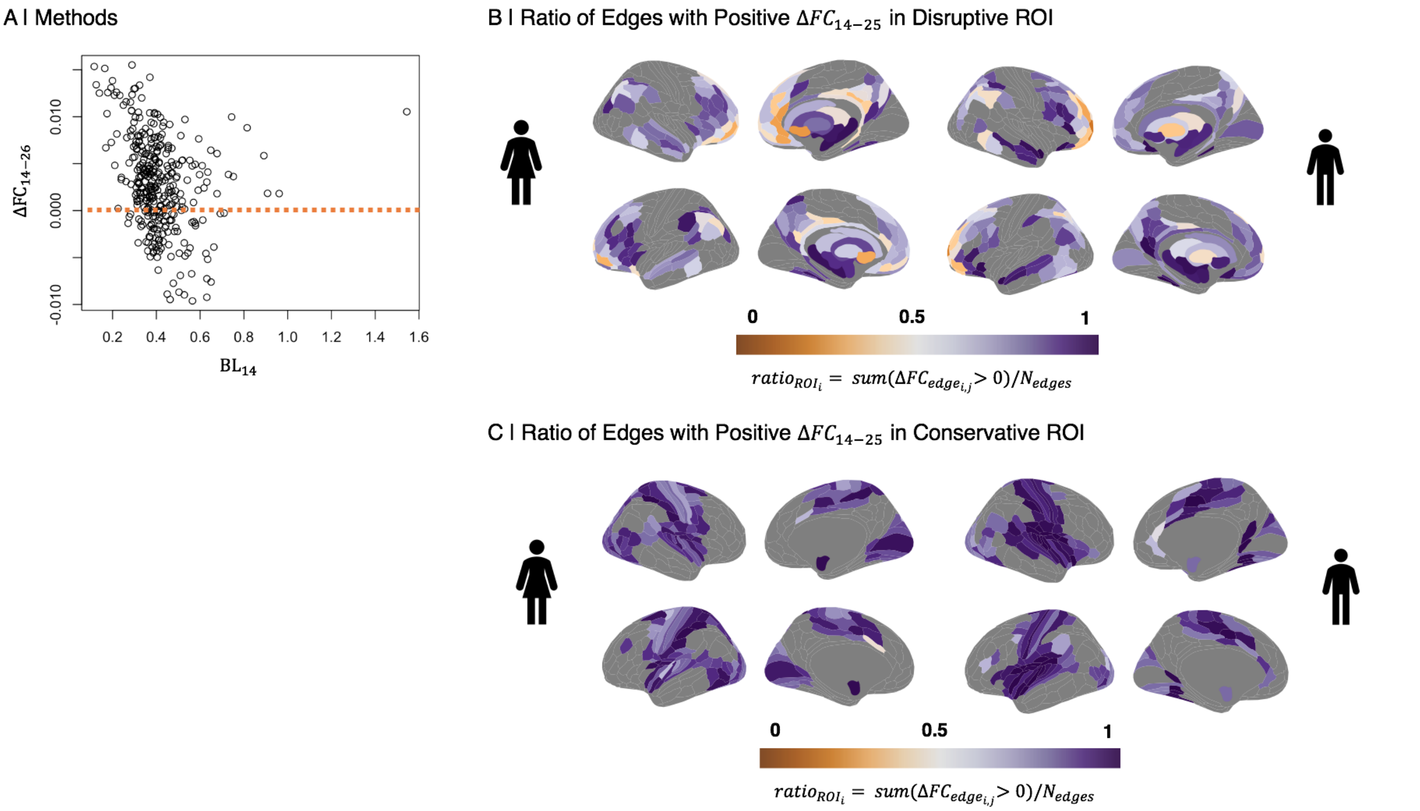

**Fig. S14: Trends in disruptive and conservative development of connectivity:** Disruptive development in a region can mean one of two trends: Either a region is strong at baseline and loses strength over the course of adolescence ('strong getting weaker'), or it is weak at baseline and gains strength ('weak getting stronger'). Conservative development in turn, can mean that either a region is strong at baseline and gains strength over the course of adolescence ('strong getting stronger'), or it is weak at baseline and looses strength ('weak getting weaker'). (A) Here, we estimate these trends for regions of disruptive and conservative change in each sex by calculating the ratio of edges with a positive adolescent rate of change connected to a node ($ratio_{ROI_i} = sum(\Delta FC_{edge_{ij}} > 0)/N_{edges}$). We then thresholded this ratio map for disruptive and conservative nodes in each sex. (B) In disruptive regions, if this $ratio_{ROI_i} > 0.5$, a region is 'strong getting weaker', if $ratio_{ROI_i} < 0.5$, it is 'weak getting stronger'. We find that disruptive regions are predominantly characterized by 'weak getting stronger' changes (78.5% of ROIs in females and 81.3% in males). (C) In conservative regions, if this $ratio_{ROI_i} > 0.5$, a region is 'strong getting strong', if $ratio_{ROI_i} < 0.5$, it is 'weak getting weaker'. We find that all regions in both sexes display 'strong getting stronger' trends only.

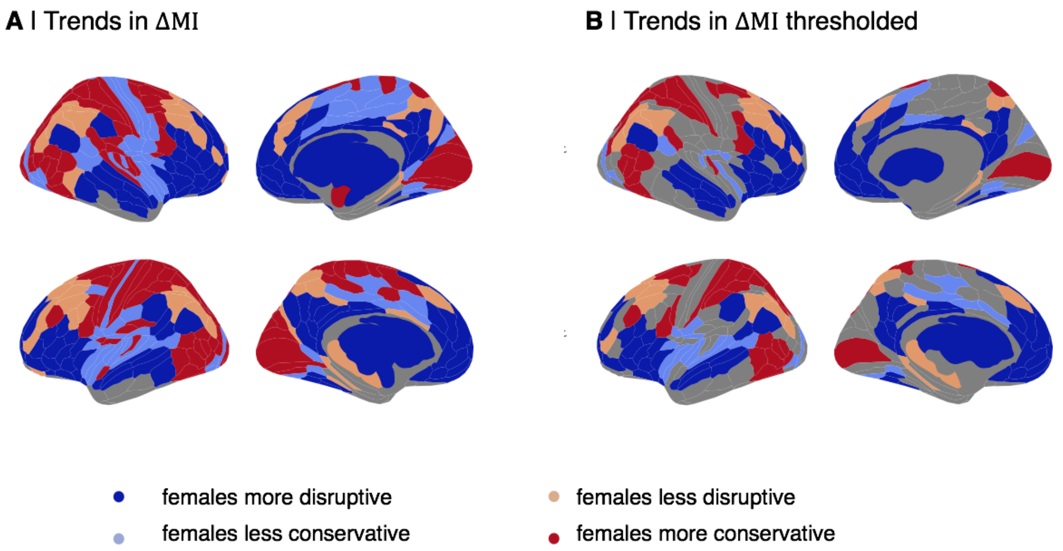

**Fig. S15: Trends in negative $\Delta MI$:** A negative $\Delta MI$ in a region can mean one of two trends: females show (1) more disruptive, or (2) less conservative development than males. Similarly, a positive $\Delta MI$ in a region can mean one of two trends: females show (1) more conservative, or (2) less disruptive development than males. Here, we disentangle these trends for all regions. (A) Trends in $\Delta MI$, unthresholded. (B) Trends in $\Delta MI$, thresholded by significance of sex difference in $\Delta MI$ ($P(\Delta MI = 0) < 0.05$).

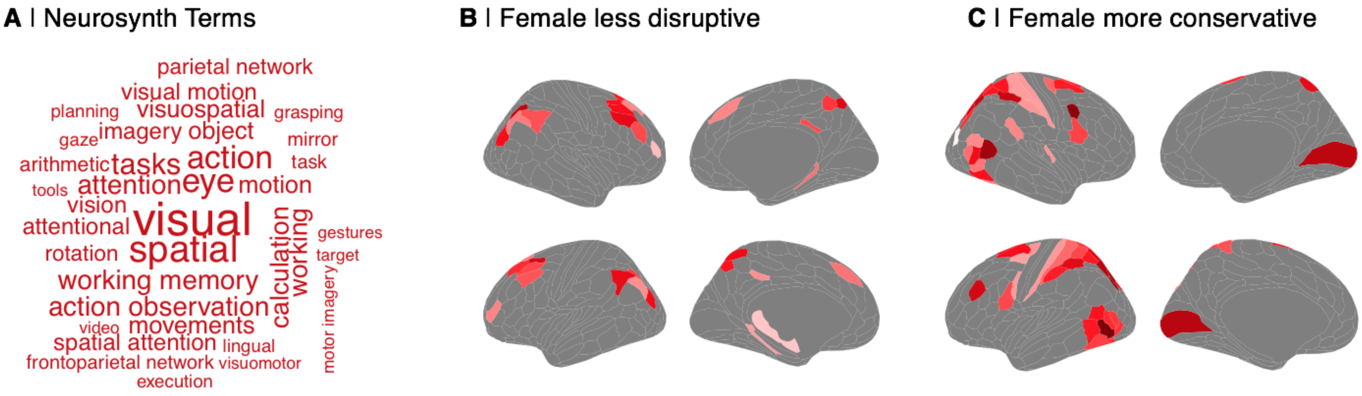

**Fig. S16: Positive $\Delta MI$:** As in our main analysis of regions displaying a negative $\Delta MI$, here we analysed regions with female > male conservative development indicated by positive $\Delta MI$. (A) Neurosynth terms associated with regions positive of positive $\Delta MI$. We uploaded the map of positive $\Delta MI$ regions to Neurosynth and found they are located in cortical areas that were activated by motor and sensory tasks. We further analysed (B) regions with a significant sex difference in $\Delta MI$ ($P(\Delta MI = 0) < 0.05$) which displayed female less disruptive development and (C) regions with a significant sex difference in $\Delta MI$ ($P(\Delta MI = 0) < 0.05$) which displayed female more conservative development. We find those regions are primarily located in association cortical areas.

| ROI | $\Delta MI$ | Z-Value | P-Value | Trend | Yeo-Network |
|---|---|---|---|---|---|
| R FOP5 | -0.79 | 10.57 | 0.00 | female more disruptive | Ventral Attention |
| R s32 | -0.71 | 11.70 | 0.00 | female more disruptive | Default Mode |
| L s32 | -0.71 | 12.05 | 0.00 | female more disruptive | Default Mode |
| L VMV2 | -0.71 | 11.59 | 0.00 | female more disruptive | Visual |
| L 47s | -0.67 | 12.00 | 0.00 | female more disruptive | Default Mode |
| R PGi | -0.64 | 9.04 | 0.00 | female more disruptive | Default Mode |
| L MI | -0.63 | 9.42 | 0.00 | female more disruptive | Ventral Attention |
| L VMV3 | -0.63 | 13.16 | 0.00 | female more disruptive | Visual |
| R IFSa | -0.60 | 9.39 | 0.00 | female more disruptive | Frontoparietal |
| R Pir | -0.59 | 6.98 | 0.00 | female more disruptive | Subcortex |
| L accumbens | -0.57 | 6.79 | 0.00 | female more disruptive | Subcortex |
| L p32 | -0.56 | 10.30 | 0.00 | female more disruptive | Default Mode |
| L a24 | -0.50 | 9.44 | 0.00 | female more disruptive | Default Mode |
| R STSvp | -0.50 | 7.07 | 0.00 | female more disruptive | Default Mode |
| R STSda | -0.49 | 7.62 | 0.00 | female more disruptive | Default Mode |
| R accumbens | -0.49 | 5.13 | 0.00 | female more disruptive | Subcortex |
| R A5 | -0.48 | 5.98 | 0.00 | female more disruptive | Somatomotor |
| L A5 | -0.47 | 6.47 | 0.00 | female more disruptive | Somatomotor |
| R 47l | -0.47 | 5.54 | 0.00 | female more disruptive | Default Mode |
| L p24 | -0.46 | 8.45 | 0.00 | female more disruptive | Default Mode |
| L 10r | -0.46 | 6.29 | 0.00 | female more disruptive | Default Mode |
| R 47s | -0.45 | 7.63 | 0.00 | female more disruptive | Default Mode |
| R FOP4 | -0.45 | 6.86 | 0.00 | female more disruptive | Ventral Attention |
| R a32pr | -0.44 | 5.85 | 0.00 | female more disruptive | Frontoparietal |
| L PoI1 | -0.44 | 6.61 | 0.00 | female less conservative | Ventral Attention |
| R AAIC | -0.44 | 5.37 | 0.00 | female more disruptive | Default Mode |
| L 45 | -0.43 | 7.17 | 0.00 | female more disruptive | Default Mode |
| L 31pd | -0.43 | 4.97 | 0.00 | female more disruptive | Default Mode |
| R STSdp | -0.43 | 5.61 | 0.00 | female more disruptive | Default Mode |
| R p24 | -0.42 | 7.54 | 0.00 | female more disruptive | Default Mode |
| L FOP4 | -0.41 | 6.32 | 0.00 | female more disruptive | Ventral Attention |
| L FOP5 | -0.41 | 5.78 | 0.00 | female more disruptive | Ventral Attention |
| L d32 | -0.40 | 6.34 | 0.00 | female more disruptive | Default Mode |
| R a24 | -0.40 | 7.02 | 0.00 | female more disruptive | Default Mode |
| L SFL | -0.40 | 5.93 | 0.00 | female more disruptive | Frontoparietal |
| L IFSa | -0.40 | 5.10 | 0.00 | female more disruptive | Frontoparietal |
| L STSda | -0.40 | 5.90 | 0.00 | female more disruptive | Default Mode |
| L AAIC | -0.39 | 6.20 | 0.00 | female more disruptive | Default Mode |
| R p32pr | -0.38 | 5.08 | 0.00 | female more disruptive | Ventral Attention |
| L 8BL | -0.38 | 5.32 | 0.00 | female more disruptive | Default Mode |
| R p32 | -0.38 | 6.19 | 0.00 | female more disruptive | Default Mode |
| L Pir | -0.38 | 3.81 | 0.00 | female more disruptive | Subcortex |
| L 25 | -0.38 | 5.11 | 0.00 | female more disruptive | Limbic |
| L pallidum | -0.37 | 5.35 | 0.00 | female more disruptive | Subcortex |
| R 9m | -0.36 | 6.81 | 0.00 | female more disruptive | Default Mode |
| L 9p | -0.36 | 5.32 | 0.00 | female more disruptive | Default Mode |
| L STGa | -0.36 | 4.59 | 0.00 | female more disruptive | Default Mode |
| R 8BL | -0.36 | 3.76 | 0.00 | female more disruptive | Default Mode |
| R POS1 | -0.35 | 4.39 | 0.00 | female more disruptive | Default Mode |
| L IFSp | -0.35 | 5.05 | 0.00 | female more disruptive | Frontoparietal |
| L 47l | -0.35 | 6.28 | 0.00 | female more disruptive | Default Mode |
| L AVI | -0.34 | 4.25 | 0.00 | female more disruptive | Frontoparietal |
| L STSdp | -0.34 | 5.28 | 0.00 | female more disruptive | Default Mode |
| R 10r | -0.34 | 4.50 | 0.00 | female more disruptive | Default Mode |
| L STSvp | -0.34 | 5.94 | 0.00 | female more disruptive | Default Mode |
| L putamen | -0.34 | 6.28 | 0.00 | female more disruptive | Subcortex |

Continued on next page

| ROI | $\Delta MI$ | Z-Value | P-Value | Trend | Yeo-Network |
|---|---|---|---|---|---|
| L POS1 | -0.33 | 3.94 | 0.00 | female more disruptive | Default Mode |
| L PoI2 | -0.33 | 4.84 | 0.00 | female less conservative | Ventral Attention |
| R VMV2 | -0.33 | 6.42 | 0.00 | female less conservative | Visual |
| L 31pv | -0.33 | 3.30 | 0.00 | female more disruptive | Default Mode |
| L 7m | -0.32 | 3.98 | 0.00 | female more disruptive | Default Mode |
| L 10v | -0.32 | 5.02 | 0.00 | female more disruptive | Limbic |
| R 46 | -0.32 | 4.93 | 0.00 | female more disruptive | Frontoparietal |
| L v23ab | -0.31 | 4.14 | 0.00 | female more disruptive | Default Mode |
| L VMV1 | -0.31 | 5.16 | 0.00 | female less conservative | Visual |
| R 25 | -0.31 | 4.42 | 0.00 | female more disruptive | Limbic |
| R 9p | -0.31 | 3.09 | 0.00 | female more disruptive | Default Mode |
| R STV | -0.31 | 4.27 | 0.00 | female less conservative | Default Mode |
| L 43 | -0.30 | 4.87 | 0.00 | female less conservative | Somatomotor |
| R TE1a | -0.30 | 5.33 | 0.00 | female more disruptive | Default Mode |
| R PoI1 | -0.30 | 4.72 | 0.00 | female less conservative | Ventral Attention |
| L OP4 | -0.30 | 4.45 | 0.00 | female less conservative | Somatomotor |
| R p47r | -0.29 | 4.87 | 0.00 | female more disruptive | Frontoparietal |
| L VVC | -0.29 | 5.25 | 0.00 | female more disruptive | Visual |
| R 23d | -0.29 | 4.01 | 0.00 | female more disruptive | Default Mode |
| R MI | -0.29 | 4.74 | 0.00 | female more disruptive | Ventral Attention |
| R PHA2 | -0.29 | 2.47 | 0.02 | female more disruptive | Visual |
| L 52 | -0.29 | 3.54 | 0.00 | female less conservative | Somatomotor |
| R IFSp | -0.29 | 4.16 | 0.00 | female more disruptive | Frontoparietal |
| R 31pd | -0.29 | 3.57 | 0.00 | female more disruptive | Default Mode |
| R pallidum | -0.28 | 4.42 | 0.00 | female more disruptive | Subcortex |
| L 9-46d | -0.28 | 4.79 | 0.00 | female more disruptive | Frontoparietal |
| R 10d | -0.28 | 4.24 | 0.00 | female more disruptive | Default Mode |
| L TPOJ1 | -0.27 | 3.99 | 0.00 | female less conservative | Ventral Attention |
| L PF | -0.26 | 3.52 | 0.00 | female more disruptive | Ventral Attention |
| R AVI | -0.25 | 2.90 | 0.01 | female more disruptive | Frontoparietal |
| L STSva | -0.25 | 4.51 | 0.00 | female more disruptive | Default Mode |
| L p24pr | -0.25 | 4.04 | 0.00 | female less conservative | Ventral Attention |
| R RSC | -0.25 | 2.60 | 0.02 | female more disruptive | Default Mode |
| R 10v | -0.25 | 3.24 | 0.00 | female more disruptive | Limbic |
| R VMV3 | -0.24 | 7.13 | 0.00 | female more disruptive | Visual |
| R STGa | -0.24 | 3.65 | 0.00 | female more disruptive | Default Mode |
| L 23d | -0.24 | 3.65 | 0.00 | female more disruptive | Default Mode |
| L a32pr | -0.24 | 3.25 | 0.00 | female more disruptive | Frontoparietal |
| L RSC | -0.24 | 2.87 | 0.01 | female more disruptive | Default Mode |
| L 24dd | -0.23 | 2.57 | 0.02 | female less conservative | Somatomotor |
| R 33pr | -0.23 | 3.22 | 0.00 | female more disruptive | Ventral Attention |
| R STSva | -0.23 | 3.33 | 0.00 | female more disruptive | Default Mode |
| R VVC | -0.23 | 4.14 | 0.00 | female more disruptive | Visual |
| L STV | -0.23 | 3.54 | 0.00 | female more disruptive | Default Mode |
| R OP2-3 | -0.23 | 4.41 | 0.00 | female less conservative | Somatomotor |
| L a47r | -0.23 | 3.93 | 0.00 | female more disruptive | Frontoparietal |
| L a10p | -0.22 | 4.17 | 0.00 | female more disruptive | Frontoparietal |
| L 10d | -0.22 | 3.86 | 0.00 | female more disruptive | Default Mode |
| L MBelt | -0.22 | 3.54 | 0.00 | female less conservative | Somatomotor |
| L A4 | -0.22 | 4.45 | 0.00 | female less conservative | Somatomotor |
| R putamen | -0.22 | 4.63 | 0.00 | female more disruptive | Subcortex |
| L thalamus | -0.22 | 3.16 | 0.00 | female more disruptive | Subcortex |
| R 47m | -0.21 | 2.78 | 0.01 | female more disruptive | Default Mode |
| R a47r | -0.21 | 4.18 | 0.00 | female more disruptive | Frontoparietal |
| L PGi | -0.21 | 2.17 | 0.05 | female more disruptive | Default Mode |
| R v23ab | -0.21 | 2.79 | 0.01 | female more disruptive | Default Mode |

Continued on next page

| ROI | $\Delta MI$ | Z-Value | P-Value | Trend | Yeo-Network |
|---|---|---|---|---|---|
| R FOP2 | -0.19 | 3.39 | 0.00 | female less conservative | Somatomotor |
| L 44 | -0.19 | 3.25 | 0.00 | female more disruptive | Frontoparietal |
| L p10p | -0.18 | 3.87 | 0.00 | female more disruptive | Frontoparietal |
| R 44 | -0.18 | 2.82 | 0.01 | female more disruptive | Frontoparietal |
| R p24pr | -0.18 | 3.27 | 0.00 | female less conservative | Ventral Attention |
| L OP2-3 | -0.17 | 2.90 | 0.01 | female less conservative | Somatomotor |
| L 9m | -0.17 | 4.29 | 0.00 | female more disruptive | Default Mode |
| L a24pr | -0.17 | 2.66 | 0.01 | female less conservative | Ventral Attention |
| R V8 | -0.16 | 4.44 | 0.00 | female less conservative | Visual |
| R SCEF | -0.16 | 2.42 | 0.02 | female less conservative | Ventral Attention |
| R IFJa | -0.16 | 2.34 | 0.03 | female more disruptive | Frontoparietal |
| L caudate | -0.15 | 2.31 | 0.03 | female more disruptive | Subcortex |
| L 23c | -0.15 | 2.29 | 0.03 | female less conservative | Ventral Attention |
| R IFJp | -0.15 | 2.54 | 0.02 | female more disruptive | Dorsal Attention |
| L LO2 | -0.14 | 2.30 | 0.03 | female less conservative | Visual |
| L FOP3 | -0.14 | 2.38 | 0.03 | female less conservative | Ventral Attention |
| R A1 | -0.13 | 2.43 | 0.02 | female less conservative | Somatomotor |
| R 9-46d | -0.12 | 2.15 | 0.05 | female more disruptive | Frontoparietal |
| R 7m | -0.12 | 2.24 | 0.04 | female more disruptive | Default Mode |
| R V6 | -0.11 | 2.73 | 0.01 | female less conservative | Visual |
| R LO2 | -0.10 | 3.42 | 0.00 | female less conservative | Visual |
| L PBelt | -0.09 | 2.21 | 0.04 | female less conservative | Somatomotor |
| R 11l | -0.09 | 2.70 | 0.01 | female more disruptive | Frontoparietal |
| R V4 | -0.08 | 3.56 | 0.00 | female less conservative | Visual |
| R VMV1 | -0.06 | 3.41 | 0.00 | female less conservative | Visual |
| L V4 | -0.01 | 2.50 | 0.02 | female less conservative | Visual |
| R V3CD | 0.02 | 3.42 | 0.00 | female more conservative | Visual |
| L hippocampus | 0.09 | -2.20 | 0.04 | female less disruptive | Subcortex |
| R a9-46v | 0.10 | -2.46 | 0.02 | female less disruptive | Frontoparietal |
| L FEF | 0.15 | -2.14 | 0.05 | female more conservative | Dorsal Attention |
| L PreS | 0.15 | -3.29 | 0.00 | female less disruptive | Visual |
| R 2 | 0.15 | -2.35 | 0.03 | female more conservative | Somatomotor |
| L 6v | 0.16 | -3.16 | 0.00 | female more conservative | Somatomotor |
| L 1 | 0.16 | -3.09 | 0.00 | female more conservative | Somatomotor |
| R 1 | 0.16 | -3.04 | 0.00 | female more conservative | Somatomotor |
| R PFt | 0.16 | -2.30 | 0.03 | female more conservative | Dorsal Attention |
| L 6d | 0.17 | -2.54 | 0.02 | female more conservative | Somatomotor |
| R 52 | 0.17 | -2.84 | 0.01 | female more conservative | Somatomotor |
| L 31a | 0.18 | -3.39 | 0.00 | female less disruptive | Frontoparietal |
| L PGs | 0.18 | -3.36 | 0.00 | female less disruptive | Default Mode |
| R 8Ad | 0.18 | -3.62 | 0.00 | female less disruptive | Default Mode |
| R TPOJ3 | 0.19 | -2.21 | 0.04 | female more conservative | Dorsal Attention |
| R 8BM | 0.19 | -2.23 | 0.04 | female less disruptive | Frontoparietal |
| R PSL | 0.19 | -3.14 | 0.00 | female more conservative | Default Mode |
| L a9-46v | 0.19 | -3.58 | 0.00 | female less disruptive | Frontoparietal |
| L V6A | 0.20 | -2.60 | 0.02 | female more conservative | Visual |
| R PGs | 0.21 | -3.24 | 0.00 | female less disruptive | Default Mode |
| L s6-8 | 0.21 | -3.06 | 0.00 | female less disruptive | Frontoparietal |
| R FFC | 0.21 | -2.95 | 0.01 | female more conservative | Visual |
| R FST | 0.22 | -3.82 | 0.00 | female more conservative | Dorsal Attention |
| L 8BM | 0.22 | -2.66 | 0.01 | female less disruptive | Frontoparietal |
| R PreS | 0.23 | -5.01 | 0.00 | female less disruptive | Visual |
| R MT | 0.23 | -2.60 | 0.02 | female more conservative | Visual |
| L 2 | 0.23 | -2.48 | 0.02 | female more conservative | Somatomotor |
| R 6d | 0.24 | -3.44 | 0.00 | female more conservative | Somatomotor |
| L 8C | 0.27 | -4.09 | 0.00 | female less disruptive | Frontoparietal |

Continued on next page

| ROI | $\Delta MI$ | Z-Value | P-Value | Trend | Yeo-Network |
|---|---|---|---|---|---|
| R PFm | 0.27 | -3.22 | 0.00 | female less disruptive | Frontoparietal |
| L 5L | 0.28 | -3.46 | 0.00 | female more conservative | Somatomotor |
| R d23ab | 0.28 | -4.89 | 0.00 | female less disruptive | Default Mode |
| R p9-46v | 0.29 | -2.83 | 0.01 | female less disruptive | Frontoparietal |
| L 6r | 0.29 | -4.24 | 0.00 | female more conservative | Ventral Attention |
| L 8Av | 0.29 | -4.25 | 0.00 | female less disruptive | Frontoparietal |
| L 7PC | 0.29 | -3.00 | 0.00 | female more conservative | Dorsal Attention |
| R 6r | 0.30 | -4.07 | 0.00 | female more conservative | Ventral Attention |
| L PH | 0.31 | -3.70 | 0.00 | female more conservative | Dorsal Attention |
| L 7AL | 0.31 | -2.80 | 0.01 | female more conservative | Dorsal Attention |
| L IP1 | 0.32 | -3.47 | 0.00 | female less disruptive | Frontoparietal |
| R PCV | 0.32 | -4.11 | 0.00 | female less disruptive | Default Mode |
| L MT | 0.32 | -2.67 | 0.01 | female more conservative | Visual |
| L V4t | 0.32 | -3.46 | 0.00 | female more conservative | Visual |
| R 6ma | 0.34 | -4.23 | 0.00 | female more conservative | Ventral Attention |
| L MST | 0.34 | -4.28 | 0.00 | female more conservative | Visual |
| R IP0 | 0.34 | -4.71 | 0.00 | female more conservative | Dorsal Attention |
| L 8Ad | 0.34 | -5.79 | 0.00 | female less disruptive | Default Mode |
| R VIP | 0.36 | -4.71 | 0.00 | female more conservative | Dorsal Attention |
| R V7 | 0.36 | -3.02 | 0.00 | female more conservative | Visual |
| L PFt | 0.36 | -4.11 | 0.00 | female more conservative | Dorsal Attention |
| R 6a | 0.37 | -3.96 | 0.00 | female more conservative | Dorsal Attention |
| L PHT | 0.38 | -4.75 | 0.00 | female more conservative | Dorsal Attention |
| R LIPv | 0.38 | -5.10 | 0.00 | female more conservative | Dorsal Attention |
| L LIPv | 0.38 | -4.90 | 0.00 | female more conservative | Dorsal Attention |
| R 7PC | 0.38 | -3.75 | 0.00 | female more conservative | Dorsal Attention |
| R s6-8 | 0.39 | -4.74 | 0.00 | female less disruptive | Frontoparietal |
| L 7Pm | 0.39 | -5.65 | 0.00 | female less disruptive | Frontoparietal |
| L TPOJ2 | 0.39 | -4.71 | 0.00 | female more conservative | Dorsal Attention |
| R PH | 0.39 | -5.95 | 0.00 | female more conservative | Dorsal Attention |
| R i6-8 | 0.40 | -5.28 | 0.00 | female less disruptive | Frontoparietal |
| L FFC | 0.40 | -6.81 | 0.00 | female more conservative | Visual |
| R PGp | 0.41 | -6.37 | 0.00 | female less disruptive | Dorsal Attention |
| R 8Av | 0.41 | -5.67 | 0.00 | female less disruptive | Frontoparietal |
| L 6a | 0.42 | -4.92 | 0.00 | female more conservative | Dorsal Attention |
| L V7 | 0.42 | -3.74 | 0.00 | female more conservative | Visual |
| R MST | 0.43 | -4.44 | 0.00 | female more conservative | Visual |
| L PGp | 0.43 | -5.79 | 0.00 | female less disruptive | Dorsal Attention |
| L 7Am | 0.44 | -6.34 | 0.00 | female less disruptive | Dorsal Attention |
| L 6ma | 0.44 | -5.45 | 0.00 | female more conservative | Ventral Attention |
| L PFm | 0.46 | -7.15 | 0.00 | female less disruptive | Frontoparietal |
| R 7Am | 0.47 | -6.02 | 0.00 | female more conservative | Dorsal Attention |
| R 8C | 0.48 | -6.24 | 0.00 | female less disruptive | Frontoparietal |
| L AIP | 0.48 | -4.87 | 0.00 | female more conservative | Dorsal Attention |
| L 7PL | 0.49 | -6.72 | 0.00 | female more conservative | Dorsal Attention |
| L VIP | 0.49 | -6.28 | 0.00 | female more conservative | Dorsal Attention |
| R 7Pm | 0.50 | -7.75 | 0.00 | female less disruptive | Frontoparietal |
| L i6-8 | 0.52 | -7.12 | 0.00 | female less disruptive | Frontoparietal |
| R IPS1 | 0.52 | -7.42 | 0.00 | female more conservative | Dorsal Attention |
| L p9-46v | 0.54 | -5.07 | 0.00 | female more conservative | Frontoparietal |
| R 7PL | 0.54 | -7.60 | 0.00 | female more conservative | Dorsal Attention |
| R AIP | 0.57 | -6.30 | 0.00 | female more conservative | Dorsal Attention |
| R MIP | 0.58 | -8.11 | 0.00 | female more conservative | Dorsal Attention |
| R IP1 | 0.59 | -7.40 | 0.00 | female less disruptive | Frontoparietal |
| R V1 | 0.60 | -8.27 | 0.00 | female more conservative | Visual |
| L V1 | 0.62 | -7.60 | 0.00 | female more conservative | Visual |

Continued on next page

| ROI | $\Delta MI$ | Z-Value | P-Value | Trend | Yeo-Network |
|---|---|---|---|---|---|
| L MIP | 0.63 | -8.23 | 0.00 | female more conservative | Dorsal Attention |
| L IPS1 | 0.63 | -8.53 | 0.00 | female more conservative | Dorsal Attention |
| R TPOJ2 | 0.68 | -9.32 | 0.00 | female more conservative | Dorsal Attention |
| R PEF | 0.76 | -10.03 | 0.00 | female more conservative | Dorsal Attention |
| L IP0 | 0.77 | -10.83 | 0.00 | female more conservative | Dorsal Attention |
| L FST | 0.80 | -11.75 | 0.00 | female more conservative | Dorsal Attention |

**Table S3: ROIs with significantly different $\Delta MI$**: We tested for sex differences in maturational index (MI). Here we show all 230 regions displaying a significant sex difference in MI ($P(\Delta MI = 0) < 0.05$). We show the regions name in the HCP parcellation; it's $\Delta MI$ value, the p-value and Z-value from the parametric test of the sex difference in $MI$; the functional network they are located in ('Yeo-Network') (15); as well as which one of four trends they display: (1) *'female more conservative'*, (2) *'female more disruptive'*, (3) *'female less conservative'*, (4) *'female less disruptive'* (cf. Supplementary Fig. S15).

#### Co-location with Depression

*Sample Overview*

The Biomarkers for Depression (BioDep) study is a case-control study of adult subjects, aged 25-50 years, with and without major depressive disorder (MDD). Data was collected from a total of 129 subjects: 46 healthy controls, and 83 MDD patients, as measured in a Structured Clinical Interview for DSM-V Depressive Disorders (SCID), as well as a global Hamilton Rating Scale for Depression (HAM-D) score of higher than 13. The MDD group contained cases with CRP $< 3$ mg/L (N=53), and MDD cases with CRP $> 3$ mg/L (N=34). Here, we are focusing on the low CRP cases, only. The final sample after quality control contained 46 healthy control and 50 MDD patients (cf. Supplementary Table S4).

*MRI Preprocessing*

We used a multi-echo (me) echoplanar imaging (EPI) sequence (16) to collect fMRI data under resting state conditions with the following parameters: relaxation time (TR) = 2.57 s; echo times ($\mathsf{TE}_{1,2,3}$) = 15, 34 and 54 ms; acquisition time = 10 mins 42.5 s = 250 time points in each fMRI time series. meEPI data were collected as 32 slices at -30 degrees to the AC-PC line, field of view: 240 mm, matrix size: 64 ×64, voxel resolution: 3.75 ×3.75 ×4 mm.

The first 6 volumes were discarded to ensure scanner equilibrium and the remaining data pre-processed using multi-echo independent component analysis (ME-ICA; (7; 8)) to identify sources of variance in the fMRI time series that scaled linearly with TE and be confidently regarded as BOLD signal. Other non-BOLD sources of variance, such as head movement, that do not scale with TE, were identified by ME-ICA and discarded. The retained independent components, representing BOLD contrast, were optimally recomposed to generate a broadband denoised fMRI time series at each voxel. This was bandpass filtered using the Maximal Overlap Discrete Wavelet Transform ("modwt" using "la8", the Daubechies orthonormal compactly supported wavelet of length L=8), resulting in a BOLD signal oscillating in the frequency range 0.02-0.1 Hz (wavelet scales 2 and 3).

Geometric re-alignment was used to estimate 6 motion parameters for each participant (3 translation and 3 rotation parameters) which were used to calculate an overall estimate of motion - framewise displacement (FD; defined as the Euclidean norm of motion and rotation derivatives in mm: $\mathsf{FD}^2 = |\vec{\Delta x}|^2 + |\vec{\Delta \theta}|^2$). For each participant, mean FD was calculated by averaging the FD time series. A total of 3 scans were excluded due to high in-scanner motion $\langle \mathsf{FD} \rangle_{\mathrm{RMS}} > 0.3$ mm or max(FD) $> 1.3$ mm and one subject was dropped due to excessively high mean correlation $> 0.7$.

Each pre-processed fMRI image was regionally parcellated into the same set of cortical and subcortical regions as the qMT data and the regional mean fMRI time series estimated for each cortical and sub-cortical region using the non-zero mean variant of the AFNI *3dROIstats* command (17). Thus we estimated a 376×244 regional time series matrix for each participant.

The functional connectivity between each regional pair of fMRI time series was estimated by Pearson's correlation coefficient $r$ for each possible pair of regions, resulting in a 376×376 symmetric association or functional connectivity matrix. The row (or column) means of this matrix comprise the vector of regional or nodal weighted degree.

| Group | Sex *female* | $\mu$ Age | $\sigma$ Age | $\mu$ FD | $\sigma$ FD | Centre | | |
|---|---|---|---|---|---|---|---|---|
| | | | | | | Cambridge | Kings | Oxford |
| Control | 27 | 35.5 | 7.5 | 0.08 | 0.05 | 36 | 6 | 4 |
| MDD | 29 | 36.8 | 7.1 | 0.01 | 0.05 | 37 | 8 | 5 |

**Table S4 BioDep Sample Overview:** A total of N=96 subjects (50 MDD patients) subjects, balanced for age and sex were scanned at three MRI imaging centres.

*Case-Control Map*

We constructed a case-control difference map by estimating the effect (t-value) of group (patient vs. control) on region-wise functional connectivity (FC) strength, controlling for sex. We corrected for multiple comparisons ($p < 0.05$, FDR-corrected).

#### Enrichment Analysis

*Partial Least Squares Regression*

Partial least squares regression has been used as a powerful tool to describe the relationship between two sets of variables (represented as two matrices) which uses latent variables to model the covariance structure between the two. Its ability to handle situations with a large number of potentially multicollinear predictors has made it useful for analysis of neuroimaging data (18; 4; 19). Briefly, PLS finds components that explain the maximum covariance between the dependent and independent variables. Here, we use PLS as a means to find the weighted gene expression pattern that is most strongly correlated with the anatomical pattern of sex differences in adolescent functional connectivity maturation, $\Delta MI$. Thus we regressed the 360-length vector of $\Delta MI$ on the 360 by 15,746 matrix of post mortem transcriptomic gene expression data from the Allen Human Brain Atlas collected from 6 donor brains (5 males) (20; 21). We analysed whether the first PLS component explains more variance than expected by chance by randomly permuting the rows of the gene expression matrix and comparing the variance explained by PLS regression of $\Delta MI$ on the observed transcriptional data with the distribution of variance in $\Delta MI$ explained by 1000 random permutations of the brain gene expression matrix. For the first PLS component (PLS1), which accounted for the greatest proportion of variance we estimated the variability of each transcript's weighting coefficient by bootstrap resampling (10,000 times) of the brain regional transcription matrix. The effect size and statistical significance of individual transcript weights on PLS1 were defined by the Z-score (observed coefficient divided by bootstrap standard error).

*Median Rank-Based Gene Enrichment*

We used a median rank-based approach to assess the enrichment of PLS1 on several published gene lists (22). This allows us to assess whether a given gene list is non-randomly represented among the most strongly weighted PLS1 genes that have brain expression anatomically co-located with the spatial pattern of sex differences in maturational index. To do this, each gene on the prior gene list of interest is ranked in terms of its Z-score weighting on PLS1 and the observed median rank is estimated; then a equivalent number of genes, matched for gene length, are randomly selected and their median rank on the PLS1 component is estimated. This second step of randomly selecting and ranking genes is repeated 10,000 times to sample the permutation distribution of median rank. Finally, the null hypothesis that the observed median rank (for the gene list of interest) was not significantly different from the median rank of a random list of genes (matched for gene length and number of genes) was tested by comparing the observed median rank to the centiles of the permutation distribution. For example, for a two-tailed test of significant enrichment with $P < 0.05$, if the observed median rank was lower than the 2.5th percentile of the permutation distribution, then the gene list of interest was significantly enriched among the most negatively weighted PLS1 genes; whereas if the observed median rank was greater than the 97.5th percentile of the permutation distribution then the gene list of interest was significantly enriched among the most positively weighted PLS1 genes.

Finally, statistical significance of observed gene set median ranks was established by comparison with the null median rank distributions from 10,000 gene rank permutations. The direction of the effect is relative to the median rank expected by chance. Thus if a gene set's real median rank is significantly lower than expected by chance the gene set is associated with or enriched for the bottom of PLS1 since that set of genes ranks lower on PLS1 than expected for a random set of genes of similar length; if it is higher, it's genes are enriched towards the top end of the component.

*Matching for Gene Length*

We matched the genes in our empirical gene list for gene length by finding a set of genes in the list of AHBA genes which were close in length by some criterion detailed further below. We then proceeded to resample from that subset of AHBA genes to find a set of genes matched for gene length with the empirical gene lists ($P < 0.05$). We used one of three approaches, in increasing order of strictness applied to find this initial set of genes to resample from. (i) In the 'standard' approach, we matched for gene length by finding the 5 nearest neighbours (Mahalanobis distance) of any given gene from an external gene set in the list of AHBA (PLS1) genes. (ii) Should the algorithm fail to find a set of genes matched for gene length in that way ($P < 0.05$), we restricted the approach to find only the 2 nearest neighbours in the same way ('2 nearest neighbours' approach). (iii) Lastly, should that approach fail, too, we resampled the 3 nearest neighbours without replacement ('without replacement' approach).

*Developmental Enrichment*

We uploaded a ranked list of genes with a significantly ($P_{FDR} < 0.05$) negative or positive PLS1 weight respectively to the cell specific enrichment analysis (CSEA) tool (23) under the category *SEA across brain regions and development*. The CSEA tool uses Human data are from BrainSpan, an atlas of the developing human brain (24), with postmortem human brain specimens collected across 13 developmental stages (4 weeks post conception to 60 years of age) in 8-16 brain structures.

*Prenatal Cell Type Enrichment*

We tested PLS1 for cell type specific enrichment using single-cell transcriptomic gene expression data from mid gestation (gestation week 17 to 18; (25)). This data included 16 unique clusters: endothelial cells (End), excitatory deep layer 1 (ExDp1), excitatory deep layer 2 (ExDp2), maturing excitatory (ExM), newborn excitatory neurons (ExN), intermediate progenitor cells (IP), microglia (Mic), oligodendrocyte precursor cells (OPC), outer radial glia (oRG), pericytes (Per), cycling progenitor G2/M phase (PgG2M), cycling progenitor S phase (PgS), ventricular radial glia (vRG).

*Adult Cell Type Enrichment*

We tested PLS1 for cell type specific enrichment. We used gene expression data of 33 distinct cellular clusters (26), including cortical excitatory (Ex) and inhibitory (In) neurons, cerebellar granule (Gran) cells and Purkinje (Purk) neurons, as well as non-neuronal cells, including endothelial cells (End), smooth muscle cells or pericytes (Per), astrocytes (Ast), oligodendrocytes (Oli), oligodendrocyte precursor cells (OPCs), and microglia (Mic). We excluded cell types expressed in the cerebellum only.

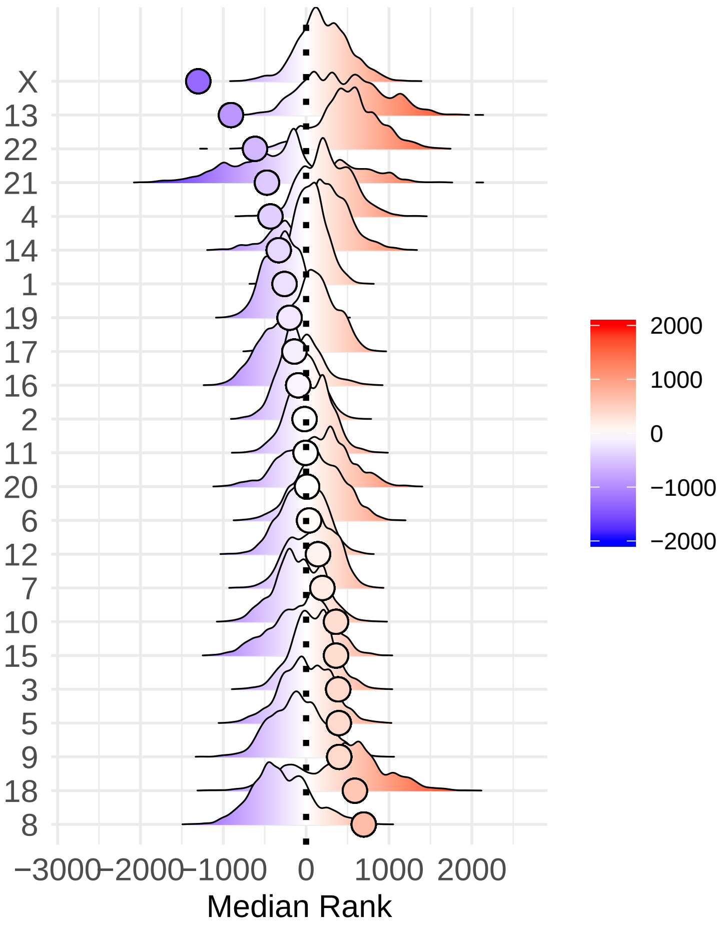

**Fig. S17: Chromosomal null models:** We assessed chromosomal enrichment of the genes on PLS1 to test our hypothesis that sex chromosomal gene expression is related to the sexual differences in adolescent brain development. We used a median gene rank approach to analyse an enrichment of PLS1 for genes located on specific chromosomes. Here, we show the permutation distributions of random gene ranks for each chromosome. We overlay the true median gene rank as a dot on each distribution.

| Chromosome | P | s | Z | N Genes | Approach |
|---|---|---|---|---|---|
| 1 | 0.23 | 0.00 | -0.89 | 5433 | standard |
| 10 | 0.71 | 0.00 | 0.51 | 2513 | standard |
| 11 | 0.14 | 0.00 | -0.85 | 3517 | standard |
| 12 | 0.55 | 0.00 | 0.11 | 3201 | standard |
| 13 | 0.55 | 0.00 | 0.10 | 1165 | standard |
| 14 | 0.42 | 0.00 | -0.08 | 2022 | standard |
| 15 | 0.67 | 0.00 | 0.17 | 1971 | standard |
| 16 | 0.17 | 0.00 | -1.10 | 2761 | standard |
| 17 | 0.05 | 0.00 | -1.45 | 3383 | standard |
| 18 | 0.65 | 0.00 | 0.35 | 970 | standard |
| 19 | 0.04 | 0.00 | -1.56 | 4043 | standard |
| 2 | 0.77 | 0.00 | 0.59 | 3960 | standard |
| 20 | 0.53 | 0.00 | 0.18 | 1884 | standard |
| 21 | 0.07 | 0.00 | -1.35 | 643 | standard |
| 22 | 0.51 | 0.00 | 0.20 | 1552 | standard |
| 3 | 0.93 | 0.00 | 1.41 | 3362 | standard |
| 4 | 0.72 | 0.00 | 0.42 | 2297 | standard |
| 5 | 0.83 | 0.00 | 0.78 | 2787 | standard |
| 6 | 0.88 | 0.00 | 1.19 | 2691 | standard |
| 7 | 0.65 | 0.00 | 0.50 | 2848 | standard |
| 8 | 0.58 | 0.00 | 0.10 | 2202 | standard |
| 9 | 0.39 | 0.00 | -0.36 | 2466 | standard |
| X | 0.45 | 0.00 | -0.23 | 2395 | standard |
| Y | 0.66 | 0.00 | -0.15 | 55 | standard |

**Table S5 Chromosomal Enrichment Null Model Statistics:** We assessed chromosomal enrichment of the genes on PLS1 to test our hypothesis that sex chromosomal gene expression is related to the sexual differences in adolescent brain development. We used a median gene rank approach to analyse an enrichment of PLS1 for genes located on specific chromosomes. We did so matching for gene length. Specifically, we resampled genes from the Allen Human Brain Atlas (AHBA) until we found a subset of genes that were not significantly different in length ($P < 0.05$) from the empirical gene list. Here, we show the results from the gene length matching algorithm, including the*P-,s- and Z-value* for the test of difference in gene length between the empirical gene list and the subset of resampled genes, the number of genes *N Genes*, to resample from, as well as which *approach* the algorithm converged on (see Supplementary Text *"Matching for Gene Length"* for details.)

| cluster | p | s | z | n | approach |
|---|---|---|---|---|---|
| ExN | 0.55 | 0.00 | 0.08 | 364 | standard |
| PgG2M | 0.39 | 0.00 | -0.54 | 1786 | standard |
| OPC | 0.91 | 0.00 | 1.47 | 2125 | standard |
| End | 0.81 | 0.00 | 0.84 | 2393 | standard |
| ExDp2 | 0.96 | 0.00 | 1.87 | 1207 | 2 nearest neighbours |
| Per | 0.48 | 0.00 | -0.12 | 3388 | standard |
| Mic | 0.57 | 0.00 | 0.34 | 2982 | standard |
| ExM | 0.77 | 0.00 | 0.52 | 1006 | standard |
| IP | 0.41 | 0.00 | -0.03 | 1100 | standard |
| ExDp1 | 0.94 | 0.00 | 1.32 | 865 | 2 nearest neighbours |
| ExCal | 0.89 | 0.00 | 1.28 | 1001 | standard |
| InSST | 0.57 | 0.00 | 0.25 | 748 | standard |
| InCALB2 | 0.60 | 0.00 | 0.42 | 683 | standard |
| oRG | 0.78 | 0.00 | 0.90 | 2482 | standard |
| PgS | 0.42 | 0.00 | -0.10 | 2200 | standard |
| vRG | 0.48 | 0.00 | 0.01 | 1925 | standard |

**Table S6 Prenatal Enrichment Null Model Statistics:** We assessed prenatal cell specific enrichment of the genes on PLS1 (25). We used a median gene rank approach to analyse an enrichment of PLS1 for genes associated with specific cell clusters. We did so matching for gene length. Specifically, we resampled genes from the Allen Human Brain Atlas (AHBA) until we found a subset of genes that were not significantly different in length ($P < 0.05$) from the empirical gene list. Here, we show the results from the gene length matching algorithm, including the*P-,s- and Z-value* for the test of difference in gene length between the empirical gene list and the subset of resampled genes, the number of genes *N Genes*, to resample from, as well as which *approach* the algorithm converged on (see Supplementary Text *"Matching for Gene Length"* for details.)

| cluster | p | s | z | n | approach |
|---|---|---|---|---|---|
| Ex1 | 0.89 | 0.00 | 1.14 | 497 | 2 nearest neighbours |
| Ex2 | 0.95 | 0.00 | 1.53 | 320 | 2 nearest neighbours |
| Ex3a | 0.59 | 0.00 | 0.38 | 738 | without replacement |
| Ex3b | 0.81 | 0.00 | 1.04 | 1851 | standard |
| Ex3c | 0.50 | 0.00 | 0.12 | 1113 | standard |
| Ex3d | 0.94 | 0.00 | 1.44 | 509 | 2 nearest neighbours |
| Ex3e | 0.42 | 0.00 | -0.34 | 1636 | standard |
| Ex4 | 0.97 | 0.00 | 1.94 | 1134 | standard |
| Ex5a | 0.92 | 0.00 | 1.40 | 1500 | standard |
| Ex5b | 0.50 | 0.00 | -0.08 | 645 | without replacement |
| Ex6a | 0.97 | 0.00 | 1.68 | 366 | 2 nearest neighbours |
| Ex6b | 0.93 | 0.00 | 1.87 | 330 | 2 nearest neighbours |
| Ex8 | 0.94 | 0.00 | 1.28 | 499 | 2 nearest neighbours |
| In1a | 0.92 | 0.00 | 1.43 | 426 | 2 nearest neighbours |
| In1b | 0.97 | 0.00 | 1.95 | 370 | 2 nearest neighbours |
| In1c | 0.93 | 0.00 | 2.04 | 299 | 2 nearest neighbours |
| In2 | 0.91 | 0.00 | 1.52 | 388 | 2 nearest neighbours |
| In3 | 0.82 | 0.00 | 1.30 | 306 | 2 nearest neighbours |
| In4a | 0.91 | 0.00 | 1.13 | 349 | 2 nearest neighbours |
| In4b | 0.95 | 0.00 | 2.24 | 476 | 2 nearest neighbours |
| In6a | 0.97 | 0.00 | 1.62 | 332 | 2 nearest neighbours |
| In6b | 0.49 | 0.00 | 0.02 | 630 | without replacement |
| In7 | 0.91 | 0.00 | 1.28 | 372 | 2 nearest neighbours |
| In8 | 0.90 | 0.00 | 1.42 | 326 | 2 nearest neighbours |
| End | 0.56 | 0.00 | -0.13 | 245 | standard |
| Ast | 0.95 | 0.00 | 1.77 | 543 | standard |
| Oli | 0.84 | 0.00 | 0.74 | 704 | standard |
| OPC | 0.87 | 0.00 | 0.76 | 200 | 2 nearest neighbours |
| Mic | 0.55 | 0.00 | -0.03 | 300 | standard |

**Table S7 Adult Cell Enrichment Null Model Statistics:** We assessed adult cell specific enrichment of the genes on PLS1 (26). We used a median gene rank approach to analyse an enrichment of PLS1 for genes associated with specific cell clusters. We did so matching for gene length. Specifically, we resampled genes from the Allen Human Brain Atlas (AHBA) until we found a subset of genes that were not significantly different in length ($P < 0.05$) from the empirical gene list. Here, we show the results from the gene length matching algorithm, including the*P-,s- and Z-value* for the test of difference in gene length between the empirical gene list and the subset of resampled genes, the number of genes *N Genes*, to resample from, as well as which *approach* the algorithm converged on (see Supplementary Text *"Matching for Gene Length"* for details.)

| p | s | z | n | approach |
|---|---|---|---|---|
| 0.75 | 0.00 | 0.36 | 287 | standard |

**Table S8 MDD Enrichment Null Model Statistics:** We used a median gene rank approach to analyse an enrichment of PLS1 for genes associated with MDD risk genes (27). We did so matching for gene length. Specifically, we resampled genes from the Allen Human Brain Atlas (AHBA) until we found a subset of genes that were not significantly different in length ($P < 0.05$) from the empirical gene list. Here, we show the results from the gene length matching algorithm, including the*P-,s- and Z-value* for the test of difference in gene length between the empirical gene list and the subset of resampled genes, the number of genes *N Genes*, to resample from, as well as which *approach* the algorithm converged on (see Supplementary Text *"Matching for Gene Length"* for details.)

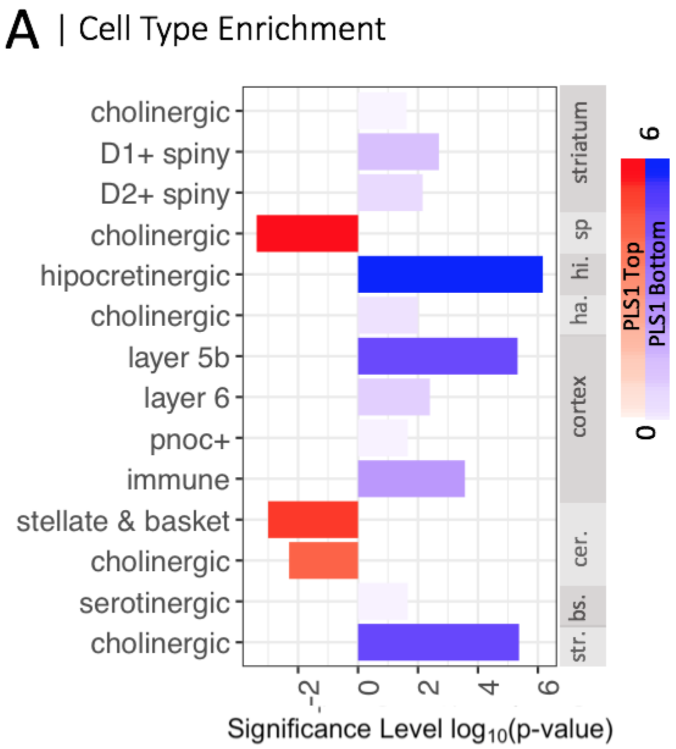

**Fig. S18: Cell type enrichment:** We uploaded the list of significantly positive and negative ($P_{FDR} < 0.05$) genes respectively to the CSEA tool's *specific expression analysis across cell types* function (23) and found $\Delta MI$ was associated with two separate transcriptomic signatures of gene expression. The bottom (Z<-2.58) of PLS1 was enriched for cholinergic and serotonergic cells across brain structures, as well as immune cells in the cortex, while the top (Z>2.58) was more sparsely associated with cholinergic and stellate and basket cells in the cerebellum and cholinergic cells in the spinal cord.

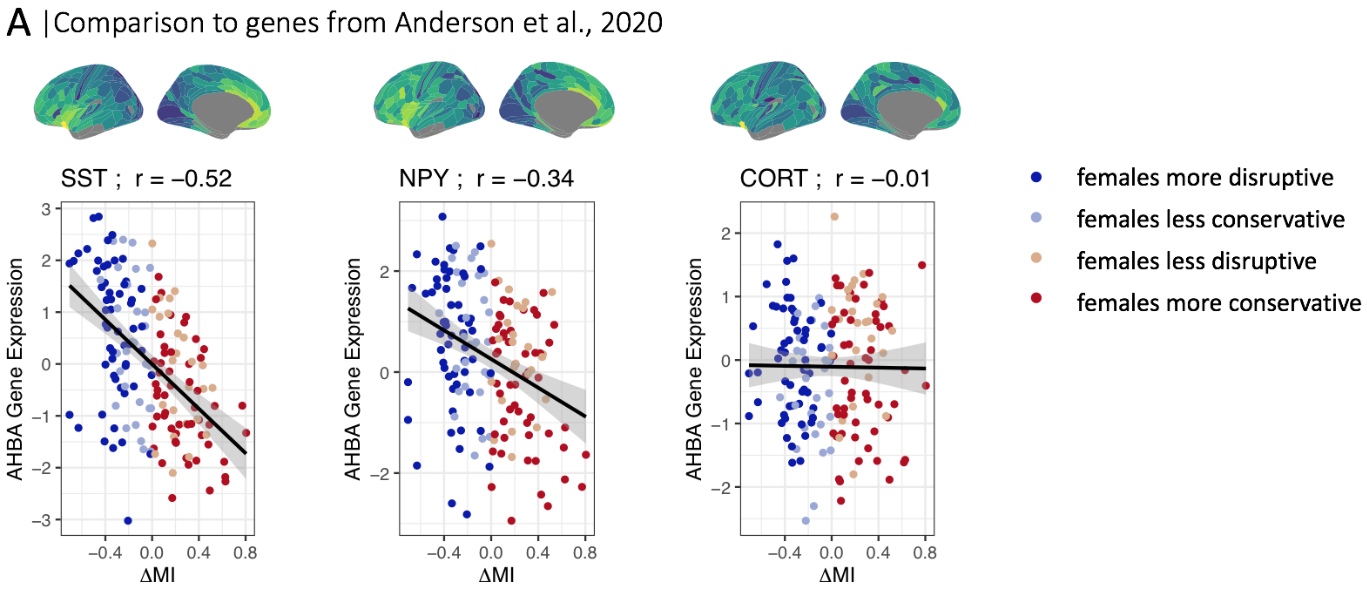

**Fig. S19: Correlation of PLS1 with Anderson et al., 2020 genes:** We correlated $\Delta MI$ with the Allen Human Brain Atlas gene expression data of three genes found to be related to major depression in (28): (A) SST, (B) NPY, and (C) CORT.

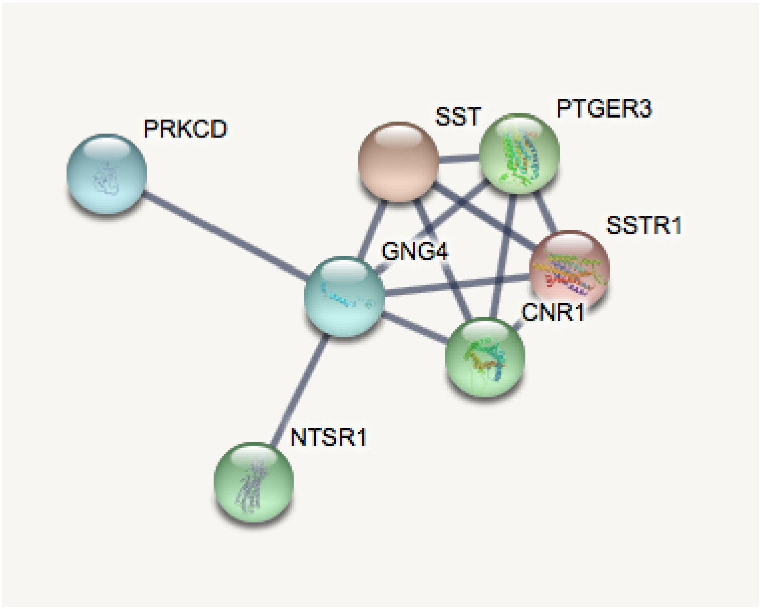

**Fig. S20: Protein-protein interaction (PPI) network:** We used STRING (29) to analyse protein-protein interactions in the 100 lowest ranking genes of PLS1. Here, we display the PPI at a confidence level of 0.9. We further varied the number of genes entered (100-250) and found the core network was stable to this variation.

#### Sensitivity Analysis

*Motion-Matched Sample*

*Sample Overview* We constructed a motion-matched subsample of the NSPN dataset by removing subjects with particularly high and low FD values from the original sample, until no significant difference was observed between males and females. If a subject was included in the sample, all of their follow-up scans were included, too. The final sample consisted of 314 subjects (156 females), 124 of which were scanned once, 89 twice and 4 three times (cf. Supplementary Table S9).

| Sex | # Scans | # Scanned | | | At Baseline | | | |
|---|---|---|---|---|---|---|---|---|
| | | 1 | 2 | 3 | $\mu$ Age | $\sigma$ Age | $\mu$ FD | $\sigma$ FD |
| female | 156 | 67 | 43 | 1 | 18.9 | 3.0 | 0.12 | 0.02 |
| male | 158 | 57 | 46 | 3 | 18.9 | 2.8 | 0.12 | 0.03 |

**Table S9 NSPN Motion-Matched Sample Overview:** We found a subset of subjects such that there was no significant sex difference in framewise displacement (FD; $P > 0.05$). We resampled subjects withing agebins and kept all scans from each single subject together, such that the original structure of the dataset was preserved.

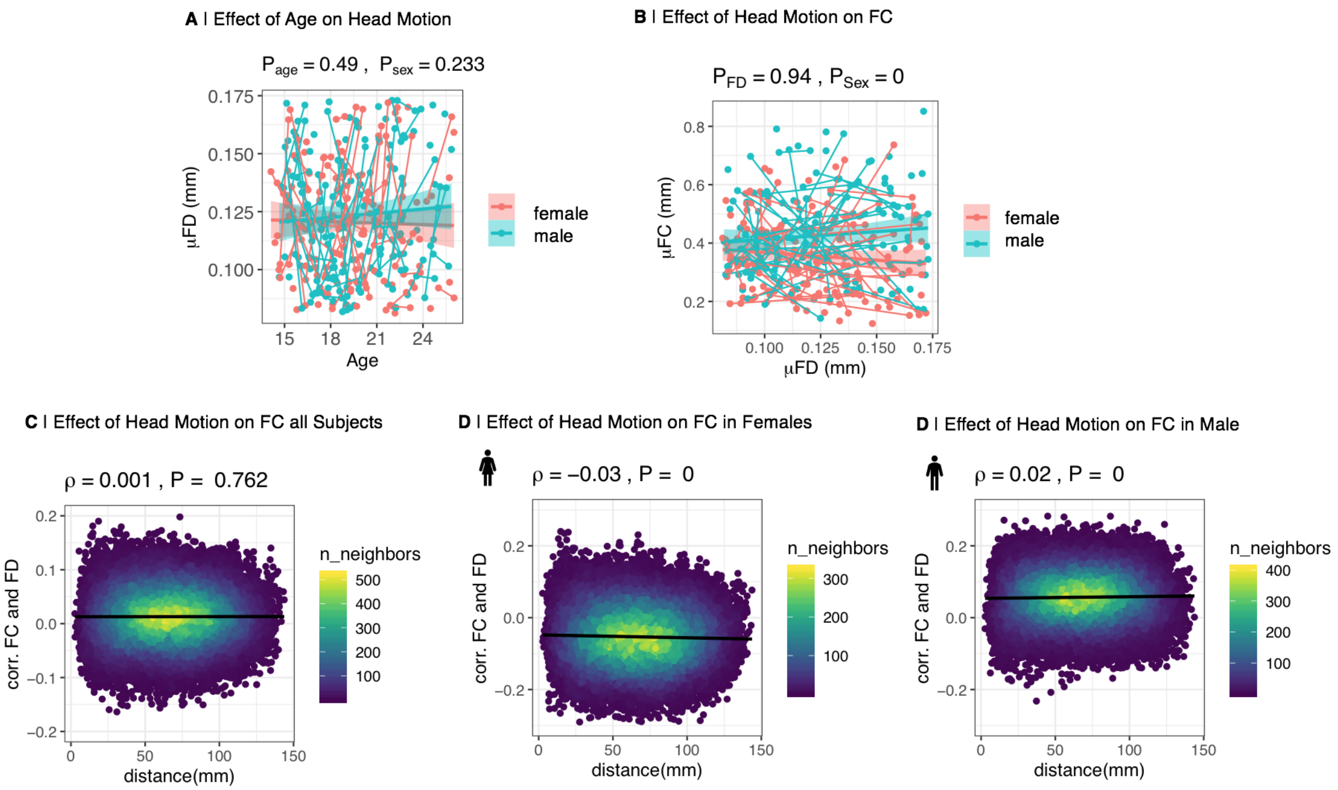

**Fig. S21: Effect of head motion (FD) on functional connectivity (FC) in the motion-matched sample:** The motion-matched sample is a subsample of the full data set, in which we removed the dependence of FC on motion in our sample by regressing FD from each edge; the residuals constitute participant-specific FD-corrected FC, with intercepts retained to maintain the relative importance of edges across the group as well as the interpretability of FC values. (A) In this subsample, average head motion, quantified as mean frame-wise displacement (FD), did not change with age ($P_{age} = 0.49$). And there was no effect of sex on FD ($P_{sex} = 0.23$). (B) The effect of participants' motion (across participants) on global FC was not significant ($P_{FD} = 0.94$). (C) There was no effect of distance on the correlation between FC and motion ($\rho = 0.001, P = 0.76$), and the average edge-wise correlation between FC and motion was almost zero (intercept = 0.01). (D) However, since our motion correction was performed across all subjects in the full sample, we still observed weak, but significant effects of distance on the correlation of FC and FD for females ($\rho = -0.03, P < 10^{-10}$) (D) and males ($\rho = 0.02, P < 10^{-10}$) (E) separately, and the average edge-wise correlation between FC and motion was non-zero ($intercept_{females} = -0.050.02, intercept_{males} = 0.05$).

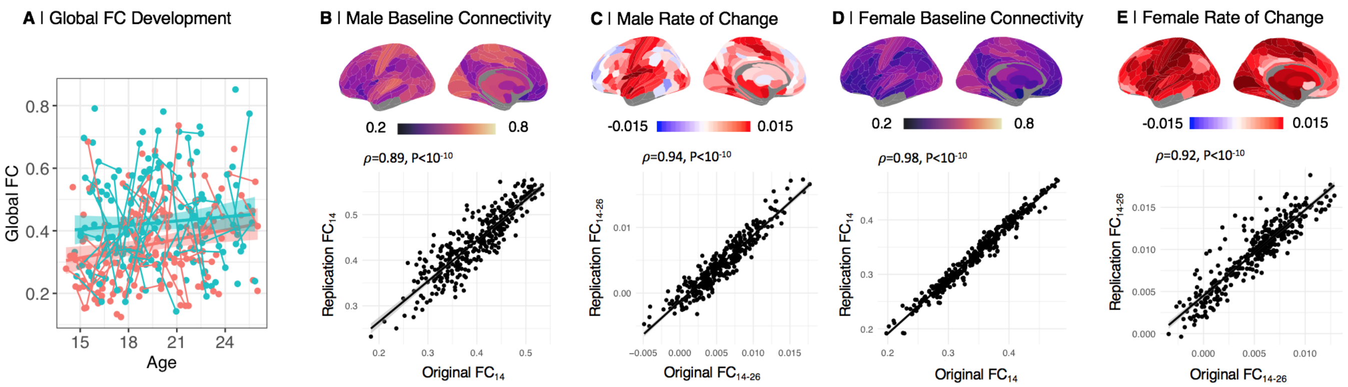

**Fig. S22: Replication of key elements of Fig. 1 in motion-matched sample:** Sex and age effects on functional connectivity (FC) were modeled using linear mixed effects models on different spatial scales. (A) Global FC increased with age ($t(94) = 2.48, P < 0.05$) and was higher in males ($t(215) = 3.84, P < 0.001$). (B)-(E) baseline connectivity at age 14 ($FC_{14}$) and adolescent rate of change ($FC_{14-26}$) in males and females were qualitatively and quantitatively highly consistent with our main results.

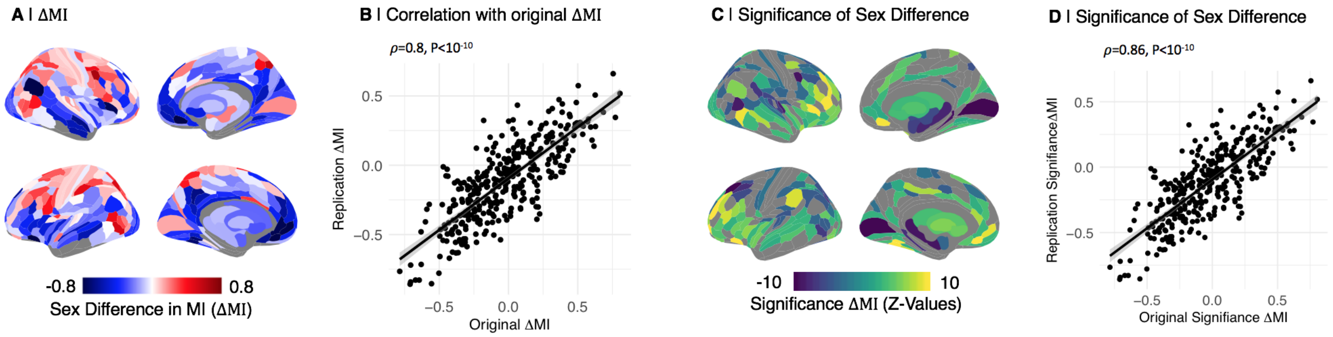

**Fig. S23: Replication of key elements of Fig. 2 in motion-matched sample:** (A) Sex difference in maturational index in the motion-matched sample ($\Delta MI$). (B) The original and replication $\Delta MI$ map were significantly correlated ($\rho = 0.8$, p < 2.2e-16) (C) The sex difference in MI was significant in 229 ROIs ($\alpha = 0.01$; -2.57 < z < 2.57). (D) The significance map of sex differences in maturational index was significantly correlated with the original map ($\rho = 0.86$, p < 2.2e-16).

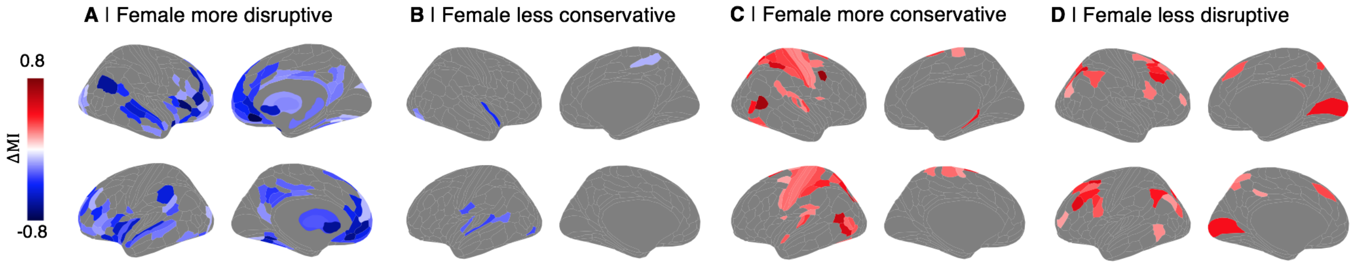

**Fig. S24: Trends in the sex difference in maturational development ($\Delta MI$) in the motion-matched sample:** Trends in $\Delta MI$, thresholded for regions with significant sex differences ($P_{FDR} < 0.05$).

*Global Signal Regression Sample*

*Sample Overview* We re-preprocessed all data with an alternative pipeline. The first pre-processing steps were the same as in the original sample. (cf. Methods) After ME-ICA pre-processing, however, we performed global signal regression (GSR). The global signal was estimated as the average time series of all cortical voxels. We regressed this time series from each region. From here, we proceeded with wavelet filtering using using brainwaver v. 1.6 and all following steps as in the original sample..

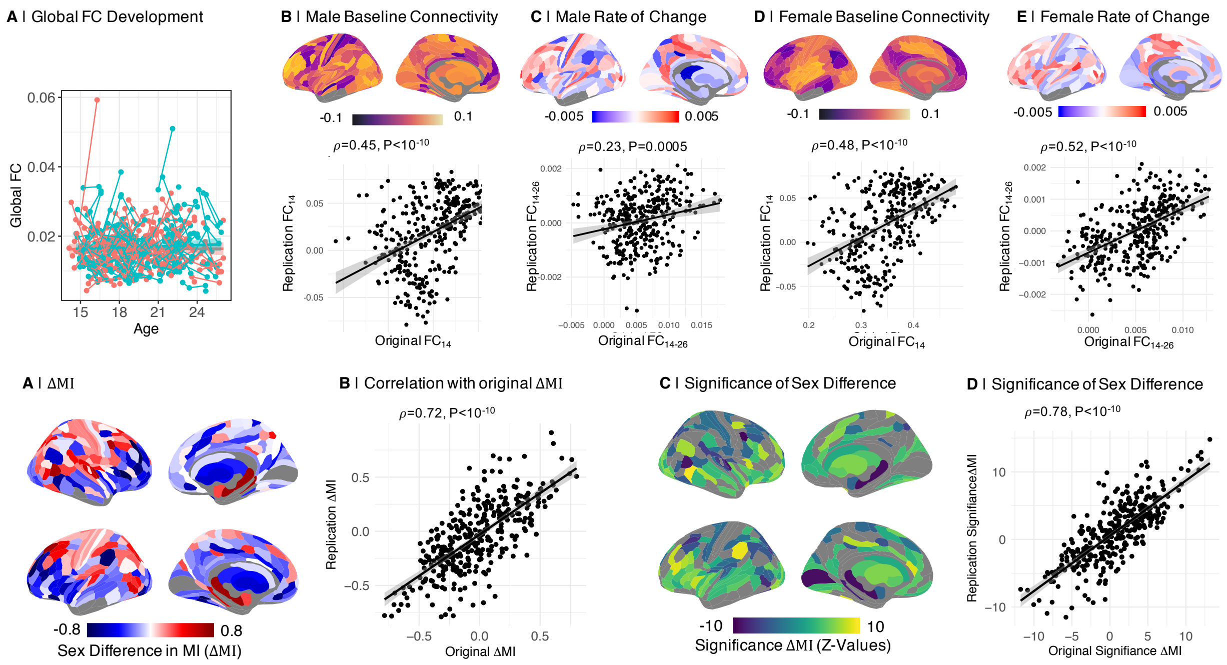

**Fig. S25: Replication of key elements of Fig. 1 in GSR sample:** Sex and age effects on functional connectivity (FC) were modeled using linear mixed effects models on different spatial scales. (A) We did not find significant effects of sex ($t(296) = 0.82, P = 0.41$) or age ($t(219) = 0.3, P = 0.77$) on global FC in the GSR sample. (B)-(E) The baseline connectivity at age 14 ($FC_{14}$) and adolescent rate of change ($FC_{14-26}$) in males and females in the GSR sample were quantitatively and qualitatively similar to the original maps.

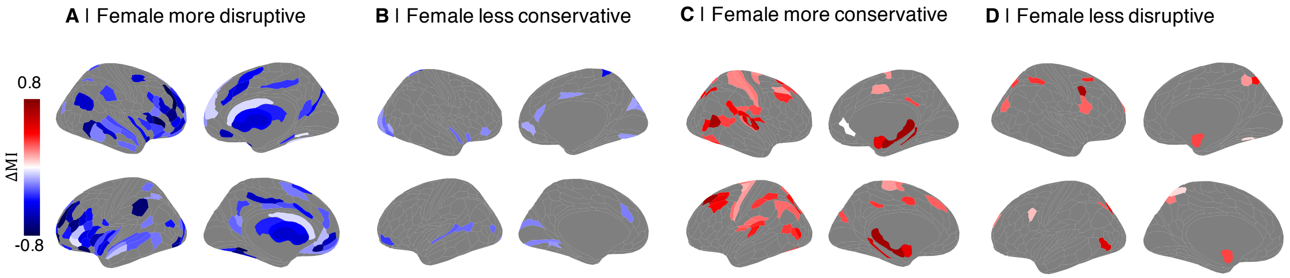

**Fig. S26: Replication of key elements of Fig. 2 in GSR sample:** (A) Sex difference in maturational index in the GSR sample ($\Delta MI$). (B) The original and replication $\Delta MI$ map were significantly correlated ($\rho = 0.71$, p < 2.2e-16) (C) The sex difference in MI was significant in 216 ROIs ($\alpha = 0.01$; -2.57 < z < 2.57). (D) The significance map of sex differences in maturational index was significantly correlated with the original map ($\rho = 0.76$, p < 2.2e-16).

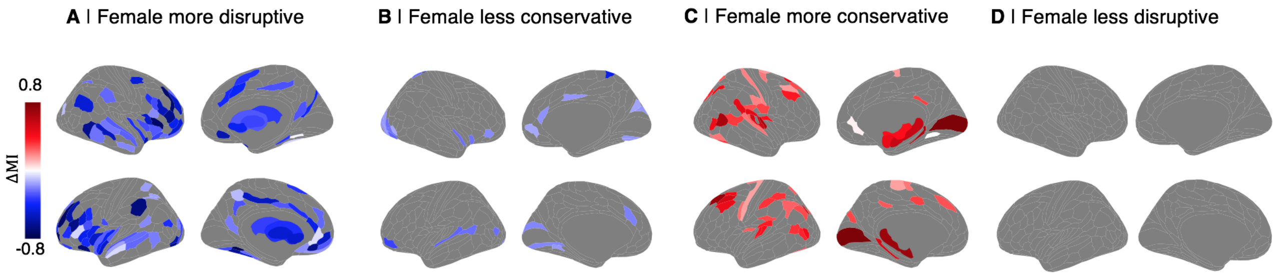

**Fig. S27: Trends in the sex difference in maturational development ($\Delta MI$) in the GSR sample:** Trends in $\Delta MI$, thresholded for regions with significant sex differences ($P_{FDR} < 0.05$).

*FD regression by sex sample*

*Sample Overview* We found that after our preprocessing pipeline including edgewise FD regression across the whole sample, there was a sex dependence of FC on motion. Therefore, we constructed a sample where we performed the edgewise FD regression per sex group instead.

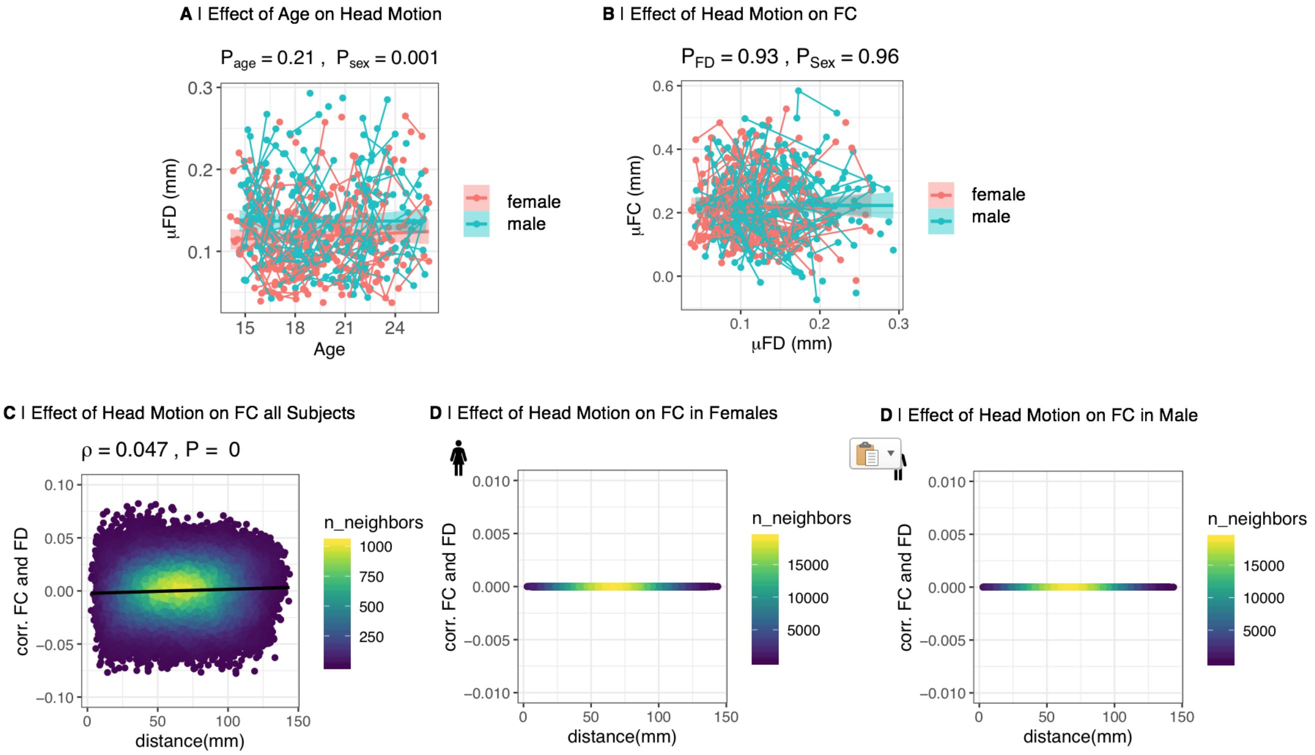

**Fig. S28: Effect of head motion (FD) on functional connectivity (FC) in the FD regression by sex sample :** To remove the sex difference in the dependence of FC on motion, mean FD was regressed from each edge for males and females separately; the residuals constitute participant-specific FD-corrected FC, with intercepts retained to maintain the relative importance of edges across the group as well as the interpretability of FC values. (A) Following this correction, subjects' average head motion, quantified as mean frame-wise displacement (FD), did not change with age ($P_{age} = 0.21, t(220) = 1.25$). However, there was a weak, but significant effect of sex on FD ($P_{sex} < 0.01, t(296) = 3.266$). (B) Mean participant motion was not related to mean FC across participants ($P_{FD} = 0.93, t(221) = 0.09$) and there is no effect of sex on the relationship ($P_{Sex} = 0.93, t(296) = -0.05$). (C) The correlation between FC at each edge and participant motion shows a weak but significant relationship with the Euclidean distance spanned by edges ($\rho = 0.05, P < 0.001$). The average edge-wise correlation between FC and motion is very close to zero (intercept = -0.002). By definition, the correlation between FC and motion almost completely vanished at the level of individual edges for males and females separately (D) and (E).

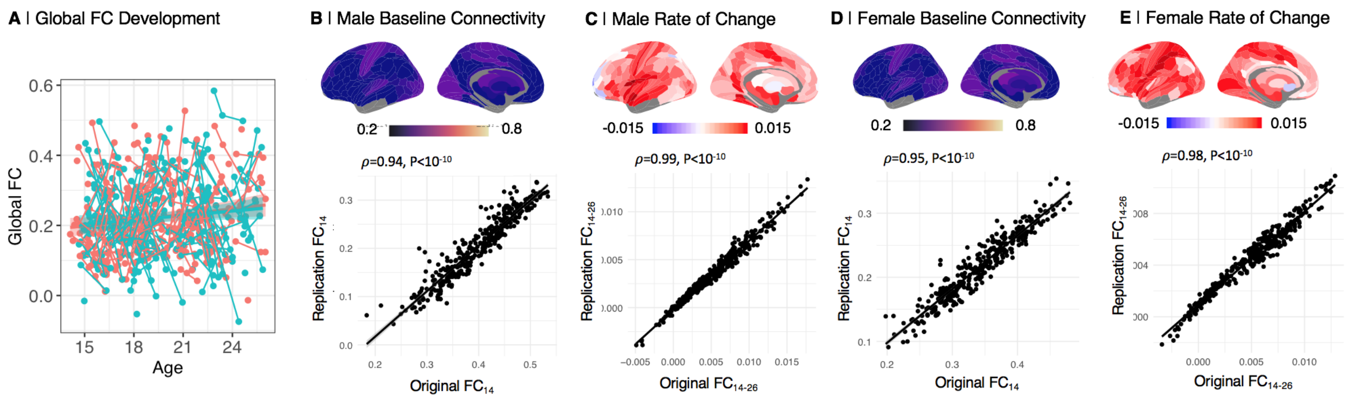

**Fig. S29: Replication of key elements of Fig. 1 in sample where FD was regressed per sex group:** Sex and age effects on functional connectivity (FC) were modeled using linear mixed effects models on different spatial scales. (A) Global FC increased with age ($t(219) = 2.94, P_{Age} < 0.05$). There was no significant effect of sex on global connectivity ($P_{Sex} = 0.73$). (B)-(E) baseline connectivity at age 14 ($FC_{14}$) and adolescent rate of change ($FC_{14-26}$) in males and females were qualitatively and quantitatively highly consistent with our main results.

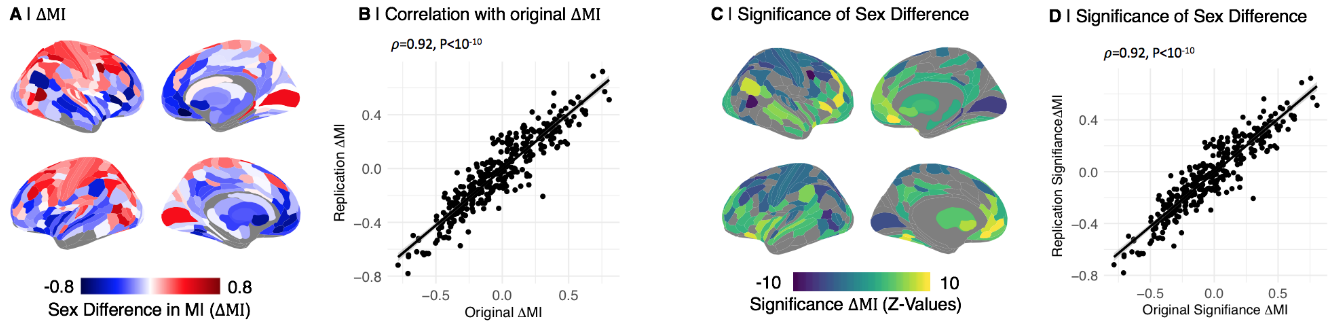

**Fig. S30: Replication of key elements of Fig. 2 in sample where FD was regressed per sex group:** (A) Sex difference in maturational index where FD was regressed per sex group: ($\Delta MI$). (B) The original and replication $\Delta MI$ map were significantly correlated ($\rho$ = 0.92, p < 2.2e-16) (C) The sex difference in MI was significant in 219 ROIs ($\alpha$ = 0.01; -2.57 < z < 2.57). (D) The significance map of sex differences in maturational index was significantly correlated with the original map ($\rho$ = 0.92, p < 2.2e-16).

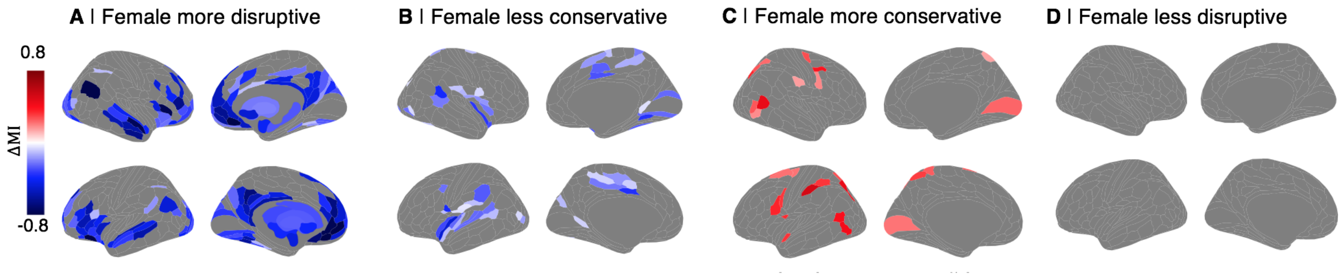

**Fig. S31: Trends in the sex difference in maturational development ($\Delta MI$) in the sample where FD was regressed by sex group:** Trends in $\Delta MI$, thresholded for regions with significant sex differences ($P_{FDR} < 0.05$).

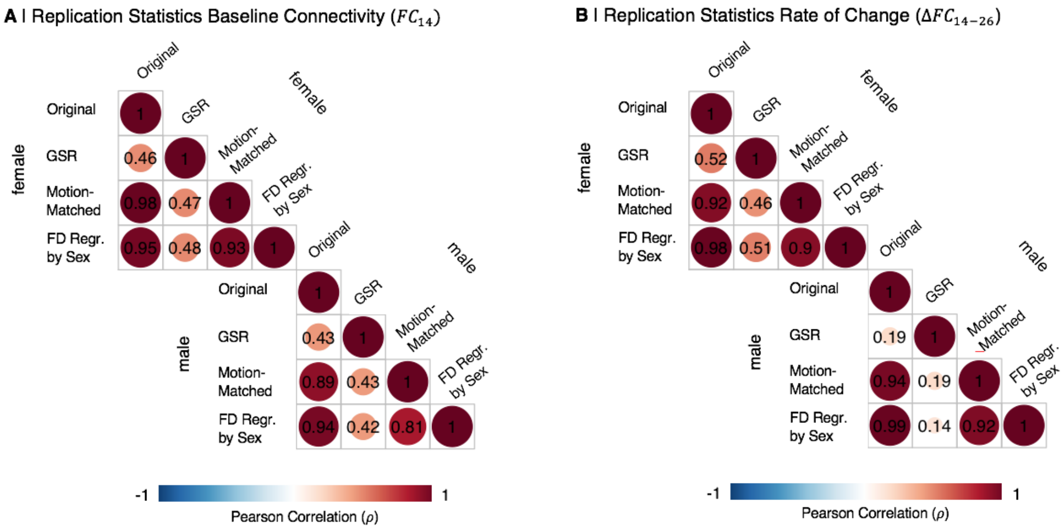

**Fig. S32: Summary of replication of baseline connectivity ($FC_{14}$) and adolescent rate of change ($FC_{14-26}$) statistics:** (A) Correlation of baseline connectivity ($FC_{14}$) across all regions calculated from original and replication data sets. (B) Correlation of adolescent rate of change ($FC_{14-26}$) across all regions calculated from original and replication data sets.

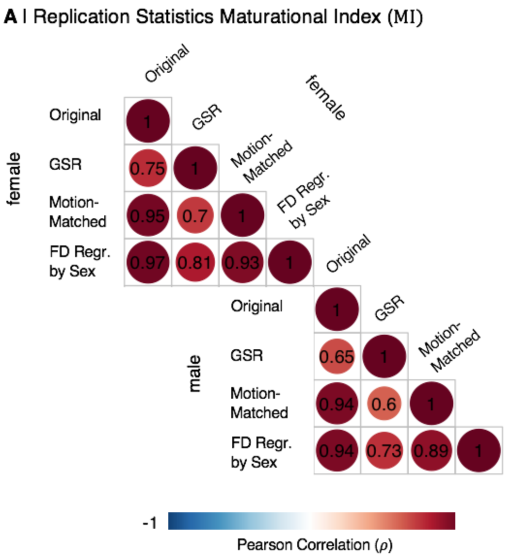

**Fig. S33: Summary of replication of maturational index ($MI$):** (A) Correlation of maturational index ($MI$) across all regions calculated from original and replication data sets. (B) Correlation of sex difference in maturational index ($\Delta MI$) across all regions calculated from original and replication data sets.

*Effects of Intracranial Volume*

To estimate the effects of intracranial volume (ICV) on our main results, we have included ICV in both the analysis of global functional connectivity, as well as in the maturational index calculation. Specifically, we included ICV as follows:

To estimate the effect of ICV on global functional connectivity, we added ICV to our model as a covariate:

$$FC_{global} \sim 1 + age * \beta_{age} + sex * \beta_{sex} + site * \beta_{site} + ICV * \beta_{ICV} + \gamma_{subject} * (1|subject) + \epsilon \quad (16)$$

Likewise, we also included ICV in our analysis of edge wise functional connectivity.

$$FC_{edge} \sim 1 + age * \beta_{age} + sex * \beta_{sex} + site * \beta_{site} + ICV * \beta_{ICV} + \gamma_{subject} * (1|subject) + \epsilon \quad (17)$$

We then carried the effect of ICV over into the calculation of baseline connectivity ($FC_{14}$), by adding $\beta_{ICV}$ (from Equation 17) coefficient multiplied by the mean ICV at age 14 for males and females respectively. Thus at each edge, we calculated baseline connectivity at age 14 as:

$$FC_{14} = 1 + 14 * \beta_{age} + sex * \beta_{sex} + 1/3 * \beta_{site_2} + 1/3 * \beta_{site_3} + \mu_{ICV_{14_{sex}}} * \beta_{ICV} + \gamma_{subject} * (1|subject) + \epsilon \quad (18)$$

where $\mu_{ICV_{14_{sex}}}$ is the mean ICV at age 14 for males and females respectively.

As before, the rate of change is calculated as:

$$FC_{14-26} = \beta_{age} \quad (19)$$

And the MI is the coefficient of the linear relationship between the ICV-corrected $FC_{14}$ and ICV-corrected $FC_{14-26}$.

We find that our key results replicate well in the sensitivity analysis including ICV. Specifically, original and ICV-corrected baseline connectivity are moderately correlated ($\rho_{female} = 0.49, P_{female} < 0.01; \rho_{male} = 0.43, P_{male} < 0.01$; Fig. S34A,C), whereas the adolescent rate of change is highly correlated ($\rho_{female} = 0.99, P_{female} < 0.01; \rho_{male} = 0.99, P_{male} < 0.01$; Fig. S34B,D).
The ICV-corrected controlled sex difference in maturational index ($\Delta MI$) is qualitatively and quantitatively similar to the original result from our main analysis ($\rho = 0.99, P < 0.01$; Fig. S35A,B).

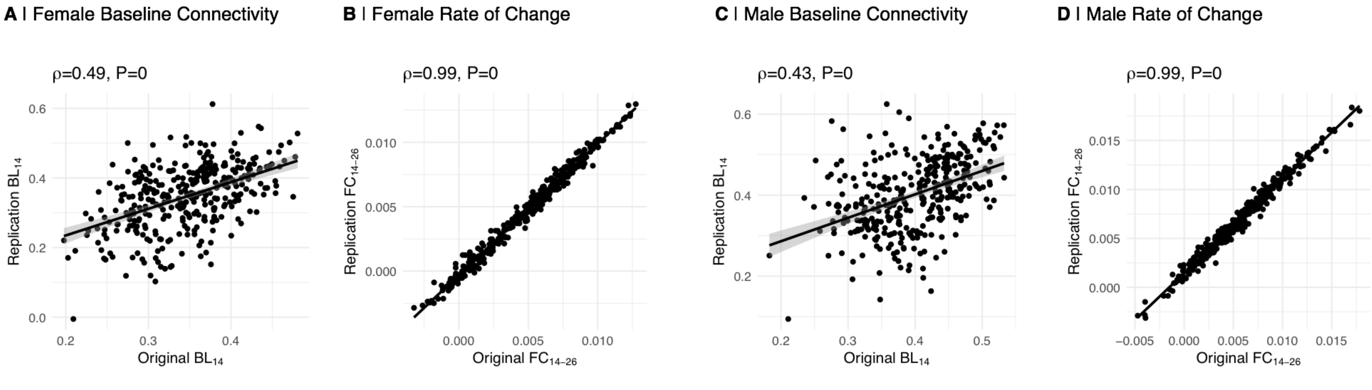

**Fig. S34: Baseline Connectivity and Rate of Change in ICV analysis:** We recalculated baseline baseline connectivity ($FC_{14}$) and adolescent rate of change ($FC_{14-26}$) controlling for mean intracranial volume (ICV). Here, we are depicting the correlation of replication and original $FC_{14}$ and $FC_{14-26}$ for females (A,B) and males (C,D) respectively.

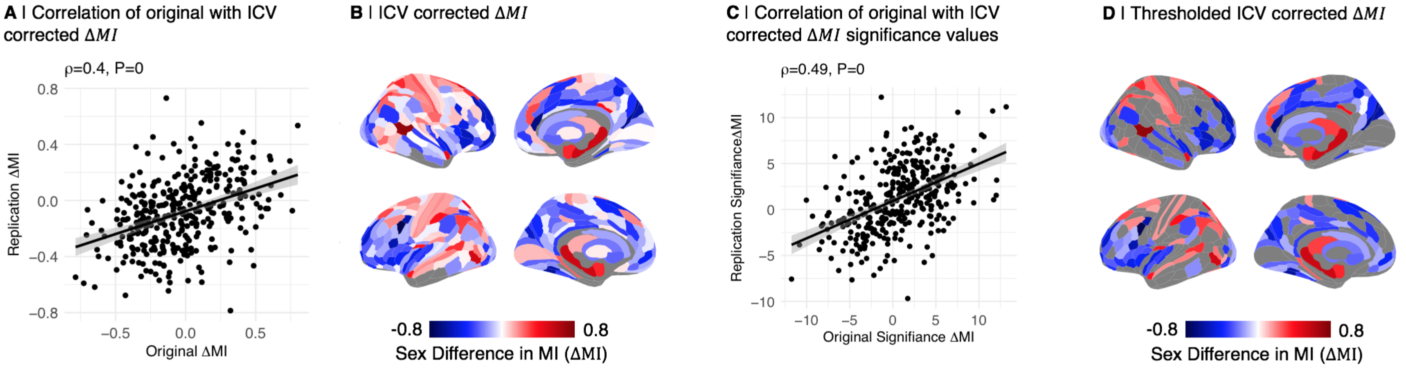

**Fig. S35: Maturational Index in ICV analysis:** We recalculated the sex difference in maturational index, $\Delta MI$, from baseline connectivity ($FC_{14}$) and adolescent rate of change ($FC_{14-26}$) that have been controlled for intracranial colume (ICV). (A) Correlation of original with ICV corrected $\Delta MI$. (B) ICV corrected $\Delta MI$. (C) Correlation of original and ICV corrected significance values (z-scores). (D) ICV corrected $\Delta MI$ thresholded for significance at p<0.05.

| Author Year | Cohort | N age | CS task | Whole-Brain | BL | Age | Inter. | Summary |
|---|---|---|---|---|---|---|---|---|
| Alarcon 2018 | in house | N=49 15-18 y. | CS, rsfMRI & flanker task | DMN, FPN | Yes | NA | NA | FC female >male in DMN and FPN during cognitive control in a self referential context<br>Mediated by higher co-rumination in females |
| Zhang 2018 | HCP | N=820 22-37 y. | CS rsfMRI | Whole-Brain | NA | NA | NA | PLS classification of gender (87%) based on whole brain FC<br>Most contributing ROIs in DMN, fronto-parietal and sensorimotor networks<br>Regressing out age had no effect |
| Mao 2017 | 1000 Funct. Con. | N=148 18-26 y. | CS, rsfMRI | Yes | Yes | NA | NA | 12 ROIs with gender differences in dynamic FC<br>Flex. female < male in amy., hipp. and parah.<br>Flex. male < female in TL, prec., MCG, SOC, IOG |
| Peterson 2017 | 1000 Func. Conn. | N=250 18-49 y. | CS, rsfMRI | VTA, SNc to whole-brain | Yes | in men | Yes | FC VTA/SNc to left posterior orbital gyrus male >women<br>Age effect in FC VTA/SNc to cortical and cerebellar ROIs in men only |
| Zhang 2016 | HCP | N=494 22-36 y. | CS rsfMRI | Whole-Brain | Yes | Yes | Yes | Edgeswise FC male>female in 1025 ROIs<br>Sex differences mostly in frontal, parietal and temporal lobes<br>Age effects in 29 ROIs<br>Interaction did not survive; greater decline in FC with age in females |
| Alarcon 2015 | Dataset | N=122 10-16 y. | CS, rsfMRI | amy. /w whole-brain | Yes. | Yes | Yes | increased FC amygdala and mPFC<br>SFA. and parieto-occipital ROIs FC decrease with age female > male<br>BLA and parieto-occipital ROIs FC decrease with age female > male |
| Scheinost 2015 | Cohort | N=103 18-65 y. | CS task | Whole-Brain | Yes | Yes | Yes | Used intrinsic connectivity distribution, ICD<br>Age by sex interaction in several networks, including DMN, FPN, visual, auditory<br>Age effect in several ROIs including putamen, caudate, amygdala, orbito frontal cortex (all pos.) and default mode (neg.)<br>Sex effect female > male in subcortex and limbic areas<br>Sex effect male > female in sensorimotor areas |
| Satterthw. 2014 | PNDC | N=674 9-22 y. | CS rsfMRI | Yes | Yes | NA | NA | Between module connectivity males >females<br>Within-module connectivity females >males<br>SVM classification of sex based on FC 71% accuracy |

| Author Year | Cohort | N age | CS task | Whole-Brain | BL | Age | Inter. | Summary |
|---|---|---|---|---|---|---|---|---|
| Filippi 2013 | Dataset | N=100 20-29 y. | CS rsFMRI | Network rsfMRI | Yes | NA | NA | Marginal sex effect in sensory networks<br>FC parietal-occipital ROIs male > female<br>FC frontal, temporal ROIs and cerebellum female > male<br>FNC cognitive, sensory male > female<br>FNC attention, right working memory female > male |
| Casanova 2012 | 1000 Func. Conn. | N=148 21 y. | CS rsfMRI | Whole-Brain | Yes | NA | NA | Lasso (62.3%) and Random Forest (65.4%) to classify sex<br>female discriminative edges: cingulate gyrus, left frontal lope, basal ganglia, thalamus, right cerebellum<br>male discriminative edges: cingulate gyrus, sensory motor cortex, cerebellum, left frontal lobe |
| Tian 2011 | in-house | N=86 18-25 y. | CS rsfMRI | within-hemi-sphere | Yes | NA | NA | Right hemisphere's clustering coefficient male >female<br>Left hemisphere's clustering coefficient female >male |
| Biswal 2010 | 1000 Func. Conn. | N=1,414 $\mu < 60$ y. | CS rsfMRI | Whole-Brain | Yes | Yes | NA | FC female >male in frontal ROIs<br>FC male >female in occipital and parietal ROIs<br>Delta Age positive in association cortex |
| Kong 2010 | Cohort | N=100 ?? | CS task | PAG to whole-brain | Yes | NA | NA | FC PAG to mid cingulate cortex female >male<br>FC PAG to left medial orbital prefrontal cortex, uncus, right insula und prefrontal cortex male >female |
| Weiss.-Fogel 2010 | in-house | N=49 21-50 y. | CS rsfMRI | ECN, SN, DMN | No | NA | NA | No within-network differences in ECN, SN, DMN |
| Liu 2009 | in-house | N=300 mu 22 y. | CS rsfMRI | Whole-Brain | Yes | NA | NA | Men showed stronger intrinsic laterality (left and right) |
| Kilpatrick 2005 | in-house | N=72 ?? | CS rsfMRI | amy. to whole | Yes | NA | NA | Right amy. FC male > female<br>Left amy. FC female > male |

**Table S10:** Reviewed papers

#### NSPN Consortium Author List

**Principal investigators:**
Edward Bullmore (CI from 01/01/2017)
Raymond Dolan
Ian Goodyer (CI until 01/01/2017)
Peter Fonagy
Peter Jones

**NSPN (funded) staff:**
Michael Moutoussis
Tobias Hauser
Sharon Neufeld
Rafael Romero-García
Michelle St Clair
Petra Vértes
Kirstie Whitaker
Becky Inkster
Gita Prabhu
Cinly Ooi
Umar Toseeb
Barry Widmer
Junaid Bhatti
Laura Villis
Ayesha Alrumaithi
Sarah Birt
Aislinn Bowler
Kalia Cleridou
Hina Dadabhoy
Emma Davies
Ashlyn Firkins
Sian Granville
Elizabeth Harding
Alexandra Hopkins
Daniel Isaacs
Janchai King
Danae Kokorikou
Christina Maurice
Cleo McIntosh
Jessica Memarzia
Harriet Mills
Ciara O'Donnell
Sara Pantaleone
Jenny Scott

**Affiliated scientists:**
Pasco Fearon
John Suckling
Anne-Laura van Harmelen
Rogier Kievit

#### NIMA Consortium Author List

**Cambridge**
Edward T. Bullmore (MD, PI, EC)[1,2,11]
Junaid Bhatti[1]
Samuel J. Chamberlain[1,2]
Marta M. Correia[1,12]
Anna L. Crofts[1]
Amber Dickinson*
Andrew C. Foster*
Manfred G. Kitzbichler[1]
Clare Knight*
Mary-Ellen Lynall[1]
Christina Maurice[1]
Ciara O'Donnell[1]
Linda J. Pointon[1]
Peter St George Hyslop[1,13,14]
Lorinda Turner[31]
Petra Vertes[1]
Barry Widmer[1]
Guy B. Williams[1,14]

**Cardiff**
B. Paul Morgan (PI)[15]
Claire A. Leckey[15]
Angharad R. Morgan*
Caroline O'Hagan*
Samuel Touchard[15]

**Glasgow**
Jonathan Cavanagh (PI, EC)[3]
Catherine Deith*
Scott Farmer[16]
John McClean[16]
Alison McColl[3]
Andrew McPherson*
Paul Scouller*
Murray Sutherland[16]

**Independent advisor**
H.W.G.M. (Erik) Boddeke (EC)[17]

**GSK**
Jill C. Richardson (EC)[18]
Shahid Khan[11]
Phil Murphy[1]
Christine A. Parker[19]
Jai Patel[11]

**Janssen**
Declan Jones (EC)[6]
Peter de Boer[4]
John Kemp[4]
Wayne C. Drevets[6]
Jeffrey S. Nye (deceased)
Gayle Wittenberg[6]
John Isaac[6]
Anindya Bhattacharya[6]

Nick Carruthers[6]
Hartmuth Kolb[6]

**Kings College London**
Carmine M. Pariante (PI)[10]
Federico Turkheimer (PI)[20]
Gareth J. Barker[20]
Heidi Byrom[10]
Diana Cash[20]
Annamaria Cattaneo[10]
Antony Gee[20]
Caitlin Hastings[10]
Nicole Mariani[10]
Anna McLaughlin[10]
Valeria Mondelli[10]
Maria Nettis[10]
Naghmeh Nikkheslat[10]
Karen Randall[20]
Hannah Sheridan*
Camilla Simmons[20]
Nisha Singh[20]
Victoria Van Loo*
Marta Vicente-Rodriguez[20]
Tobias C. Wood[20]
Courtney Worrell*
Zuzanna Zajkowska*

**Lundbeck**
Niels Plath (EC)[21]
Jan Egebjerg[21]
Hans Eriksson[21]
Francois Gastambide[21]
Karen Husted Adams[21]
Ross Jeggo*
Christian Thomsen[21]
Jan Torleif Pederson[21]
Brian Campbell*
Thomas Möller*
Bob Nelson*
Stevin Zorn*

**University of Texas (sub-contracted to Lundbeck)**
Jason O'Connor[22]

**Oxford**
Mary Jane Attenburrow (PI)[7,23]
Alison Baird
Jithen Benjamin[23]
Stuart Clare[25]
Philip Cowen[7]
I-Shu (Dante) Huang[24]
Samuel Hurley*
Helen Jones[23]
Simon Lovestone[7]
(AD, PI, EC) Francisca Mada*
Alejo Nevado-Holgado[7]
Akintayo Oladejo*
Elena Ribe7

Katy Smith[23]
Anviti Vyas*

**Pfizer**
Zoe Hughes*
Rita Balice-Gordon*
James Duerr*
Justin R. Piro*
Jonathan Sporn*

**Southampton**
V. Hugh Perry (PI)[27]
Madeleine Cleal*
Gemma Fryatt[27]
Diego Gomez-Nicola[27]
Renzo Mancuso[32]
Richard Reynolds[27]

**Sussex**
Neil A. Harrison (PI, EC)[28]
Mara Cercignani[28]
Charlotte L. Clarke[28]
Elizabeth Hoskins*
Charmaine Kohn*
Rosemary Murray*
Lauren Wilcock[29]
Dominika Wlazly[30]

**University of Toronto (sub-contracted to Cambridge)**
Howard Mount[13]

MD = Mood disorder workpackages lead
AD = Alzheimer's disease workpackages lead
PI = Principal Investigator
EC = Executive committee member
1 Department of Psychiatry, School of Clinical Medicine, University of Cambridge, CB2 0SZ, UK
2 Cambridgeshire and Peterborough NHS Foundation Trust, Cambridge, CB21 5EF, UK
3 Sackler Centre, Institute of Health & Wellbeing, University of Glasgow, Sir Graeme Davies Building , Glasgow, G12 8TA, UK
4 Neuroscience, janssen Research & Development, Janssen Pharmaceutica NV, Turnhoutseweg 30, B-2340, Beerse, Belgium
5 The Maurice Wohl Clinical Neuroscience Institute, Cutcombe Road, London, SE5 9RT, UK
6 Neuroscience, janssen Research & Development, LLC, Titusville, NJ, 08560, USA 7 Department of Psychiatry, University of Oxford, Warneford Hospital, Oxford, OX3 7JX, UK
8 Brighton & Sussex Medical School, University of Sussex, Brighton, BN1 9RR, UK 9 Sussex Partnership NHS Foundation Trust, Swandean, BN13 3EP, UK
10 Kings College London, Institute of Psychiatry, Psychology and Neuroscience, Department of Psychological Medicine, London, SE5 9RT, UK
11 Immuno-Psychiatry, Immuno-Inflammation Therapeutic Area Unit, GlaxoSmithKline R&D, Stevenage SG1 2NY, UK
12 MRC Cognition and Brain Sciences Unit, 15 Chaucer Road, Cambridge CB2 7EF, UK
13 Tanz Centre for Research in Neurodegenerative Diseases, 60 Leonard Avenue, Toronto, ON M5T 2S8 Canada
14 Department of Clinical Neurosciences, University of Cambridge, CB2 0SZ, UK 15 Cardiff University, Cardiff CF10 3AT, UK
16 NHS Greater Glasgow and Clyde, 1055 Great Western Rd, Glasgow G12 0XH, UK 17 University of Groningen, 9712 CP Groningen, Netherlands
18 Neurosciences Virtual PoC DPU, GlaxoSmithKline R&D, Stevenage SG1 2NY, UK
19 Experimental Medicine Imaging, GlaxoSmithKline R&D, Stevenage SG1 2NY, UK
20 King's College London, Department of Neuroimaging Sciences, Institute of Psychiatry, Psychology & Neuroscience, De Crespigny Park, London SE5 8AF, UK
21 H. Lundbeck A/S Ottiliavej 9, 2500, Valby, Denmark
22 University of Texas Health Science Center at San Antonio, 7703 Floyd Curl Dr, San Antonio, TX 78229, USA

23 NIHR Oxford cognitive health Clinical Research Facility, Warneford Hospital, Oxford, OX3 7JX, UK
24 The Kennedy Institute of Rheumatology, Roosevelt Dr, Oxford OX3 7FY, UK
25 Oxford Centre for Functional MRI of the Brain, John Radcliffe Hospital, Oxford OX3 9DU, UK
26 Pfizer, Inc, 1 Portland Street, Cambridge MA, USA
27 Centre for Biological Sciences, University of Southampton, Southampton, UK
28 Clinical Imaging Sciences Centre (CISC), University of Sussex, Brighton, BN1 9RR, UK
29 Sussex Partnership NHS Foundation Trust, Nevill Avenue, Hove BN3 7HZ, UK
30 Brighton & Sussex University Hospitals NHS Trust, Brighton BN2 5BE, UK
31 Department of Medicine, School of Clinical Medicine, University of Cambridge, CB2 0SZ, UK
32 VIB-KU Leuven Center for Brain & Disease Research, Campus Gasthuisberg, Herestraat 49, bus 602, 3000 Leuven, Belgium

*Former consortium members

### References

**1** Nikolaus Weiskopf, John Suckling, Guy Williams, Marta M Correia, Becky Inkster, Roger Tait, Cinly Ooi, Edward T Bullmore, and Antoine Lutti. Quantitative multi-parameter mapping of r1, pd(*), mt, and r2(*) at 3t: a multi-center validation. *Frontiers in Neuroscience*, 7:95, 2013.

**2** Markus Barth, Jürgen R Reichenbach, Ramesh Venkatesan, and Ewald Moserand Mark E Haacke. High-resolution, multiple gradient-echo functional MRI at 1.5 T. *Magnetic Resonance Imaging*, 17(3):321–329, 1999.

**3** Bruce Fischl, Martin I. Sereno, and Anders M. Dale. Cortical surface-based analysis: Ii: Inflation, flattening, and a surface-based coordinate system. *NeuroImage*, 9(2):195–207, 1999.

**4** Kirstie J Whitaker, Petra E Vértes, Rafael Romero-Garcia, František Váša, Michael Moutoussis, Gita Prabhu, Nikolaus Weiskopf, Martina F Callaghan, Konrad Wagstyl, Timothy Rittman, Roger Tait, Cinly Ooi, John Suckling, Becky Inkster, Peter Fonagy, Raymond J Dolan, Peter B Jones, Ian M Goodyer, the NSPN NSPN Consortium, and Edward T Bullmore. Adolescence is associated with genomically patterned consolidation of the hubs of the human brain connectome. *Proceedings of the National Academy of Sciences of the United States of America*, 113(32):9105–10, 2016.

**5** František Váša, Jakob Seidlitz, Rafael Romero-Garcia, Kirstie J Whitaker, Gideon Rosenthal, Petra E Vértes, Maxwell Shinn, Aaron Alexander-Bloch, Peter Fonagy, Raymond J Dolan, Peter B Jones, Ian M Goodyer, Olaf Sporns, and Edward T Bullmore. Adolescent Tuning of Association Cortex in Human Structural Brain Networks. *Cerebral Cortex*, 28(1):281–294, 2018.

**6** Ziad S Saad, Daniel R Glen, Gang Chen, Michael S Beauchamp, Rutvik Desai, and Robert W Cox. A new method for improving functional-to-structural mri alignment using local pearson correlation. *NeuroImage*, 44(3):839–848, 2009.

**7** Prantik Kundu, Souheil J Inati, Jennifer W Evans, Wen-Ming Luh, and Peter A Bandettini. Differentiating BOLD and non-BOLD signals in fMRI time series using multi-echo EPI. *NeuroImage*, 60(3):1759–1770, 2012.

**8** Prantik Kundu, Noah D Brenowitz, Valerie Voon, Yulia Worbe, Petra E Vértes, Souheil J Inati, Ziad S Saad, Peter A Bandettini, and Edward T Bullmore. Integrated strategy for improving functional connectivity mapping using multiecho fMRI. *Proceedings of the National Academy of Sciences of the United States of America*, 110(40):16187–92, 2013.

**9** Edward T Bullmore, Jalal Fadili, Voichita Maxim, Levent Şendur, Brandon Whitcher, John Suckling, Michael Brammer, and Michael Breakspear. Wavelets and functional magnetic resonance imaging of the human brain. *NeuroImage*, 23:S234–S249, 2004.

**10** Matthew F Glasser, Timothy S Coalson, Emma C Robinson, Carl D Hacker, John Harwell, Essa Yacoub, Kamil Ugurbil, Jesper Andersson, Christian F Beckmann, Mark Jenkinson, Stephen M Smith, and David C Van Essen. A multi-modal parcellation of human cerebral cortex. *Nature*, 536(7615):171–178, 2016.

**11** P A Filipek, C Richelme, D N Kennedy, and V S Caviness. The Young Adult Human Brain: An MRI-based Morphometric Analysis. *Cerebral Cortex*, 4(4):344–360, 1994.

**12** František Váša, Rafael Romero-Garcia, Manfred G Kitzbichler, Jakob Seidlitz, Kirstie J Whitaker, Matilde M Vaghi, Prantik Kundu, Ameera X Patel, Peter Fonagy, Raymond J Dolan, Peter B Jones, Ian M Goodyer, Petra E Vértes, and Edward T Bullmore. Conservative and disruptive modes of adolescent change in human brain functional connectivity. *Proceedings of the National Academy of Sciences*, 117(6):3248–3253, 2020.

**13** Philip Reiss, Lei Huang, Yin-Hsiu Chen, Lan Huo, Thaddeus Tarpey, and Maarten Mennes. Massively parallel nonparametric regression, with an application to developmental brain mapping. *Journal of computational and graphical statistics*, 23:232–248, 02 2014.

**14** Raymond Paternoster, Robert Brame, Paul Mazerolle, and Alex Piquero. Using the Correct Statistical Test for the Equality of Regression Coefficients. *Criminology*, 36(4):859–866, 1998.

**15** B T Yeo, F M Krienen, J Sepulcre, M R Sabuncu, D Lashkari, M Hollinshead, J L Roffman, J W Smoller, L Zöllei, J R Polimeni, B Fischl, H Liu, and R L Buckner. The organization of the human cerebral cortex estimated by intrinsic functional connectivity. *Journal of neurophysiology*, 106(3):1125–1165, 2011.

**16** Benedikt A Poser, Maarten J Versluis, Johannes M Hoogduin, and David G Norris. Bold contrast sensitivity enhancement and artifact reduction with multiecho epi: Parallel-acquired inhomogeneity-desensitized fmri. *Magnetic Resonance in Medicine*, 55(6):1227–1235, 2006.

**17** Robert W Cox. AFNI: Software for Analysis and Visualization of Functional Magnetic Resonance Neuroimages. *Computers and Biomedical Research*, 29(3):162–173, 1996.

**18** Petra E Vértes, Timothy Rittman, Kirstie J Whitaker, Rafael Romero-García, František Váša, Manfred G Kitzbichler, Konrad Wagstyl, Peter Fonagy, Raymond J Dolan, Peter B Jones, Ian M Goodyer, and Edward T Bullmore. Gene transcription profiles associated with inter-modular hubs and connection distance in human functional magnetic resonance imaging networks. *Philosophical Transactions of the Royal Society B: Biological Sciences*, 371(1705):20150362, 2016.

**19** Sarah E Morgan, Jakob Seidlitz, Kirstie J Whitaker, Rafael Romero-Garcia, Nicholas E Clifton, Cristina Scarpazza, Therese van Amelsvoort, Machteld Marcelis, Jim van Os, Gary Donohoe, David Mothersill, Aiden Corvin, Andrew Pocklington, Armin Raznahan, Philip McGuire, Petra E Vértes, and Edward T Bullmore. Cortical patterning of abnormal morphometric similarity in psychosis is associated with brain expression of schizophrenia-related genes. *Proceedings of the National Academy of Sciences of the United States of America*, 116(19):9604–9609, 2019.

**20** A Fornito A Arnatkeviciute, B D Fulcher.

**21** M Hawrylycz, E Lein, A Guillozet-Bongaarts, and et al. An anatomically comprehensive atlas of the adult human brain transcriptome. *Nature*, page 391–399, 2012.

**22** Jakob Seidlitz, Ajay Nadig, Siyuan Liu, Richard A I Bethlehem, Petra E Vértes, Sarah E Morgan, František Váša, Rafael Romero-Garcia, François M Lalonde, Liv S Clasen, Jonathan D Blumenthal, Casey Paquola, Boris Bernhardt, Konrad Wagstyl, Damon Polioudakis, Luis de la Torre-Ubieta, Daniel H Geschwind, Joan C Han, Nancy R Lee, Declan G Murphy, Edward T Bullmore, and Armin Raznahan. Transcriptomic and cellular decoding of regional brain vulnerability to neurogenetic disorders. *Nature Communications*, 11(3358), 2020.

**23** Xiaoxiao Xu, Alan B Wells, David R O'Brien, Arye Nehorai, and Joseph D Dougherty. Cell type-specific expression analysis to identify putative cellular mechanisms for neurogenetic disorders. *Journal of Neuroscience*, 34(4):1420–1431, 2014.

**24** Jeremy A Miller, Song-Lin Ding, Susan M Sunkin, Kimberly A Smith, Lydia Ng, Aaron Szafer, Amanda Ebbert, Zackery L Riley, Joshua J Royall, Kaylynn Aiona, James M Arnold, Crissa Bennet, Darren Bertagnolli, Krissy Brouner, Stephanie Butler, Shiella Caldejon, Anita Carey, Christine Cuhaciyan, Rachel A Dalley, Nick Dee, Tim A Dolbeare, Benjamin A C Facer, David Feng, Tim P Fliss, Garrett Gee, Jeff Goldy, Lindsey Gourley, Benjamin W Gregor, Guangyu Gu, Robert E Howard, Jayson M Jochim, Chihchau L Kuan, Christopher Lau, Chang-Kyu Lee, Felix Lee, Tracy A Lemon, Phil Lesnar, Bergen McMurray, Naveed Mastan, Nerick Mosqueda, Theresa Naluai-Cecchini, Nhan-Kiet Ngo, Julie Nyhus, Aaron Oldre, Eric Olson, Jody Parente, Patrick D Parker, Sheana E Parry, Allison Stevens, Mihovil Pletikos, Melissa Reding, Kate Roll, David Sandman, Melaine Sarreal, Sheila Shapouri, Nadiya V Shapovalova, Elaine H Shen, Nathan Sjoquist, Clifford R Slaughterbeck, Michael Smith, Andy J Sodt, Derric Williams, Lilla Zöllei, Bruce Fischl, Mark B Gerstein, Daniel H Geschwind, Ian A Glass, Michael J Hawrylycz, Robert F Hevner, Hao Huang, Allan R Jones, James A Knowles, Pat Levitt, John W Phillips, Nenad Šestan, Paul Wohnoutka, Chinh Dang, Amy Bernard, John G Hohmann, and Ed S Lein. Transcriptional landscape of the prenatal human brain. *Nature*, page 199–206, 2014.

**25** Damon Polioudakis, Luis [de la Torre-Ubieta], Justin Langerman, Andrew G Elkins, Xu Shi, Jason L Stein, Celine K Vuong, Susanne Nichterwitz, Melinda Gevorgian, Carli K Opland, Daning Lu, William Connell, Elizabeth K Ruzzo, Jennifer K Lowe, Tarik Hadzic, Flora I Hinz, Shan Sabri, William E. Lowry, Mark B Gerstein, Kathrin Plath, and Daniel H Geschwind. A single-cell transcriptomic atlas of human neocortical development during mid-gestation. *Neuron*, 103(5):785 – 801.e8, 2019.

**26** N Lake, S Chen, B Sos, J Fan, G E Kaeser, Y C Yung, T E Duong, D Gao, J Chun, P V Kharchenko, and K Zhang. Integrative single-cell analysis of transcriptional and epigenetic states in the human adult brain. *Nature Biotechnology*, 36:70–80, 2018.

**27** M Li, G Santpere, Y Imamura Kawasawa, O V Evgrafov, F O Gulden, S Pochareddy, S M Sunkin, Z Li, Y Shin, Y Zhu, A Sousa, D M Werling, R R Kitchenand H J Kang, M Pletikos, J Choi, S Muchnik, X Xu, D Wang, B Lorente-Galdos, . . . , and N Sestan. Integrative functional genomic analysis of human brain development and neuropsychiatric risks. *Science*, 362, 2018.

**28** Kevin M Anderson, Meghan A Collins, Ru Kong, Kacey Fang, Jingwei Li, Tong He, Adam M Chekroud, Thomas B T Yeo, and Avram J Holmes. Convergent molecular, cellular, and neural signatures of major depressive disorder. *bioRxiv*, 2020.

**29** D Szklarczyk, AL Gable, D Lyon, A Junge, S Wyder, J Huerta-Cepas, M Simonovic, NT Doncheva, JH Morris, P Bork, LJ Jensen, and CV Mering. String v11: protein-protein association networks with increased coverage, supporting functional discovery in genome-wide experimental datasets. *Nucleic Acids Res*, 28:3442–3444, 2019.
